# Automated Virtual Pathology Panels for ML-Assisted Interpretation of Mass Spectrometry Imaging

**DOI:** 10.64898/2026.07.26.740867

**Authors:** Jacob Gildenblat, Jens Pahnke

## Abstract

Mass spectrometry imaging (MSI) records rich molecular spectra at each pixel, but pathology-oriented interpretation benefits from complementary views of the same tissue, loosely analogous to the multiple stains used in histopathology. We present an expert-aligned framework for constructing multi-view MSI panels. *Soft Landmark Contrast Edges* (SoLaCE) extracts molecular boundaries directly from high-dimensional spectra. Because standard structure-preservation metrics aligned poorly with rankings by pathologists, we combine luminance contrast and chromatic diversity with SpecEdge-Dice, a boundary-aware measure of agreement between visualization edges and SoLaCE boundaries. *Parametric MiCS+LMC* (pMiCS) uses a neural network trained on sub-sampled data to distill multiple MSI segmentations into a reusable spectral-to-RGB mapping, enabling rapid full-image inference, out-of-sample projection, and more consistent color semantics across aligned images. A concept-based interpretation procedure explains pMiCS outputs through sparse mixtures of spectral concepts, scoped as a model-output-equivalent summary rather than a molecular identification. We integrate these components into *Virtual Pathology Panels*, which use hyperparameter optimization to select fast, high-performing or spatially complementary views. This framework supports future workflows that combine automated, AI/ML-supported, morphology-oriented tissue assessment and molecular analysis within a single MSI acquisition.

**Teaser:** Virtual pathology panels transform MSI spectra into complementary views for scalable, interpretable tissue analysis.

## Introduction

Mass spectrometry imaging (MSI) captures spatially resolved molecular information by measuring pixel-wise mass spectra. This high-dimensional structure is biologically rich (*1*) but difficult to inspect directly (*2*). In practice, RGB visualizations are used to summarize spectral variation and reveal tissue regions, boundaries, and pathology-relevant patterns.

Complete molecular annotation of every discriminating mass-to-charge (*m*/*z*) signal is not required to obtain a useful *exploratory* visualization, although molecular identification and orthogonal validation remain necessary for biological or clinical claims. Surgical pathology offers a partial analogy: diagnosis often begins with morphological patterns in hematoxylin and eosin (H&E) staining, which reports tissue and cellular morphology rather than a single defined molecular target, and is then refined with targeted assays. Immunohistochemistry, by contrast, is explicitly target-specific: each antibody visualizes a selected, known antigen. Our visualizations correspond to the first, exploratory step; they are intended to prioritize regions and signals for the molecular identification and validation that follow, not to replace them.

Analogously, MSI visualizations can first reveal reproducible spatial patterns, boundaries, and tissue states. Targeted molecular annotation can then be applied to diagnostically or biologically relevant regions and features to support interpretation and characterize biological behavior. In principle, a single MSI acquisition can provide morphological context and molecular information simultaneously, creating opportunities for more integrated tissue analysis.

For translational pathology research, these visualizations are not merely exploratory figures; they function as interpretive interfaces for reviewing molecular tissue structure. Pathologists need to move rapidly between whole-slide context and fine local detail, compare regions across samples, and iteratively test hypotheses against molecular contrast. MSI visualization methods must therefore be fast enough for interactive use, stable enough to preserve interpretable color meaning across images, and expressive enough to reveal subtle pathology-relevant structure that may be missed by sparse ion-image inspection. Because different visualizations may reveal different aspects of the high-dimensional data in different tissue regions, a single visualization may not be sufficient.

Current MSI visualization workflows face persistent challenges. First, modern MSI acquisitions can contain very large, high-resolution images, making repeated optimization expensive. At 5 *µ*m resolution in particular, nonparametric full-image nonlinear visualization can become impractical in routine workflows. Second, most modern MSI visualizations remain nonparametric and therefore cannot directly project unseen data. This limitation causes color semantics to drift across images, restricts reliable cross-sample comparison in translational settings, and complicates application across laboratories.

A related challenge is quality evaluation. Metrics commonly used in dimensionality reduction and previously applied to MSI, such as local or global structure preservation (*3, 4*, 5), are useful geometric diagnostics, but they do not take spatial organization into account and may not reflect what an expert pathologist considers visually informative in anatomical visualizations. For pathology-oriented review, coherent region boundaries and transitions may matter more than abstract manifold structure.

MSI analysis has moved well beyond single-ion images. Established workflows include unsupervised segmentation and clustering of spatially indexed spectra (*6, 7*, 8, 9, 10), multivariate and manifold visualization (*5, 11*, 12, 13), deep-learning representations (*14*), and community data and software ecosystems (*15*), reviewed in (*16, 17*, 18). Our contributions build on this body of work rather than replace it: SoLaCE provides a continuous molecular-boundary view complementary to region-labeling segmentation; pMiCS turns segmentation-style supervision into a fast, reusable parametric mapping; and the panels and concept interpretation add expert-guided view selection and molecular traceability to existing pipelines.

We address these challenges through five linked contributions.

### SoLaCE for molecular edge detection

First, we consider high-dimensional boundary detection in MSI images. Molecular boundary detection provides a pathology-oriented visualization in its own right and, as we show, a basis for visualization evaluation. Because MSI interpretation depends on transitions between tissue states, we introduce *Soft Landmark Contrast Edges* (SoLaCE), a high-dimensional edge detector designed to recover molecular boundaries directly from spectra. Magnitude-based edge methods may miss biochemically meaningful transitions when the signal is distributed across multiple correlated channels or when total intensity remains similar across regions. SoLaCE addresses this limitation by representing each pixel through soft similarities to a sparse set of landmark spectra, averaging these signatures locally, and measuring contrast across opposite sides of each pixel while suppressing within-region texture. The resulting edge maps highlight coherent molecular boundaries.

### Metrics for visualization evaluation

Second, we consider how to evaluate MSI visualizations for pathology-oriented review. We encode pathology-oriented preferences for high-contrast images that reveal anatomical boundaries using quantifiable metrics. For color variety, we use luminance RMS contrast and LAB chroma entropy to quantify light–dark separation and chromatic diversity over tissue pixels. For structure-aware evaluation, we introduce *SpecEdge-Dice*, which quantifies how well edges in RGB visualizations agree with edges extracted from high-dimensional MSI data. We compare these metrics against standard structure-preservation metrics and blinded ranking by expert pathologists.

### pMiCS for parametric MSI visualization

Third, we address how to create parametric MSI visualizations that are fast to generate, support out-of-sample projection, and align with pathology-oriented preferences. We propose *parametric MiCS+LMC* (pMiCS), in which MiCS+LMC (*19*) is used as a target embedding on sampled points and a lightweight neural network learns a reusable mapping from spectra to RGB. As described in the pMiCS subsection of Materials and Methods, the MiCS objective distills several segmentations of the MSI data into a single visualization. This builds on the common MSI practice of clustering-based segmentation of spatial MSI data. Instead of relying on a single clustering model that assigns each pixel to a discrete class, MiCS combines several segmentations into a continuous visualization. pMiCS enables rapid full-image inference, direct out-of-sample projection, and consistent color semantics across datasets when preprocessing and feature alignment are matched. In the blinded comparison benchmark, pMiCS and TOP3 formed the top-ranked pair; pMiCS also performed strongly on the quantitative metrics that correlated positively with the rankings.

### Concept-based interpretability

Fourth, we address how to connect a visually useful pMiCS rendering back to molecular spectra. The proposed interpretation procedure represents MSI spectra as nonnegative molecular concepts, then asks which sparse mixture of concepts is sufficient to reproduce the output of the frozen pMiCS model. This separates two related but distinct quantities: concepts that are abundant in the pixel and concepts that the visualization model uses to generate the displayed contrast. The full procedure is described in the concept-based interpretation subsection of Materials and Methods.

### Virtual Pathology Panels (VPP)

Finally, we address the need for MSI to support a coordinated panel of complementary views rather than only a single visualization. We refer to such automatically assembled collections as VPPs. Each VPP view is analogous to a non-IHC histochemical view: it is an unsupervised spectral phenotype map rather than a readout of a known molecular target. A panel therefore indicates *where* tissue differs across views and is hypothesis-generating rather than a validated molecular assignment. The framework integrates the preceding contributions by treating MSI visualization as a model-selection and panel-construction problem rather than assuming that one RGB rendering can capture all pathology-relevant structure. Candidate views are generated by sweeping model configurations, ranked using expert-aligned criteria, and selected for either overall quality or spatial complementarity; selected views can subsequently undergo sparse molecular concept interpretation.

### Long-term aim – AI/ML-assisted, pathologist-controlled tissue assessment

The framework computationally “stains” molecular tissue information by converting MSI spectra into standardized views selected using expert-aligned criteria. These views present molecular boundaries and spectral phenotypes for review while keeping the pathologist in control of interpretation and follow-up. Because pMiCS is parametric, a frozen model can repeatedly generate the same type of molecular view for appropriately matched tissue samples. With standardized preprocessing, feature alignment, and acquisition controls, a validated model could in principle be transferred between research or pathology laboratories. Model versioning, quality-control procedures, external validation, and prospective clinical studies would be required before regulatory qualification or clinical deployment.

## Results

### SoLaCE reveals molecular boundaries missed by magnitude-based edges

SoLaCE produces boundary maps that summarize high-dimensional MSI structure as a single pathology-oriented image. Rather than deriving edges from total ion current, a single ion image, or the magnitude of spatial differences, SoLaCE estimates boundary strength from changes in spectral composition. In brief, each pixel is represented by soft similarities to landmark spectra; locally averaged landmark signatures are then compared across opposite sides of each pixel to emphasize molecular transitions while suppressing within-region texture (the SoLaCE subsection of Materials and Methods).

Across different tissue MSI sections, SoLaCE recovered coherent anatomical and pathological boundaries directly from molecular information (Figure 2 and Supplementary Figure S1). The resulting maps delineate tissue interfaces and internal structures that are difficult to summarize with a single intensity image. Thus, SoLaCE can be read as a compact molecular boundary image, complementary to common diagnostic overview baselines such as total ion current (TIC): TIC primarily reports signal abundance, whereas SoLaCE reports where molecular composition changes across space.

**Figure 1:**
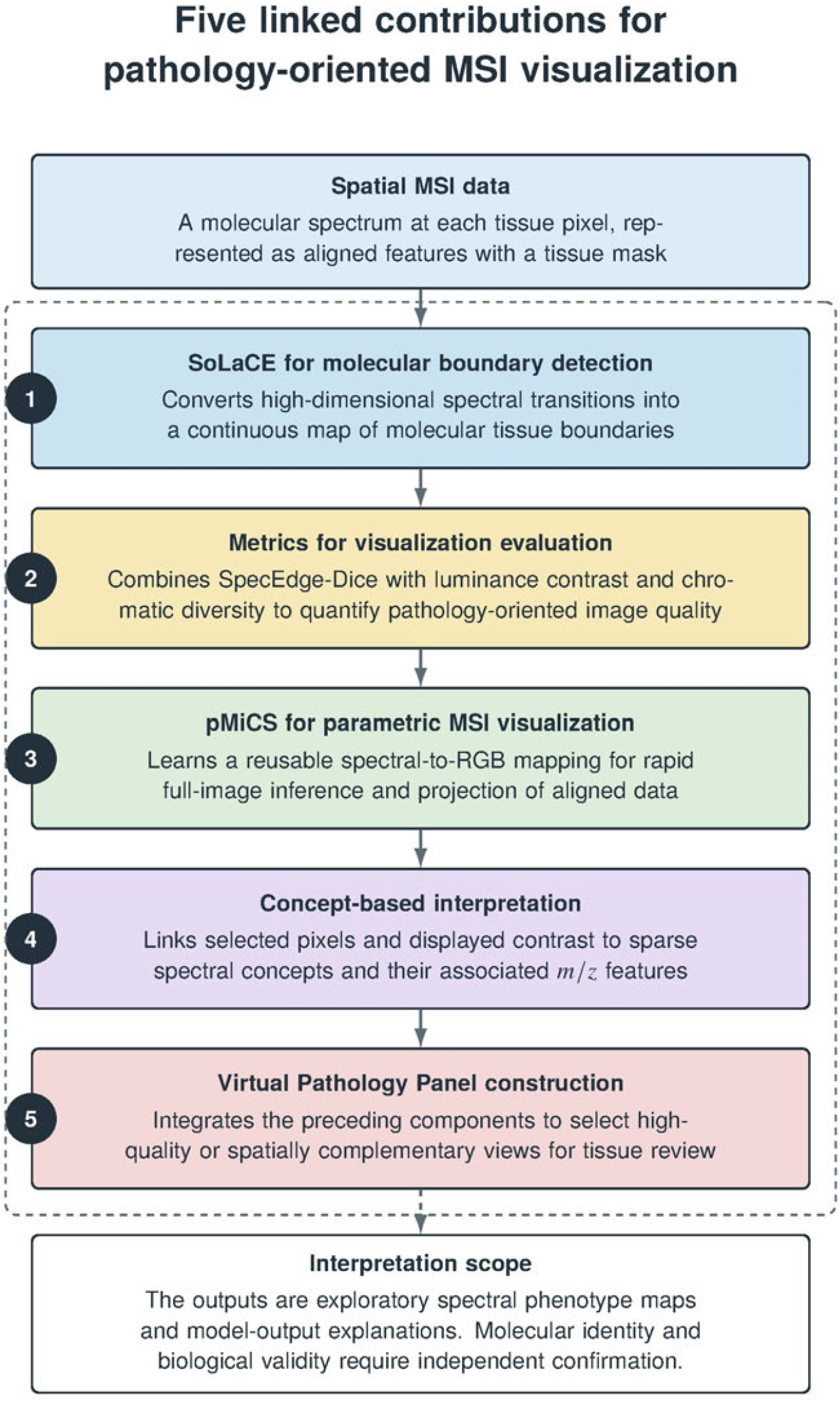
Contributions. Spatially indexed MSI data provide the common input. SoLaCE maps molecular boundaries; pathology-oriented metrics evaluate visualization quality; pMiCS learns a reusable spectral-to-RGB mapping; concept-based interpretation links displayed contrast to sparse spectral concepts and associated *m*/*z* features; and Virtual Pathology Panel construction selects high-quality or spatially complementary views. The outputs are exploratory spectral phenotype maps and model-output explanations; molecular identity and biological validity require independent confirmation.

**Figure 2:**
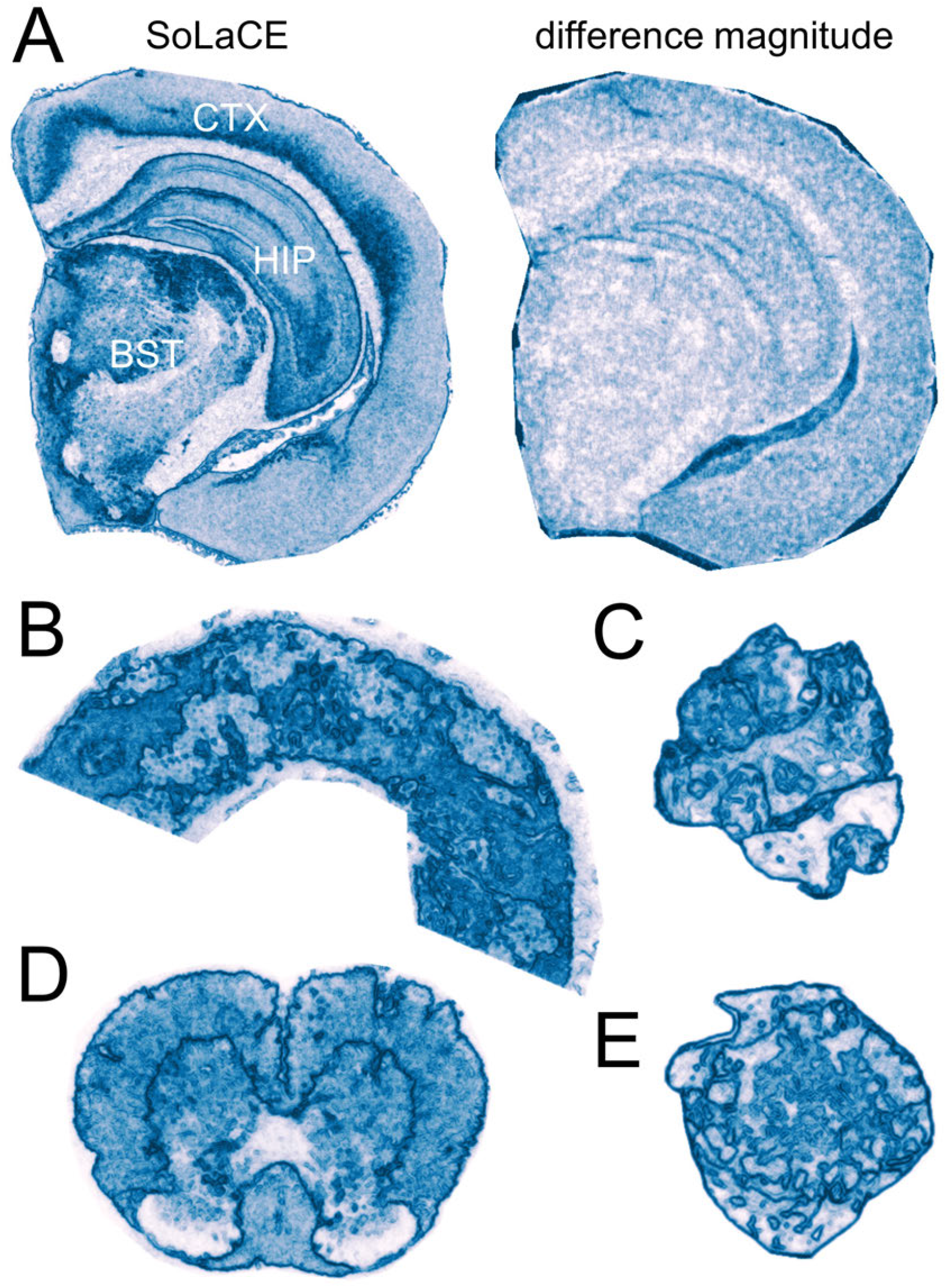
SoLaCE reveals molecular tissue boundaries across MSI sections. (**A**) Soft Landmark Contrast Edges (SoLaCE; left) and a difference-magnitude baseline (right) in a mouse brain hemisphere (5 *µ*m resolution; bregma −2.7 mm (*20*)). The baseline uses the maximum Euclidean distance to neighboring spectra; isolated responses obscure internal detail. SoLaCE compares locally averaged soft landmark signatures to reveal coherent transitions in the cortex (CTX), hippocampus (HIP), and brainstem (BST). (**B**–**E**) SoLaCE maps of a human kidney needle biopsy (**B**; 5 *µ*m resolution), colorectal cancer liver metastases (**C, E**; 20 *µ*m resolution), and mouse spinal cord (**D**; 5 *µ*m resolution). In the SoLaCE maps, darker blue denotes stronger local spectral boundaries, rank-normalized over tissue pixels within each image; off-tissue pixels are masked. Intensities are relative, not quantitatively comparable across sections, and do not encode tissue area, molecular abundance, or the direction of spectral change.

The advantage over a simple high-dimensional magnitude-edge baseline is illustrated in Figure 2A. The baseline assigns each pixel the maximum Euclidean distance between its spectrum and spectra in the local neighborhood. Although this is a direct spectral edge measure, it is dominated by isolated high-distance responses, whereas most within-tissue distances remain comparatively small. As a result, anatomical detail is suppressed. By comparing locally averaged landmark signatures instead, SoLaCE preserves spatially coherent tissue structure and remains sensitive to subtle molecular differences within related regions. In the brain example, these features include cortical structure and focal boundaries within the hippocampus and brainstem that are largely absent from the magnitude baseline.

To quantify alignment with expert-defined anatomy, we benchmarked SoLaCE against difference-magnitude and TIC-Sobel baselines, and against a spatially aware *k*-means segmentation modeled on Cardinal’s spatialKMeans (*9*) at fixed cluster counts (*k* = 8, 32, 128), using our polygon annotations from MSI-ATLAS (*1*) (Figure 3) (see Evaluation against expert anatomical boundaries in Materials and Methods). Because these annotations delineate anatomical regions rather than molecular ground-truth boundaries, the benchmark assessed enrichment at annotated borders relative to region interiors and local surroundings, together with boundary-ranking and density-matched detection performance. Against the difference-magnitude and TIC-Sobel baselines, SoLaCE achieved the highest values for all pooled metrics displayed and the highest F1 score at each evaluated edge density. It also outperformed the spatial *k*-means segmentation at every fixed *k*: SoLaCE gave the highest border enrichment (1.37 versus at most 1.24), average precision (0.79 versus 0.70), ROC-AUC (0.69 versus 0.60), and mean F1 (0.29 versus 0.24); the only measure on which a segmentation setting edged ahead was border-to-surrounding enrichment, where the densest clustering (*k* = 128) reached 0.985 against SoLaCE’s 0.977, as expected for a very dense binary edge map. Because SoLaCE uses a single fixed configuration whereas the segmentation baseline requires a choice of *k*, this is a tuning-free comparison against the clustering used in established MSI software. A one-at-a-time sensitivity analysis over wide ranges of SoLaCE’s own hyperparameters (Supplementary Figure S2) confirms that this fixed configuration is a mid-range choice rather than a tuned optimum: boundary performance varies smoothly and modestly with each setting, is robust to the landmark seed, and requires only that some spatial averaging is used (patch radius *≥* 1). SoLaCE’s relative performance varied across anatomical superclasses, indicating that expert anatomical borders and molecular transitions do not coincide uniformly across tissue types. All SoLaCE edge maps and their corresponding baseline maps were also reviewed qualitatively by pathologists. Across all reviewed images, the pathologists preferred SoLaCE to both baselines, judging its boundary maps more informative and anatomically coherent (Supplementary Figure S1).

**Figure 3:**
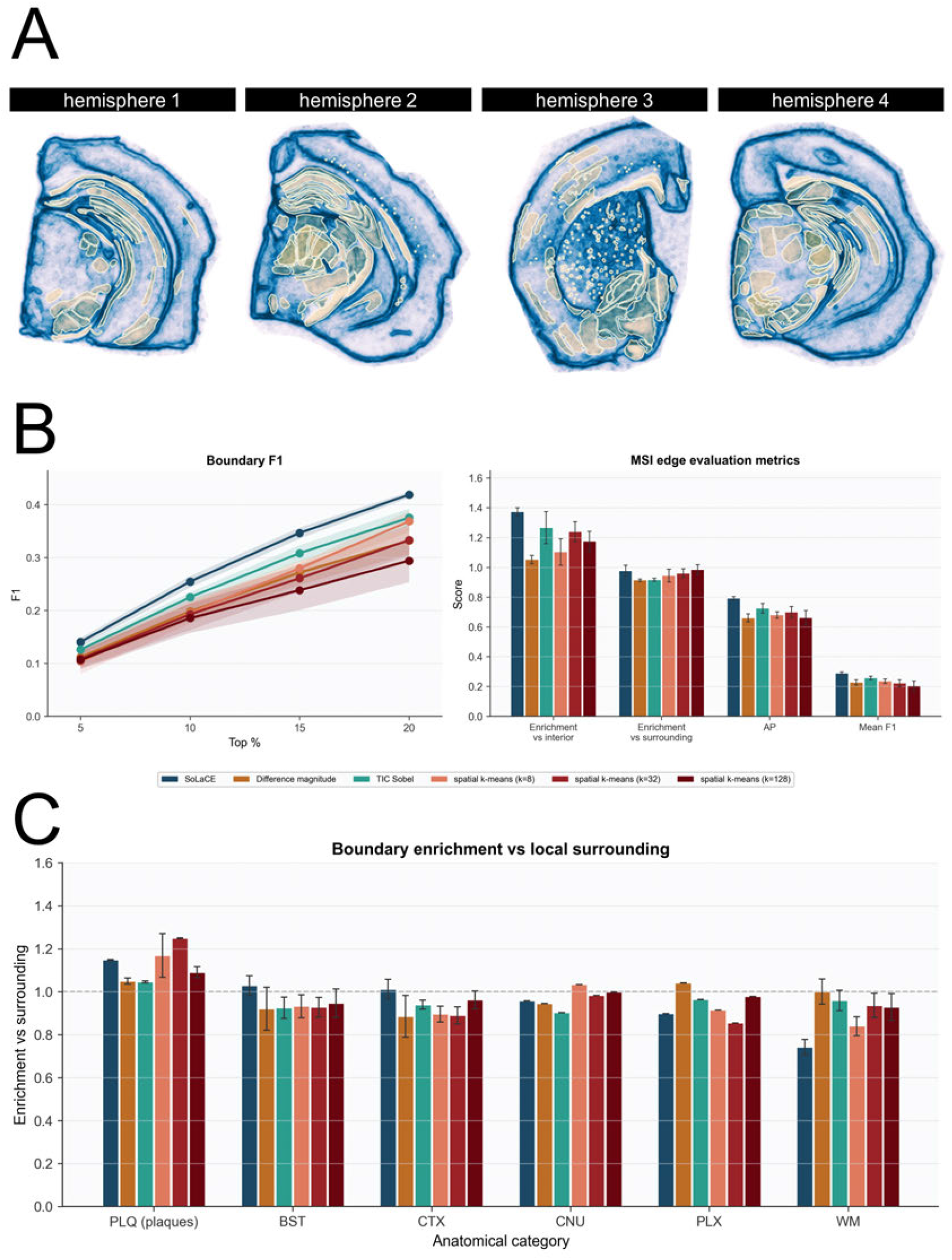
Evaluation of SoLaCE against expert anatomical boundaries. (**A**) SoLaCE maps with expert boundaries for four mouse brain hemispheres from the MSI-ATLAS study (*1*). (**B**) Boundary F1 at four edge densities (top 5–20%) and pooled metrics—enrichment versus region interior, enrichment versus local surroundings, average precision (AP), and mean F1—comparing SoLaCE with the difference-magnitude and TIC-Sobel baselines and spatial *k*-means segmentation at *k* = 8, 32, 128 (modeled on Cardinal’s `spatialKMeans` (*9*)). (**C**) Boundary enrichment versus local surroundings for the same methods, stratified by anatomical superclass. Values are means across four hemispheres; error bars and shaded bands show standard deviations. The annotations are anatomical references rather than molecular ground truth.

SoLaCE and RGB MSI visualizations therefore provide different types of information. RGB visualizations encode regional molecular composition using color and can separate tissue states within relatively homogeneous regions. SoLaCE deliberately discards this color information and focuses instead on molecular boundaries. This makes it useful both as a standalone boundary-focused summary and as the high-dimensional reference edge map used for visualization evaluation through SpecEdge-Dice.

### pMiCS enables full-section MSI visualization with coherent molecular contrast

Large MSI sections require visualizations that remain interpretable at diagnostic resolution while preserving biochemically meaningful tissue transitions. pMiCS addresses this requirement by turning nonparametric MiCS+LMC into a parametric spectral-to-RGB mapping, in the spirit of parametric dimensionality-reduction methods (*5, 21*, 22) (see also Supplementary Materials 2). During training, MiCS+LMC distills several clustering-based segmentations of the MSI data into a single continuous visualization, so the final image reflects multiple segmentation resolutions rather than a single hard partition. Because pMiCS learns this mapping from sampled pixels, training remains practical on large MSI sections; the trained network can then be applied efficiently to the remaining pixels and, when preprocessing and feature alignment are matched, to new MSI images containing similar molecular content.

Representative outputs show spatially coherent RGB structure across different tissue types (Figure 4). The visualizations reveal major brain regions and substructures, glomerular and tubular organization in kidney, spinal cord architecture, and molecular heterogeneity in metastases while avoiding per-image non-linear re-optimization at full resolution. In the benchmark with pathologists, pMiCS and TOP3 (*2*) ranked highest under both sampling strategies and pMiCS performed strongly on the proposed pathologist-aligned metrics. Section-level inferential results based on the pathologist rankings are reported in Supplementary Figure S13.

**Figure 4:**
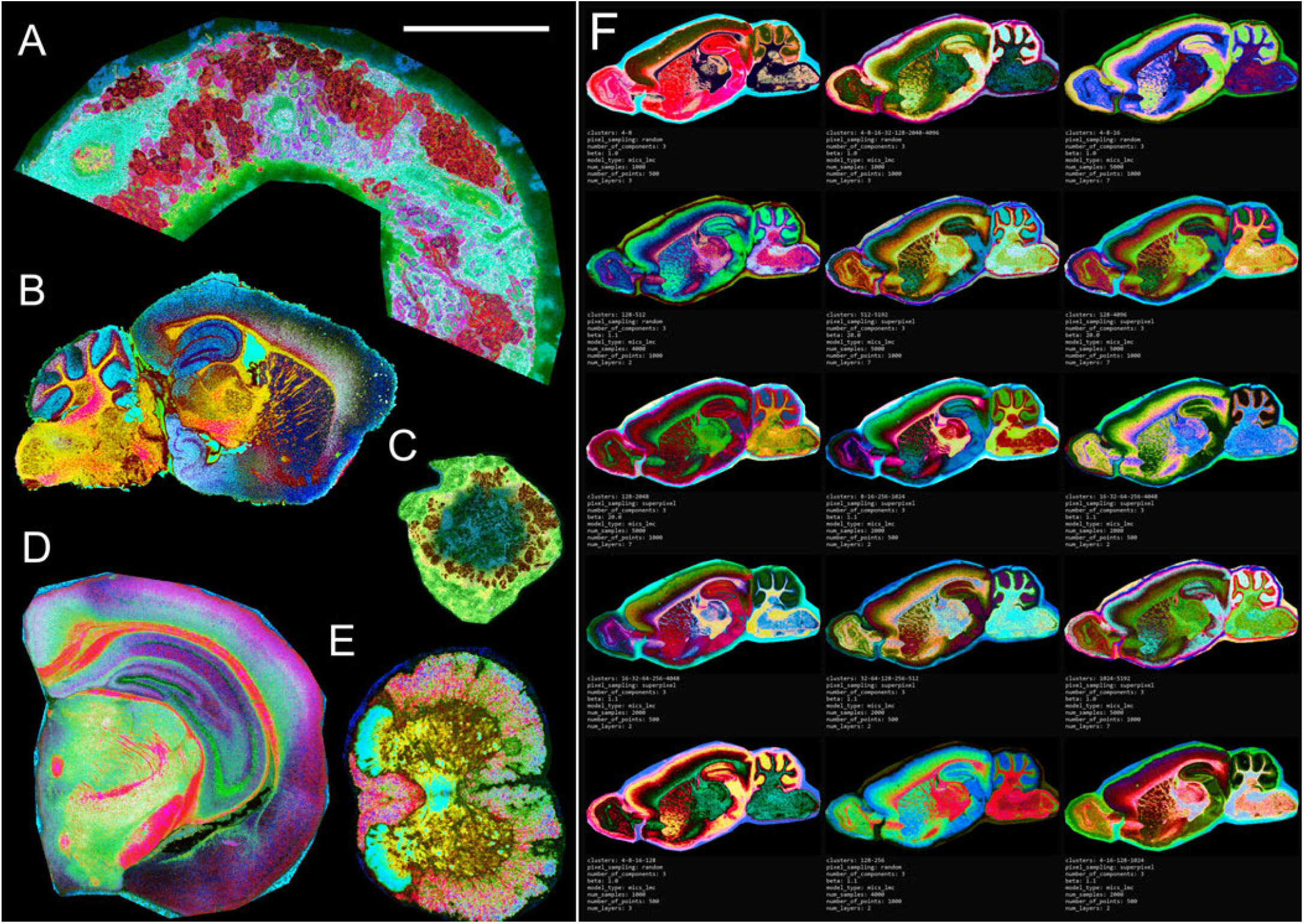
Examples of pMiCS visualizations across tissue types and different parameter settings. Representative full-section visualizations are shown for (**A**) a human kidney needle biopsy with glomerular, tubular, and fibrotic regions (5 *µ*m spatial resolution); (**B**) a sagittal mouse brain section (20 *µ*m spatial resolution); (**C**) a colorectal cancer liver metastasis; (**D**) a mouse brain hemisphere containing the cortex, hippocampus, and brainstem (5 *µ*m spatial resolution); and (**E**) a mouse spinal cord cross section showing gray and white matter (5 *µ*m spatial resolution). Panel (**F**) shows 15 views of a sagittal mouse brain section generated using different pMiCS parameter settings to enhance specific anatomic structures; enlarge the figure to read the parameter tables beneath the views. The learned parametric mapping produces spatially coherent RGB contrast across these tissues and enables direct projection when preprocessing and feature alignment are matched (see also Supplementary Figure S5). Scale bars: (**A**) 1.00 mm, (**B**) 4.14 mm, (**C**) 8.45 mm, (**D**) 2.18 mm, (**E**) 1.25 mm, and (**F**) 8.20 mm.

To demonstrate whether the learned parametric mappings can be reused beyond the sampled training pixels, we trained pMiCS and pUMAP on 5,000 valid pixels from each source image, corresponding on average to approximately 7% of valid pixels, and projected the remaining pixels out of sample. We then applied the model trained on each source image to the other images using matched preprocessing and feature alignment. This comparison demonstrates the practical advantage of parametric visualization: pMiCS and pUMAP can render unsampled pixels without re-optimizing the embedding, whereas PCA provides a linear reference projection. Within each row of Figure S14, pMiCS preserves visually consistent color semantics across the projected images, supporting scalable full-image rendering and cross-image comparison.

Runtime measurements on the brain benchmark images further support the practical value of pMiCS for iterative visualization (Figure S15). In this comparison, pMiCS and pUMAP were trained on 500 sampled pixels and executed on GPU, whereas PCA used the scikit-learn CPU implementation. pMiCS had the lowest average runtime for both training and inference under these practical benchmark settings. The pUMAP settings used here were intentionally reduced from the default configuration, because default pUMAP training required approximately five hours per image on these datasets and was therefore not practical for repeated visualization sweeps. Thus, the runtime comparison should be interpreted as a practical workflow comparison under constrained pUMAP settings, not as an exhaustive optimization of all possible pUMAP configurations.

The pMiCS and SoLaCE views are complementary. pMiCS provides color-coded regional molecular contrast, whereas SoLaCE highlights molecular transitions supported directly by the spectra rather than by RGB contrast alone. Together, these two visualizations provide the basis for assembling pathology-oriented MSI panels that combine regional composition with explicit molecular boundary information.

### Concept-based interpretation separates molecular presence from model usage

Beyond revealing tissue structure, an MSI visualization is more useful when one can ask why a given pixel receives a particular color. This analysis can prioritize pixels or regions for targeted molecular follow-up and identify when a visualization emphasizes only part of the underlying spectral information; by itself, however, it does not establish molecular identity, cell type, or mechanism. We therefore added a concept-based interpretation step that explains the frozen pMiCS output at each pixel using sparse mixtures of spectral concepts (the concept-based interpretation subsection of Materials and Methods). The aim is to separate two questions that are often conflated during visual review: which molecular concepts are present in the pixel, and which of those concepts the visualization model uses to generate the displayed contrast.

The procedure first learns nonnegative spectral concepts from the MSI matrix and then optimizes, for each pixel, a sparse normalized mixture of these concepts that reproduces the pMiCS embedding or color. The resulting explanation weights summarize model usage, whereas the original NMF activations summarize molecular abundance. Comparing these two vectors provides a per-pixel measure of agreement between molecular presence and visualization usage. Low discrepancy indicates that pMiCS uses concepts in proportion to their local abundance, whereas high discrepancy indicates that the visualization emphasizes a subset or reweighting of the available molecular concepts.

In representative examples, this interpretation converts a color-coded pMiCS image into ranked molecular concept maps that can be inspected through their underlying spectra and dominant *m*/*z* peaks (Figures 5 and 6). For four selected concepts, Figure 5B shows corresponding ion images on two independent sections, and Supplementary Table S1 provides the full ordered feature lists. This approach increases transparency without requiring complete molecular annotation of every peak: the image remains a pathology-oriented overview, while selected pixels or regions can be traced to sparse spectral concepts for targeted molecular interpretation.

**Figure 5:**
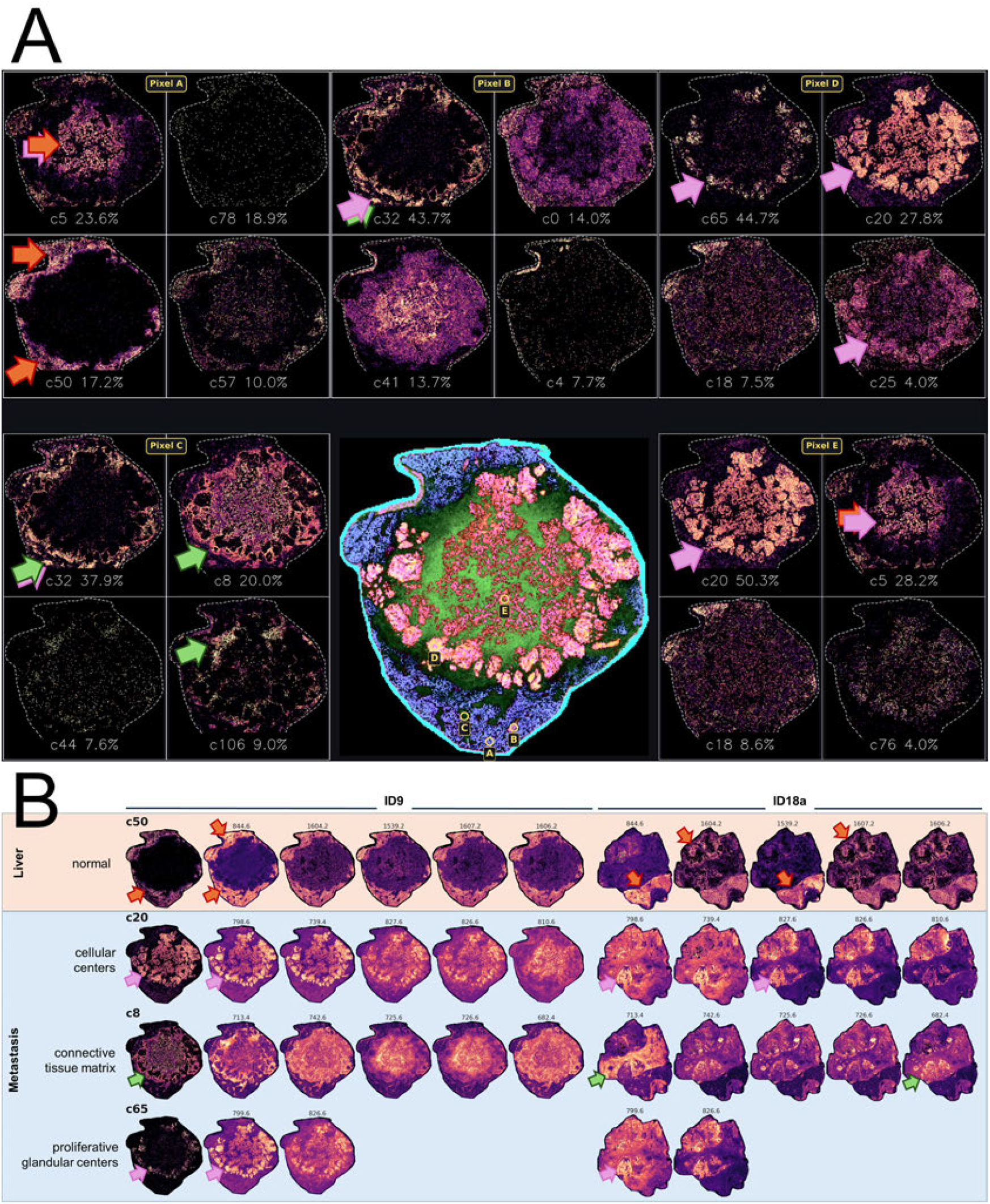
Concept-based interpretation of a pMiCS visualization of a colorectal cancer liver metastasis. (**A**) The central panel is a pMiCS rendering of a adenocarcinoma metastasis and surrounding liver; pixels A–E are each explained by their four highest-weighted NMF concepts (spatial maps, with concept labels and optimized explanation percentages). The five locations— peripheral liver, the tumor–liver interface, a peripheral island, a proliferative-appearing nodule, and the lesion center—require different concept mixtures, so the contrast is not produced by a single component. Percentages are relative explanation weights, not molecular abundances or cell-type assignments. (**B**) Ion images of the defining *m/z* for four concepts (c50, normal liver; c20, cellular centers; c8, connective-tissue matrix; c65, proliferative glandular centers) in two independent sections (ID9 and ID18a, positive-ion mode). For each concept, *m/z* values are ranked by TIC-normalized loading; among the peaks accounting for the first 50% of the loading, those with spatial enrichment ≥ 1 are ordered by enrichment, and up to five are shown (leftmost: the ID9 concept map). Enrichment is the concept-weighted mean ion intensity over tissue divided by its overall mean (1 indicates no preferential localization). The same *m/z* values recapitulate corresponding anatomical patterns in ID18a (arrows mark corresponding regions), indicating cross-section consistency. Each map is scaled to its own 1st–99th percentile; off-tissue pixels are shown in white. Full feature lists are provided in Table S1.

**Figure 6:**
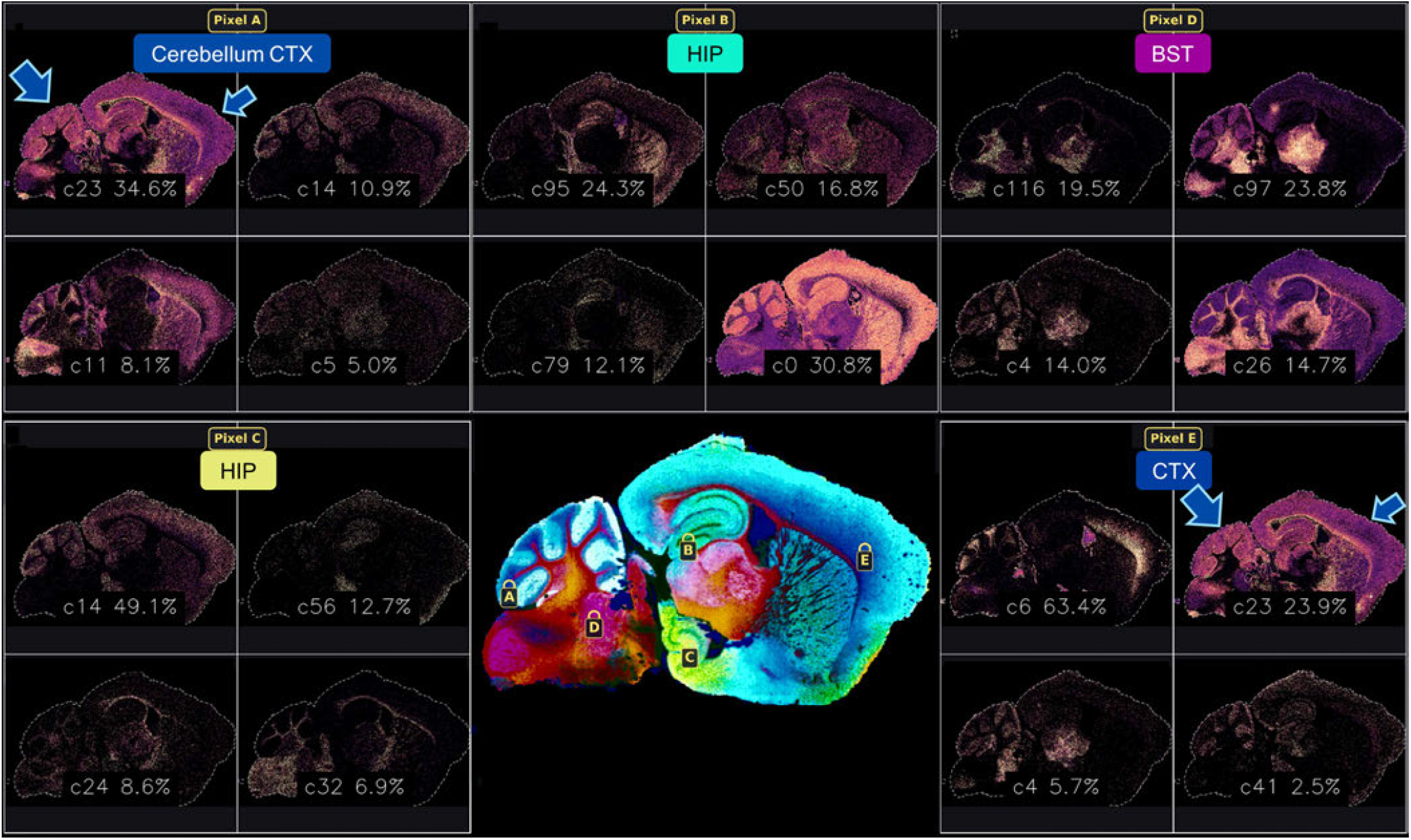
Concept-based interpretation of a pMiCS visualization of mouse brain. The central panel is a pMiCS rendering of a sagittal mouse-brain section; pixels A–E (cerebellar cortex, dorsal and ventral hippocampus, brainstem, and lower frontal cortex) are each explained by their four highest-weighted NMF concepts (spatial maps, with concept labels and optimized explanation percentages). Anatomically distinct locations—including separate sites within the hippocampal formation—require different concept mixtures, so the regional color contrast is not produced by a single component. Percentages are relative explanation weights, not molecular abundances or anatomical or cell-type assignments.

Two clarifications delimit this interpretation. First, each concept is a weighted spectrum with a spatial activation map, not a single molecule; its dominant *m*/*z* peaks can be read directly from the concept spectrum and traced to the original MSI data. Second, because many concept mixtures can reproduce the same three-channel model output, the recovered mixture is a *model-output-equivalent* summary of what the visualization uses rather than a unique or causal set of molecular drivers, a limitation shared by concept-and attribution-based explanations generally. The associated deletion test (Materials and Methods) supports the relevance of the identified concepts to the displayed color under the tested perturbation protocol; it establishes model relevance rather than biological truth. The sparse reconstruction also reproduces the model’s rendered color to within a few percent (median rendered-color error 0.014–0.027; Supplementary Figure S8), indicating that the concepts capture what the model displays even where the full spectrum is only approximately reconstructed. We accordingly distinguish three levels of evidence: association with the displayed visualization, association with specific spectral features (the dominant *m*/*z* of a concept), and confirmed molecular identity, which requires tandem MS or authentic standards and is not claimed here.

### Pathologist rankings place pMiCS in the leading tier

A blinded ranking benchmark compared five visualization methods across eight MSI images and repeated seeds. The pathologists independently ranked each image–seed comparison (Figure 7). TOP3 had the lowest mean rank under both sampling strategies (1.92 random; 2.00 superpixel-guided), with pMiCS second (2.59 and 2.17, respectively). In head-to-head rankings, pMiCS was placed above NMF, pUMAP, PCA, and TOP3 in 57%, 75%, 75%, and 34% of random-sampling rankings, respectively; the corresponding superpixel-guided values were 70%, 84%, 81%, and 48%. Agreement among pathologists was above chance (Kendall’s *W* = 0.67 for random sampling and *W* = 0.78 for superpixel-guided sampling; chance = 0.25). Thus, pMiCS ranked ahead of baselines in most comparisons, while TOP3 was preferred head-to-head in brain tissue.

**Figure 7:**
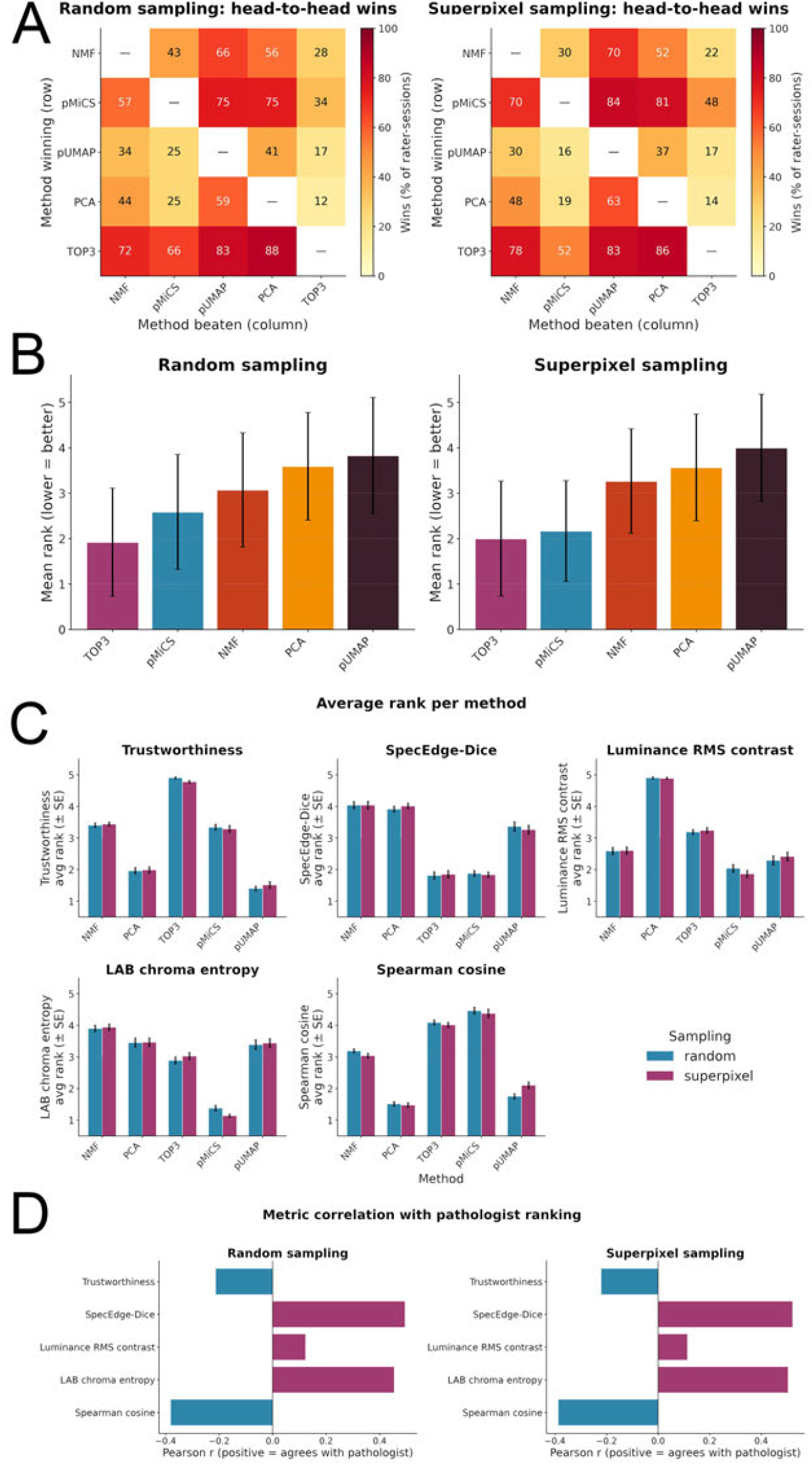
Pathologist rankings and quantitative evaluation of MSI visualizations. The pathologists ranked five methods per image–seed comparison. **A**) Pairwise win matrices show the percentage of rankings in which the row method outranked the column method. (**B**) Mean pathologist rank by method and sampling strategy (lower indicates stronger preference); error bars show the standard deviation across pathologists and sessions. (**C**) Mean method ranks for trustworthiness, SpecEdge-Dice, luminance RMS contrast, LAB chroma entropy, and Spearman cosine; error bars show standard errors. (**D**) Pearson correlations between each metric and mean pathologist ranking; positive values indicate association with pathologist preference.

Section-level tests of the pathologist rankings use the eight tissue sections as independent units (Supplementary Figure S13). After Holm correction, pMiCS had a significantly lower mean rank than PCA under random sampling (2.59 versus 3.56; *p* = 0.047) and than pUMAP under superpixel-guided sampling (2.17 versus 4.00; *p* = 0.031). Other pMiCS-versus-baseline mean-rank comparisons and all pairwise win-count tests did not reach corrected significance. pMiCS was not statistically distinguishable from TOP3 in either sampling condition (two-sided *p* = 0.109 and *p* = 0.844). Given the eight independent sections (six brain hemispheres, one kidney, one spinal cord), these inferential results remain exploratory.

This ranking result is important for pathology-scale MSI because the most flexible nonparametric embeddings are difficult to apply directly to large full-resolution images. In contrast, pMiCS trains on sampled pixels and then projects the full image through the learned spectral-to-RGB function. This makes it practical to compare many candidate visualizations, inspect full tissue sections, and assemble pathology-oriented panels even for images that are otherwise difficult to visualize interactively.

The metric analysis compared each quantitative score with the mean pathologist ranking. Trustworthiness and Spearman cosine correlated negatively with preference under both sampling strategies, indicating that these geometric criteria did not track the visual properties favored in this review. SpecEdge-Dice showed the strongest positive correlation with pathologists’ preferences (*r* = 0.50 for random sampling and *r* = 0.52 for superpixel-guided sampling), followed closely by LAB chroma entropy (*r* = 0.45 and *r* = 0.50; Figure 7). These associations support joint consideration of boundary fidelity, color variety, and geometric diagnostics; they do not establish biological validity.

Two controlled interventions clarify why these axes are complementary rather than interchangeable. First, global histogram equalization of each RGB visualization – which spreads colors to reveal more anatomical detail – raises SpecEdge-Dice and luminance RMS contrast for almost every method while lowering the geometric metrics (trustworthiness, Spearman cosine, and average image saliency). Thus, a transform that improves contrast and boundary fidelity can reduce geometric fidelity (Supplementary Figure S11). Second, a converse control guards against interpreting the boundary and color metrics in isolation: rendering *k*-means segmentation directly as a visualization (one arbitrary color per cluster) and increasing the number of clusters *k* drives SpecEdge-Dice and LAB chroma entropy above every dimensionality-reduction baseline while the Spearman correlation collapses toward zero. A fine segmentation can therefore appear structured and score well while retaining no information about which regions are molecularly similar (Supplementary Figure S12). High boundary and color scores are therefore necessary but not sufficient and must be interpreted jointly with the geometric metrics against which they can be traded; visualization quality is inherently multi-objective.

### Virtual Pathology Panels balance view quality and spatial complementarity

A useful multi-view panel requires more than several individually high-scoring renderings: additional views should also contribute information not already represented. We therefore compare greedy, spatial, and coreset selection against a random baseline across five MSI sections (Figure 8; Supplementary Figure S6). Greedy selection prioritizes whole-image quality, spatial selection maximizes marginal local-quality gains, and coreset selection prioritizes coverage in DINOv2 embedding space. Three measures track these objectives as gallery size increases: the mean local-quality envelope, embedding covering radius, and weakest included whole-image score (defined in the Panel construction subsection of Materials and Methods). These measures describe complementary aspects of selection, rather than provide independent biological validation.

**Figure 8:**
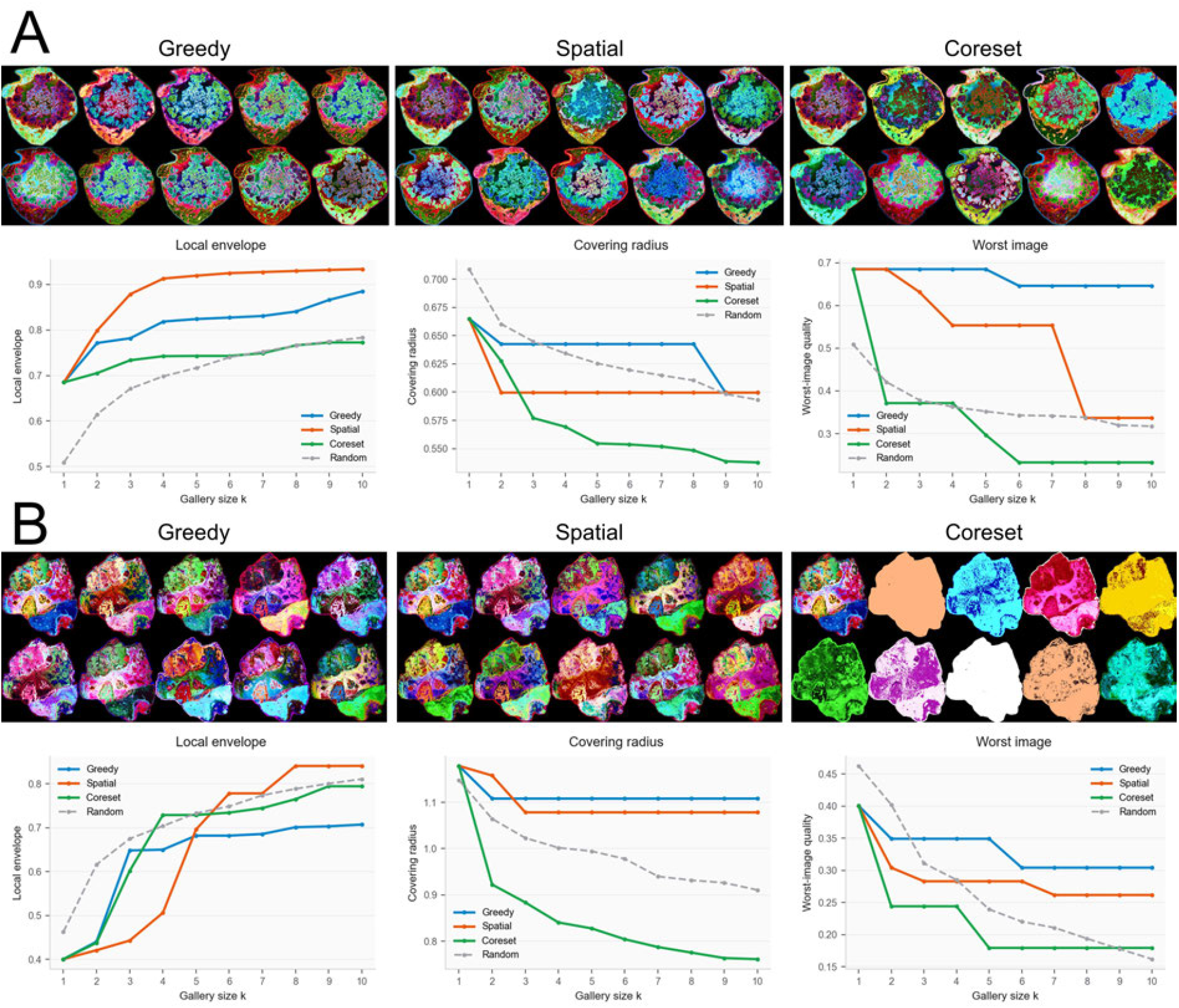
Gallery-construction trade-offs in two colorectal cancer liver metastases. ID9 (**A**) and ID18a (**B**). Image grids show ten pMiCS views per selection strategy. Curves report the mean local-quality envelope (higher is better), DINOv2 embedding covering radius (lower is better), and weakest included whole-image quality score (“Worst image”; higher is better) as functions of gallery size *k*. Blue, orange, and green denote greedy, spatial, and coreset selection, respectively; gray dashed curves denote random selection. Greedy includes similar high-scoring renderings, whereas coreset includes near-uniform views in ID18a despite improving embedding coverage. Spatial selection achieves the highest local envelope at *k* = 10 in both sections, but does not lead on every measure or at every gallery size. These measures reflect selection objectives, not independent biological validation. Supplementary Figure S6 extends the comparison to five MSI sections.

The comparisons reveal distinct trade-offs. Greedy selection can include near-duplicate high-scoring views, whereas coreset selection can include near-uniform renderings that improve embedding coverage but reveal little tissue detail. In ID9 and ID18a, spatial selection achieves the highest local envelope at ten views, although its advantage depends on section and gallery size and does not extend to every measure. Thus, increasing the number of views does not itself establish complementarity. Spatial selection instead favors views that improve tissue locations represented weakly by the current panel. Example spatially selected and quality-ranked kidney panels are shown in Supplementary Figures S17 and S18, respectively, each accompanied by a SoLaCE boundary map.

Quality-ranked VPPs illustrate the resulting review workflow in two colorectal cancer liver metastases from dataset D7. Figure 9 shows the rounded ID9 lesion and surrounding liver; Supplementary Figure S20 shows ID18a, including residual liver at the lower right and compact and gland-forming tumor regions. Positive- and negative-ion-mode views provide complementary spectral contrast for reviewing these morphologically defined compartments. Recurring spatial patterns across views nominate regions for targeted *m*/*z* analysis. The panel format is loosely analogous to reviewing complementary stains, but its colors represent unsupervised spectral phenotypes rather than known molecular targets or definitive cell types.

**Figure 9:**
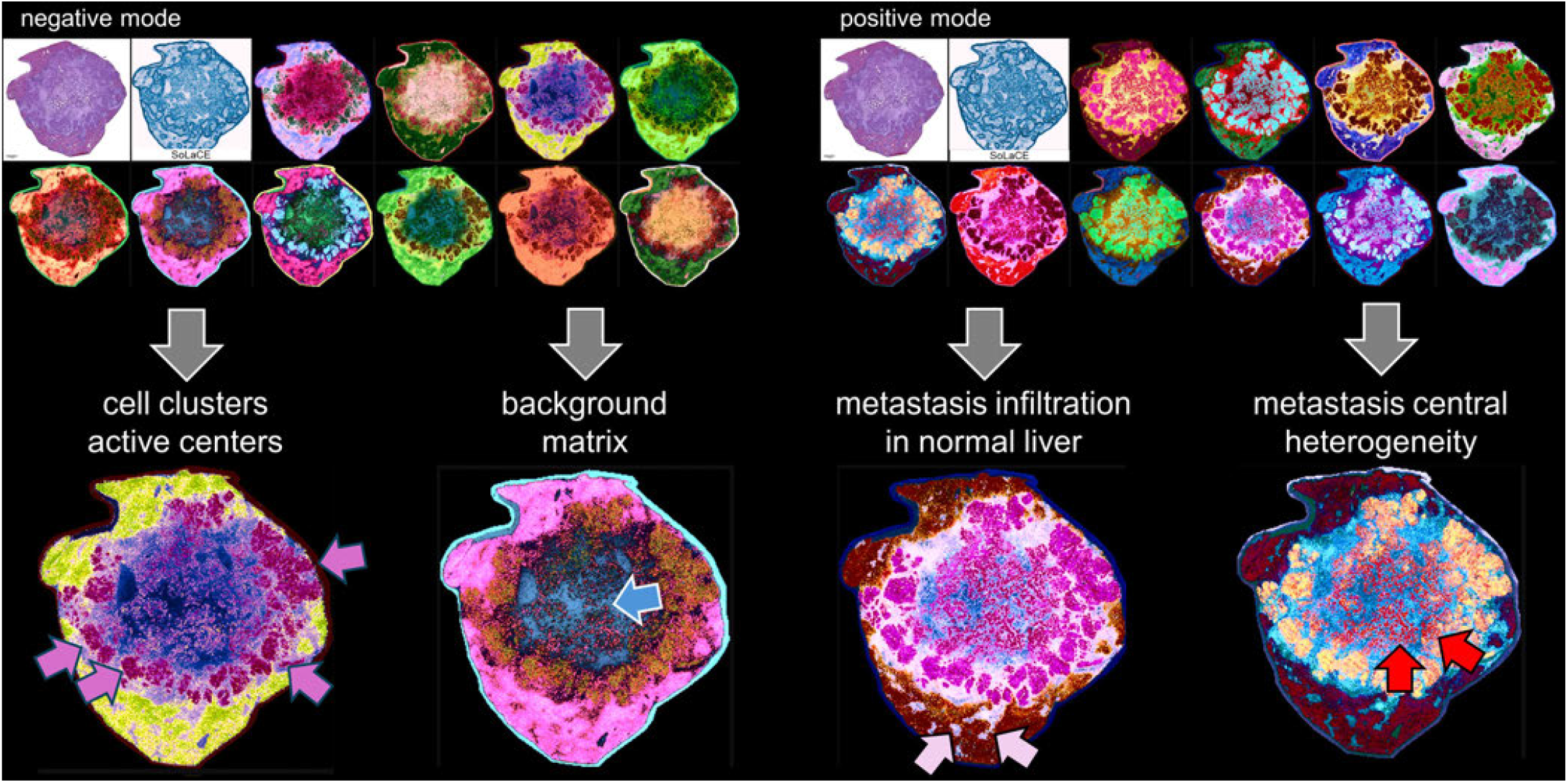
Quality-ranked Virtual Pathology Panels of a colorectal cancer liver metastasis (ID9). Negative-ion-mode (left) and positive-ion-mode (right) panels each show H&E, the SoLaCE boundary map, and ten highest-scoring pMiCS views. The full-page H&E image is provided in Supplementary Figure S19. Enlarged renderings below highlight, from left to right, peripheral cellular clusters and nodular centers, a putative matrix-associated compartment, the tumor–liver interface, and intratumoral heterogeneity with nodular foci (arrows). These author-selected examples relate to the concept analysis in Figure 5 and nominate regions for targeted *m/z* mapping. Recurrence across views is a qualitative observation, not independent validation. Anatomical descriptors follow H&E morphology; colors represent spectral phenotypes rather than confirmed cell types, molecular identities, or proliferative activity.

### Virtual Pathology Panels link tumor morphology with spectral heterogeneity

VPPs combine H&E morphology, molecular boundary maps, complementary positive- and negative-ion-mode pMiCS views, and targeted ion-level analysis. In ID9, these views distinguish the main metastatic mass, residual liver, a putative expanding border region, and a small putative invasive cell group (Figure 10A). Virtual pathology stains (VPS) relate the displayed tissue patterns to individual *m*/*z* signals.

**Figure 10:**
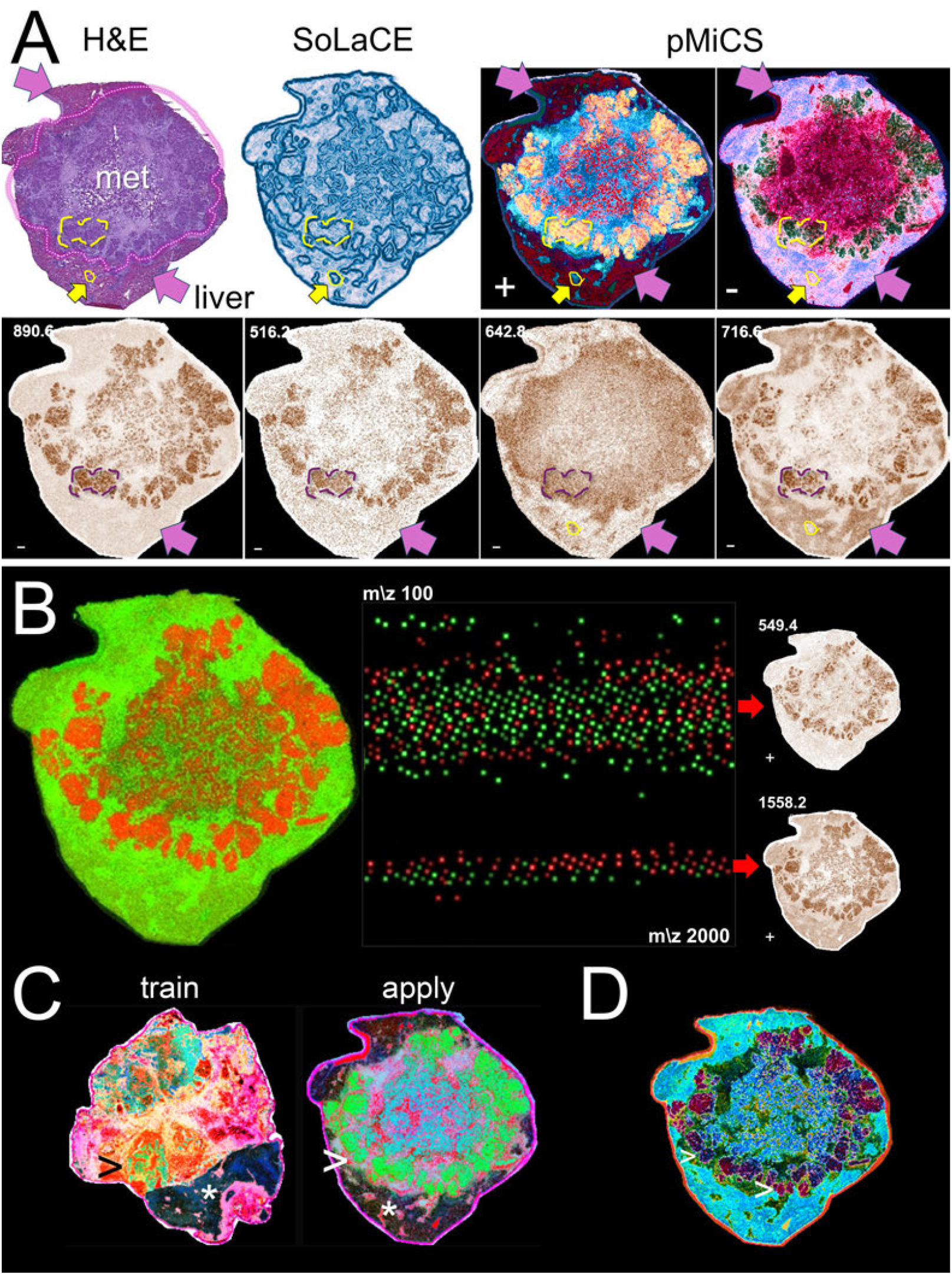
Morphological context, spectral heterogeneity, and cross-section projection in colorectal cancer liver metastases. (**A**) ID9 shown by H&E, SoLaCE, and positive-(+) and negative-ion-mode (−) pMiCS. The dotted outline marks the adenocarcinoma metastasis; magenta arrows indicate residual liver tissue. Yellow outlines mark a putative expanding border region and a small putative invasive cell group. VPS (*2*) show *m/z* 890.6 and 516.2 in proliferative-appearing nodules, *m/z 6*42.8 in tumor and putative infiltrative structures, and *m/z* 716.6 in tumor and liver regions. (**B**) Green and red ROIs compared using MSI-VISUAL (*2*). SpecQR summarizes differential signals across *m/z* 100–2000; positive-ion-mode VPS show *m/z 5*49.4 and 1558.2 enriched in nodular regions marked by the red ROI. (**C**) A pMiCS model trained on ID18a (left) is applied to ID9 (right) with matched preprocessing and spectral features. Residual liver appears dark blue to black (asterisk) and glandular regions light green (arrows) in both sections. (**D**) Coarse and fine MiCS clustering heads (*k* ∈ {8, 16, 8192}) highlight spectral differences among ID9 tumor nodules (arrows); *k* denotes cluster counts. Anatomical descriptors derive from H&E morphology. Colors and ion patterns indicate exploratory spectral associations, not confirmed molecular identities, proliferation markers, or clonal relationships.

Comparing selected regions with SpecQR further identifies differential signals, including *m*/*z* 549.4 and 1558.2 enriched in the nodular regions marked by the red ROI (Figure 10B). Together, these observations nominate spatially defined regions and spectral features for molecular identification and hypothesis testing. To illustrate cross-section reuse, a pMiCS model trained on ID18a was applied to ID9 with matched preprocessing and spectral features (Figure 10C). Residual liver appears dark blue to black, whereas morphologically interpreted glandular, proliferative regions appear light green in both sections. These corresponding color patterns demonstrate reuse of a shared mapping in the tested pair.

Within ID9, combining coarse and fine MiCS clustering heads with *k* ∈ {8, 16, 8192} highlights spectral differences among tumor nodules (Figure 10D). Each *k* specifies the cluster count of a separate segmentation contributing to the rendering, rather than the number of panel views. The resulting contrast supports targeted comparison of intratumoral spectral heterogeneity.

## Discussion

This study presents a linked pathology-oriented MSI workflow: SoLaCE extracts molecular boundaries directly from spectra, pMiCS provides scalable parametric color visualization, concept-based interpretation connects contrast to spectral concepts, and pathologist-aligned metrics support VPP construction. Each component addresses a different limitation of current MSI visualization—boundary visibility, computational scalability, molecular traceability, or view selection. Together, they enable complementary boundary-focused and color-coded views rather than relying on a single embedding.

### SoLaCE provides a molecular boundary view of MSI

A central result is that molecular boundary detection is itself a useful MSI representation, not only an intermediate for visualization scoring. SoLaCE summarizes changes in high-dimensional spectral composition while suppressing within-region texture. This is important because a biochemical transition may be distributed across correlated *m*/*z* channels even when total signal abundance changes little. Unlike TIC Sobel (*23*), which responds to total-intensity gradients, or direct spectral-difference magnitude, which can be dominated by isolated high-distance pixels, SoLaCE produced coherent internal boundaries and tissue interfaces across kidney, brain, spinal cord, and colorectal cancer liver metastasis sections (Figure 2). The resulting map compresses a high-dimensional transition signal into a form that can be inspected alongside conventional pathology images.

The anatomical benchmark supported this interpretation. SoLaCE had the highest border enrichment, average precision, ROC-AUC, and mean F1 among the evaluated methods, and the highest density-matched F1 at each tested edge density. The densest spatial *k*-means segmentation (*k* = 128) was slightly higher on border-to-surrounding enrichment, showing that no method dominated every measure. Separately, one expert pathologist reviewed SoLaCE and the difference-magnitude and TIC-Sobel baselines across the qualitative comparison set. Performance also varied across anatomical superclasses, confirming that molecular transitions do not coincide uniformly with every annotated border. These results support SoLaCE as a continuous molecular-boundary representation, not as a replacement for histological segmentation or pixel-level molecular ground truth.

SoLaCE serves two related roles. As a standalone view, it directs attention to tissue compartments, lesion interfaces, and small spatially distinct regions; in the liver metastasis example, it delineates the main lesion, tumor–liver interface, and putative invasive cell groups. As a spectral reference, it enables SpecEdge-Dice to assess whether RGB visualizations retain high-dimensional transitions independently of their color conventions. This complements region-labeling approaches in software such as Cardinal (*9*) and SCiLS Lab (*10*): segmentation assigns discrete region labels, whereas SoLaCE estimates continuous boundary strength. It also complements pMiCS, which uses color to encode regional spectral composition while SoLaCE emphasizes where that composition changes, helping to precisely annotate structures for further analyses.

### Efficient parametric visualization for iterative review

The practical contribution of pMiCS lies in its deployment characteristics as well as its ranking performance. Training on sampled pixels separates model selection from full-image rendering, making hyperparameter sweeps and subsequent projection of large sections feasible without repeated full-image optimization. This distinction becomes particularly important at 5 *µ*m resolution or finer and for larger whole-section MSI datasets, where computational and memory constraints can obstruct iterative review. Unlike nonparametric embeddings that must be re-optimized for each image, pMiCS supports direct out-of-sample projection and can preserve corresponding color patterns when preprocessing and feature alignment are matched. This was observed both across the related brain sections in Supplementary Figure S14 and when a model trained on metastasis section ID18a was applied to ID9 (Figure 10C), where residual liver and glandular regions retained corresponding colors. Representative sampling supports learning the source section’s spectral variation but does not by itself establish transferability. These experiments demonstrate reuse in matched settings, not independently validated tissue classification or generalization across laboratories, instruments, or acquisition protocols.

### Interpreting visualization contrast through molecular concepts

Concept-based interpretation adds a molecular explanation layer by identifying sparse NMF-derived mixtures that approximately reproduce the frozen pMiCS output. Three analyses support model-output faith-fulness under the tested protocol. Concept deletion disrupts the rendering more when concepts are removed according to optimized usage than when they are removed according to abundance or at random; the reconstructed CIELAB colors have median relative errors of 1.4–2.7% in the three illustrated sections; and the Jensen–Shannon discrepancy between explanation weights and scale-adjusted NMF concept contributions identifies where the model reweights the available spectral mixture. The discrepancy is related to, but not reducible to, pixel mixedness. Together, these analyses distinguish concepts used to reproduce displayed contrast from concepts merely abundant in the spectrum and narrow follow-up to a smaller set of dominant *m*/*z* features. They do not establish a unique or causal attribution, molecular identity, cell type, or biological mechanism; those claims still require peak annotation and orthogonal validation.

### Why color-variety and edge-aware metrics align better with pathologist judgment

The mismatch between local and global structure-preservation metrics and the pathologist ranking reflects different evaluation targets. These geometric metrics treat spectra as a point cloud and quantify neighborhood or pairwise relationships without directly considering pixel adjacency, contour continuity, or visual contrast. Because boundary pixels represent only a small fraction of an image, interfaces may be blurred, displaced, or fragmented with little effect on average structure-preservation scores; conversely, emphasizing separation between tissue states may improve pathology-oriented readability while reducing geometric fidelity. These metrics nevertheless remain valuable: good global structure preservation can support qualitative reasoning about molecular similarities between anatomically separated regions because spectrally similar regions remain nearby in the visualization space. Their weaker association with pathologist preference therefore does not invalidate them but shows that geometric fidelity alone does not describe pathology-oriented visual usefulness.

Pathology-oriented review and SpecEdge-Dice instead prioritize coherent, high-contrast spatial interfaces. Luminance RMS contrast and LAB chroma entropy quantify display variety, while SoLaCE supplies a spectral reference for assessing whether displayed edges coincide with molecular transitions. One possible explanation for TOP3’s strong SpecEdge-Dice performance is its pointwise construction: it maps the three largest spectral intensities independently at each pixel without spatial averaging. These dominant intensities may vary smoothly within homogeneous regions, whereas molecular interfaces may alter their magnitude, sparsity, or relative balance, producing sharp RGB transitions at their original spatial locations. Ignoring weaker signals may also suppress within-region spectral texture. This strong edge agreement does not imply complete spectral preservation, however, because TOP3 discards ion identity and may miss transitions that do not alter the dominant-intensity profile.

These findings favor multi-objective evaluation rather than replacement of one metric family by another. Geometric preservation, color variety, boundary fidelity, computational cost, and pathologists’ preferences describe different properties, and the appropriate balance will depend on whether the intended use is exploration, cross-sample comparison, panel construction, or targeted pathology review.

### From single images to Virtual Pathology Panels

As in routine anatomical and surgical pathology, one view may not capture all relevant structures. Efficient generation and scoring therefore turns MSI visualization into a panel-construction problem. The greedy strategy prioritizes individually strong visualizations but can select similar renderings; the spatial strategy adds the view that most improves local tissue coverage beyond the current panel; and the coreset strategy improves coverage of the candidate pool in DINOv2 embedding space but can select visually uninformative outliers. In ID9 and ID18a, spatial selection achieved the highest local envelope at ten views, but did not lead on every measure or at every gallery size (Figure 8). The five-section comparison likewise shows strategy-specific trade-offs rather than universal superiority (Supplementary Figure S6).

Panel size and composition therefore require practical calibration. The local envelope quantifies marginal coverage of tissue locations, the covering radius quantifies representation of the candidate pool in embedding space, and the weakest included whole-image score guards against adding low-quality views. These quantities are pool-relative and aligned with the corresponding selection objectives; they are not independent measures of biological validity, and covering radius is not a direct measure of visual redundancy. Combining coarse and fine MiCS clustering heads can also reveal spectral differences among morphologically related tumor nodules (Figure 10D), nominating regions for further molecular comparison and analysis. Future studies should determine whether such panels improve region identification, review time, inter-pathologist agreement, or selection of signals for follow-up compared with a single visualization.

### Limitations

This study evaluates visualization quality, not diagnostic accuracy or clinical outcomes. Lesion interfaces and spatially distinct cell groups should therefore be regarded as hypothesis-generating observations. Although four pathologists ranked the methods and section-level analyses assess these rankings, the findings do not establish diagnostic benefit or generalizable consensus; a larger, prospective, independently designed study is planned. pMiCS is tunable, thus can be adopted to the datasets, and its outputs depend on pixel sampling, model configuration, preprocessing, and feature alignment, whereas TOP3 is not tunable. Quantitative comparisons should therefore be interpreted only within the datasets and conditions evaluated here.

#### Scope of applicability

The methods operate on a spatially indexed pixel-by-feature matrix with a tissue mask, and are in that sense acquisition-agnostic once compatible preprocessing (feature alignment, total-ion-current normalization, and a common *m*/*z* binning) has been applied. They were validated on MALDI-MSI at 5–30 *µ*m spatial resolution, Bruker .d (Table 1, dataset D3,4) and imzML (*24*) standard input formats (Table 1, datasets D1,2,5,6,7,8). Applicability to other ionization or desorption modalities, for example DESI, and to coarser 50–200 *µ*m acquisitions was successfully tested, and is expected to work also for other modalities and acquisitions on algorithmic grounds. Runtime and memory — the practical limitations — scale with the number of pixels and features per dataset, the number of landmarks, and the sampling fraction.

**Table 1:** MSI datasets.

| Dataset | Used in | Organ | Species | Resolution | Data source | Link |
| --- | --- | --- | --- | --- | --- | --- |
| D1 | Benchmarking and Figures 2B, 4A, S1C, S6E, S16, S17, and S18 | Kidney needle biopsy | Human | 5 $\mu\text{m}$ | METASPACE, 2025-01-15_23h54m25s | https://metaspace2020.org/datasets?ds=2025-01-15_23h54m25s |
| D2 | Benchmarking and Figures 2D, 4E, and S1E | Spinal cord cross section | Mouse | 5 $\mu\text{m}$ | METASPACE, 2024-07-02_13h47m39s | https://metaspace2020.org/datasets?ds=2024-07-02_13h47m39s |
| D3 | Benchmarking and Figures 2A, 4D, S6CD, and S17 | Three coronal brain hemispheres | Mouse | 5 $\mu\text{m}$ | In-house, Zenodo, 23373081 | https://doi.org/10.5281/zenodo.23373081 |
| D4 | Benchmarking and Figures 3, S3, and S14 | Four coronal brain hemispheres | Mouse | 20 $\mu\text{m}$ | In-house acquisition, deposited in PRIDE (PXD056609) (1, 2) | https://www.ebi.ac.uk/pride/archive/projects/PXD056609 |
| D5 | Figures 4B, 6, and S8A | Sagittal brain | Mouse | 20 $\mu\text{m}$ | METASPACE, 2025-07-01_19h15m43s | https://metaspace2020.org/datasets?ds=2025-07-01_19h15m43s |
| D6 | Figure S1A | Brain with tumor | Mouse | 20 $\mu\text{m}$ | METASPACE, 2023-02-23_22h35m34s, (36) | https://metaspace2020.org/datasets?ds=2023-02-23_22h35m34s |
| D7 | Figures 2C,E, 4C, 5, 8, 9, 10, S1D, S6AB, S8BC, S9, S19, and S20 | Liver metastases (ID9, ID18a) | Human | 20 $\mu\text{m}$ | Zenodo, 18495815, (35) | https://doi.org/10.5281/zenodo.18495815 |
| D8 | Figures 4F, S3 | Cerebellum | Mouse | 30 $\mu\text{m}$ | RIKEN, DM00076, (37) | http://prime.psc.riken.jp/menta.cgi/prime/drop_index |

The proposed metrics and explanations assess complementary but incomplete aspects of quality. Color-variety metrics measure legibility, SpecEdge-Dice emphasizes boundary agreement and may underrepresent gradual patterns, and concept weights are model explanations rather than molecular identifications. Future studies should combine multi-reader evaluation, geometric and structure-aware metrics, direct molecular validation, improved sampling strategies, and predefined review tasks that measure region identification, review time, reader agreement, and the reproducibility and yield of molecular follow-up.

### Outlook

MSI has the potential to connect morphology-oriented tissue assessment with molecular analysis in a single acquisition. Future workflows could provide pathologists with VPP that display tissue architecture, molecular boundaries, and region-specific spectral phenotypes together. The next step is to test whether these panels improve predefined review tasks relative to a single visualization, including localization of relevant regions, consistency across readers, review time, and selection of molecular signals for validation. Clinical deployment will additionally require validated sample preparation, acquisition, annotation, quality-control, and interpretation procedures.

The same integration is relevant to pharmacological research and biomarker discovery. Spatial molecular panels could reveal heterogeneous drug responses, treatment-resistant compartments, metabolic changes, or molecularly distinct invasive margins that may be diluted in bulk-tissue analyses. SoLaCE could identify treatment-associated molecular boundaries, pMiCS could summarize regional response patterns, and concept-based interpretation could prioritize spectral features for molecular identification and validation as candidate pharmacodynamic or diagnostic biomarkers.

Continued improvements in instrumentation, computational hardware, algorithms, and data standards should make high-resolution MSI analysis increasingly practical. Faster acquisition and inference may eventually permit large tissue sections to be reviewed through interactive, multi-view molecular maps while preserving access to the underlying spectra. In this setting, MSI could become a distinctive analytical platform that complements histopathology, immunohistochemistry, genomics, and targeted molecular assays in integrated tissue diagnostics and translational research.

## Conclusions

We present pMiCS, which turns MiCS+LMC from a per-image optimization procedure into a reusable parametric mapping for MSI visualization. In our benchmark, pMiCS and TOP3 formed the leading pair; pMiCS additionally supports efficient full-image inference and projection to matched sections. We combine color-variety metrics with SpecEdge-Dice to evaluate complementary aspects of visual usefulness, while retaining geometric metrics as additional diagnostics. Concept-based explanations connect displayed contrast to sparse spectral concepts and show model-output faithfulness. Finally, VPP selection exposes trade-offs between individual view quality, local complementarity, and embedding-space coverage. Together, these components provide a scalable framework for exploratory, pathology-oriented MSI review and targeted molecular follow-up, while clinical and biological claims require independent validation.

## Materials and Methods

Supplementary Materials 2 provides explanatory schematics and concise descriptions of the equations used in the following sections.

### Parametric MiCS+LMC for MSI visualization

MiCS+LMC was introduced as a two-part objective that combines global and local structure preservation in a single training process (*19*). The LMC term (*Landmark Mantel Correlation*) promotes global faithfulness by correlating high-dimensional and low-dimensional distance structures computed with respect to a landmark set, thereby keeping computation tractable for large datasets. The term encourages preservation of large-scale relationships between spectra in the embedding.

The *Multi-resolution Cluster Supervision* (MiCS) component complements the LMC term by promoting local and multiscale fidelity. It constructs cluster assignments at multiple resolutions in the high-dimensional space and requires these assignments to remain predictable from the low-dimensional embedding. In this way, MiCS encourages the embedding to preserve neighborhood structure consistently across multiple scales.

This perspective naturally aligns with MSI visualization. Previous approaches typically rely on a single segmentation or clustering of spectra, producing discrete region assignments that are sensitive to algorithm choice and resolution. In contrast, MiCS can be interpreted as aggregating supervision from *multiple segmentations* simultaneously, each corresponding to a different resolution of the data. The resulting embedding acts as a distilled visualization that preserves structure across these multiple clusterings, rather than committing to a single partition.

We implement a parametric variant (pMiCS) by training a lightweight neural network — we tested 2–7 layers — that maps each pixel’s *m*/*z* spectrum directly to a low-dimensional embedding used to generate RGB colors. This yields a continuous, spatially coherent visualization in which similar spectra are mapped to nearby colors. Unlike hard segmentation, this formulation avoids discretization artifacts and allows smooth transitions between tissue states, which are common in MSI data.

During optimization, MiCS+LMC combines these components in a weighted objective,

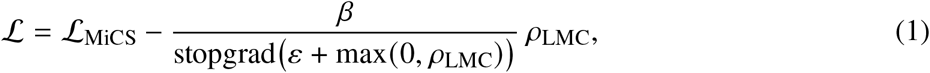

with scheduling of the LMC contribution during training (*19*).

### Concept-based interpretation of pMiCS visualizations

To interpret pMiCS visualizations, we explain the model output at each pixel in terms of molecular spectral concepts. The goal is to distinguish molecular concepts that are present in a pixel from the subset of concepts that the visualization model uses to generate the displayed contrast.

Let 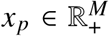 denote the preprocessed MSI spectrum at pixel *p*, where *M* is the number of *m*/*z* bins, and let 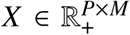 denote the full pixel-by-*m*/*z* matrix. We first decompose *X* using nonnegative matrix factorization (NMF) (*25, 26*),

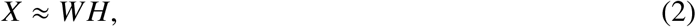

where 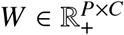 contains the activation of each concept in each pixel, and 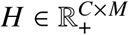 contains the learned spectral concepts. In our experiments, we used *C* = 128 concepts. The *k*-th row *h*_*k*_ of *H* represents one molecular spectral concept, which can be inspected by its dominant *m*/*z* peaks.

We then ask which sparse mixture of these concepts is sufficient to reproduce the output of the trained visualization model. Let *f*_*θ*_ denote the frozen pMiCS model, and let

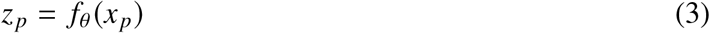

be the original model output for pixel *p* in the embedding or color space used for visualization. For each pixel, we optimize unconstrained concept logits *α*_*p*_ ∈ ℝ^*C*^. These logits are converted into a sparse normalized concept mixture using a top-*s* softmax operator,

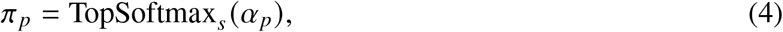

where only the *s* largest entries of *α*_*p*_ are retained, all other entries are set to zero, and the retained weights are normalized to sum to one. The explanation spectrum reconstructed from the selected concepts is

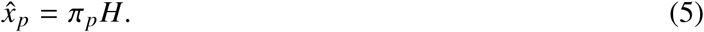

This reconstructed spectrum is then passed through the frozen visualization model,

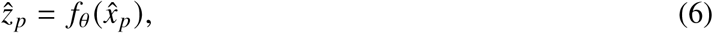

and the concept weights are optimized to preserve the original model output:

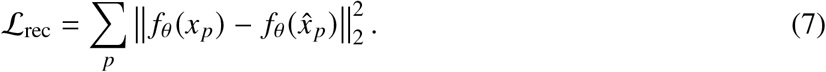

To encourage compact explanations, we add an entropy penalty on the normalized concept mixture,

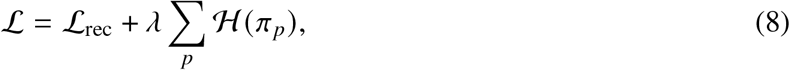

where

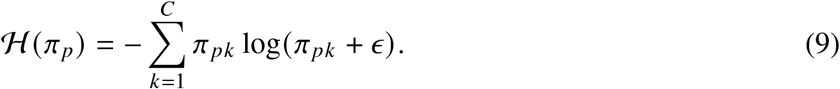

Here, *λ ≥* 0 controls the strength of the sparsity preference, and *ϵ* is a small numerical constant. Lower entropy encourages the explanation to concentrate on fewer dominant concepts.

After optimization, each pixel can be interpreted by ranking concepts according to the optimized weights *π*_*pk*_ . For selected pixels or regions, we report the highest-weighted concepts and inspect their corresponding spectra *h*_*k*_, especially the dominant *m*/*z* peaks. These optimized weights represent concept usage by the visualization model, not simply molecular abundance.

Finally, to compare model *usage* against molecular *presence*, the two are put on a common scale. Because the NMF concepts differ in magnitude, each concept (row *h*_*k*_ of *H*) is L1-normalized to unit norm and its abundance is multiplied by the same factor *l*_*k*_ = ∥*h*_*k*_∥_1_, giving a molecular-abundance distribution 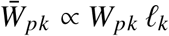 over the *C* concepts; the usage mixture *π*_*p*_ is already such a distribution. Their departure is measured by the Jensen–Shannon divergence (*27*),

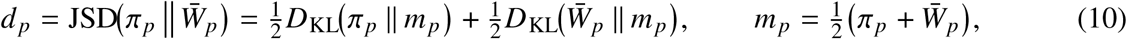

where *D*_KL_ is the Kullback–Leibler divergence (*28*) and *m* _*p*_ is their midpoint distribution. Unlike a cosine score, the Jensen–Shannon divergence is a bounded, symmetric comparison of two *distributions* (*d*_*p*_ ∈ [0, 1] with base-two logarithms) and is therefore directly comparable across pixels. A low *d*_*p*_ means the model uses concepts in proportion to their abundance; a high *d*_*p*_ means the visualization emphasizes a subset or reweighting of them.

To check that the usage weights *π*_*p*_ are meaningful rather than an arbitrary fit, we apply a concept-deletion test in the spirit of perturbation-based faithfulness measures (*29, 30*). The top-*K* concepts under a given ranking are deleted, the mixture is re-optimized on the remainder, and the resulting change in the pixel’s rendered LAB color, relative to the full-mixture rendering, is measured (Supplementary Figure S7). The same deletion and re-optimization is applied to all three rankings, so the comparison isolates the effect of the ranking rather than of the refitting procedure. Removing the concepts identified by the optimized usage (*π*) disrupts the visualization more than either baseline at every budget: more than random throughout, and more than the most abundant concepts with the largest margin at small *K*, narrowing as *K* grows and nearly closing by *K* = 16. The margin over the abundance ranking is itself informative: the model does not simply use the most abundant concepts, which is precisely the premise *d*_*p*_ measures. Read against the pixel’s abundance entropy 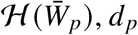, *d*_*p*_ is related but not reducible to how mixed the pixel is: across sections the per-pixel Spearman correlation between *d*_*p*_ and 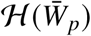 is positive but only moderate (median 0.255, negative on a minority of sections), so *d*_*p*_ tracks where model usage departs from abundance rather than re-encoding pixel mixedness (Supplementary Figures S9 and S10).

### High-dimensional edge detection for MSI

To extract molecular boundaries directly from the spectra, we computed high-dimensional edge maps using a soft landmark contrast approach. Let *x*_*i*_ ∈ ℝ^*M*^ denote the spectrum at pixel *i*, where *M* is the number of *m*/*z* bins, and let *z*_*i*_ denote the tissue spectrum after TIC normalization, projection onto the top 64 PCA components (fit on tissue pixels), and *l*_2_ normalization. A set of *L* = 1,000 landmark spectra *L*_land_ = {*l*_1_, …, *l*_*L*_} was sampled uniformly from valid tissue pixels. Each pixel was represented by its cosine similarity to the landmarks,

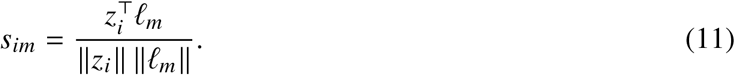

For each pixel, we retained the top *K* = 80 landmarks,

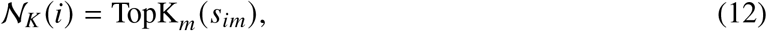

and converted these similarities into a sparse soft landmark signature,

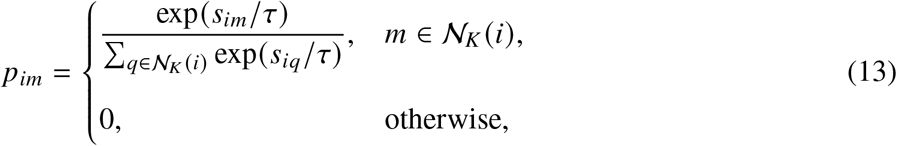

where *τ* = 0.06 is a softmax temperature. To reduce pixel-level noise, signatures were locally averaged over a 3 × 3 patch *P*(*i*) centered on each pixel,

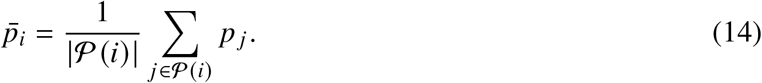

Edge strength was then estimated by comparing patch-averaged signatures on opposite sides of each pixel. For direction *d*, the across-boundary contrast was

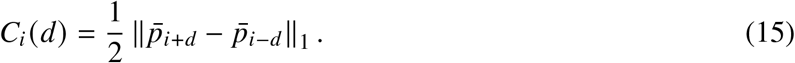

To suppress responses from internally heterogeneous regions, we subtracted a local texture term,

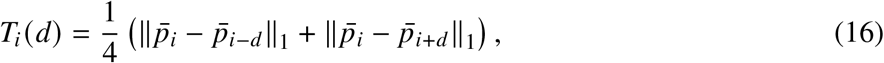

giving the directional score

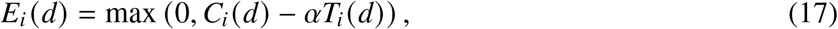

where *α* = 0.22 controls texture suppression. The final edge map was obtained by aggregating over the two undirected spatial axes *D* = {(1, 0), (0, 1)},

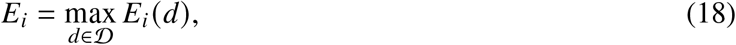

followed by percentile normalization for visualization. This procedure produces high-dimensional molecular edge maps that can be compared directly with edges extracted from RGB visualizations. The sparse landmark representation emphasizes changes in local spectral relationships and can yield sharper boundaries than direct contrast of the continuous spectra. An ablation of these components is shown in Supplementary Figure S3.

### Evaluation against expert anatomical boundaries

Because the expert annotations delineate anatomical regions rather than molecular transitions, we did not treat them as pixel-level molecular ground truth. Instead, we evaluated whether each soft edge map assigned high edge strength to expert-defined anatomical borders while suppressing responses within annotated regions and their immediate surroundings. The reference annotations comprised polygon annotations (*31*) from four mouse brain hemispheres from MSI-ATLAS (*1*). Background polygons were excluded from all analyses. In addition to the quantitative evaluation, all SoLaCE edge maps and their corresponding difference-magnitude and TIC-Sobel baseline maps were reviewed qualitatively by a single expert pathologist. This descriptive assessment was not treated as an independent quantitative endpoint. The benchmark design and quantitative results are summarized in Figure 3.

We compared SoLaCE with three baseline methods. For the *difference-magnitude* baseline, TIC-normalized spectra were percentile-scaled independently for each *m*/*z* channel. At each tissue pixel, the edge score was the maximum *L*_2_ spectral distance to neighboring tissue pixels within a one-pixel spatial radius. Scores were then z-score normalized across tissue pixels. For the *TIC-Sobel* baseline, a total-ion-current image was generated from the unnormalized spectra, percentile-stretched, and processed with a Sobel operator using a 3 × 3 kernel. For the *spatial k-means* baseline, we followed Cardinal’s Gaussian `spatialKMeans` (*9*): each spectrum was TIC-normalized and Gaussian-smoothed over a Chebyshev radius-1 (3 × 3) neighborhood (standard deviation (2*r* + 1)/4 = 0.75, with weights renormalized over tissue pixels so that off-tissue pixels do not contribute), using Euclidean distances between smoothed spectra (Cardinal’s `average` metric); Fastmap then embedded the tissue pixels into 10 components (one pivot pass each), and *k*-means was run on the scores for *k* ∈ {8, 32, 128} (10 random starts, 10 iterations; `sklearn` Lloyd, seed 0, in place of Cardinal’s Hartigan–Wong). Its edge map is the 4-connected boundary between cluster labels, with off-tissue pixels left unlabeled. Before evaluation, each continuous edge map was rank-normalized over tissue pixels to place the methods on a common scale.

Expert anatomical boundaries were derived from the polygons. Polygon outlines were dilated by one pixel to form reference boundary bands. Region interiors were defined as the filled polygons after removal of the corresponding boundary bands. For comparisons with the local surroundings, a three-pixel exterior ring was constructed around each annotated region.

Across the union of all annotated regions, we computed four groups of measures:

1. **Border-to-interior enrichment**, defined as the mean edge score on reference boundaries divided by the mean edge score within annotated interiors.
2. **Border-to-surrounding enrichment**, defined as the mean edge score on reference boundaries divided by the mean edge score within the three-pixel exterior rings.
3. **Boundary-ranking performance**, quantified by average precision (AP) and the area under the receiver operating characteristic curve (ROC-AUC) for discriminating boundary pixels from interior pixels using continuous edge scores.
4. **Density-matched detection performance**, quantified by the F1 score after thresholding each edge map to retain the top 5%, 10%, 15%, or 20% of edge pixels within the annotated tissue. The mean F1 across these four operating points was used as a summary measure.

Results were summarized as the mean ± standard deviation across the four hemispheres.

To assess performance across tissue types, border-to-surrounding enrichment was also computed separately for each anatomical superclass (`main cat1`): white matter (WM), cortex (CTX), brainstem (BST), cerebral nuclei (CNU), amyloid-*β* plaques (PLQ), and choroid plexus (PLX). Each superclass was evaluated using its own reference boundaries and corresponding three-pixel exterior rings. Superclasses with insufficient annotated boundary pixels in a hemisphere were excluded from the analysis of that hemisphere.

### Parametric versus nonparametric embeddings in MSI

Nonparametric embeddings provide flexible geometry but typically require re-optimization for each new image. Parametric methods learn an explicit function from spectra to embedding coordinates, enabling scalable inference and consistent mapping across datasets. Parametric UMAP is a widely used example of parametric dimensionality reduction (*5*). For high-resolution MSI, this capability is important for maintaining comparable color semantics across cohorts while keeping inference costs practical.

#### Color-variety metrics

We evaluated two simple color-variety metrics directly from each RGB visualization without comparison with a high-dimensional MSI summary or an independently validated histological reference. Both were computed only over valid tissue pixels. Luminance RMS contrast was defined as the standard deviation of Rec. 601 luminance, *Y* = 0.299*R* + 0.587*G* + 0.114*B*, with RGB values scaled to [0, 1]. LAB chroma entropy was computed by converting quantized RGB values to OpenCV-style *L*^∗^*a*^∗^*b*^∗^ color space, forming a 16 × 16 joint histogram over the *a* and *b* channels, and computing Shannon entropy, *H* = −∑_*k*_ *p*_*k*_ log *p*_*k*_, over occupied bins. Higher contrast indicates stronger global light–dark separation, whereas higher chroma entropy indicates use of more diverse chromatic categories. Empty valid masks yield undefined scores.

#### Edge-based evaluation of MSI visualizations

A central goal of MSI visualization is to reveal spatial transitions between distinct biochemical states. In neuro-oncology and brain tissue analysis, these transitions can mark boundaries between biologically or pathologically distinct regions, such as tumor versus normal tissue, infiltrative margins, necrotic cores, white versus gray matter, and inflamed regions. From a pathology perspective, the delineation of such boundaries is often more informative than absolute similarity relationships, as it directly reflects underlying biological organization and supports expert interpretation. A useful visualization should therefore faithfully preserve not only local neighborhoods or global geometry, but also the location, sharpness, and coherence of these transitions.

Standard embedding metrics do not directly capture this property. Global measures, such as distance correlation, assess overall geometric fidelity but are largely insensitive to localized discontinuities. Local measures, such as trustworthiness, evaluate whether nearest neighbors are preserved, but do not explicitly penalize distortions of boundaries. In particular, boundaries typically occupy a small fraction of the data and can be blurred, shifted, or widened without substantially affecting neighborhood rankings. As a result, an embedding may achieve high scores under these metrics while failing to preserve pathology-relevant structure.

To address this limitation, we evaluate visualizations based on their ability to preserve edge structure. Given an MSI dataset defined over a spatial grid M, we compute edge maps in both the high-dimensional spectral space and the low-dimensional visualization. Let *h*_*i*_ denote the molecular edge strength at pixel *i* computed from the MSI data using the soft landmark contrast procedure above, and let *v*_*i*_ denote the corresponding edge strength in the visualization. We quantify their agreement using SpecEdge-Dice,

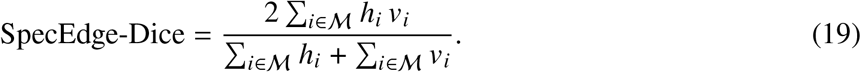

This metric quantifies whether transitions in the spectral reference coincide with transitions in the visualization while remaining less sensitive to variation within homogeneous regions.

#### Illustrative example

To illustrate why neighborhood preservation alone does not ensure visible tissue contrast, consider scalar spectral values *x*_*i*_ sampled along a spatial tissue line, with a transition between two regions. A rendering *f* (*x*_*i*_) = *αx*_*i*_+*c*, with 0 < *α* < 1, preserves all pairwise distance rankings because | *f* (*x*_*i*_)− *f* (*x* _*j*_ ) | = *α*|*x*_*i*_−*x* _*j*_ |. With the same tie-breaking rule, nearest-neighbor rankings and trustworthiness are unchanged, although displayed contrast is reduced.

For a conceptual gradient-based edge detector on a fixed common scale, let *h*_*i*_ = |*x*_*i*+1_ − *x*_*i*_ |/*A*, where *A* > 0 is chosen so that *h*_*i*_ ∈ [0, 1]. The rendered edge map is then *v*_*i*_ = *αh*_*i*_. For a nonzero binary reference edge map, SpecEdge-Dice becomes 2*α*/(1 + *α*) and decreases as contrast is compressed. This is an illustrative construction, not a simulation of the benchmark’s SoLaCE pipeline: independent edge-map normalization can remove a uniform scaling effect. It demonstrates that distance rankings alone cannot guarantee displayed contrast; sensitivity of the implemented edge score also depends on its extraction and normalization procedures. Supplementary Materials 2 gives the explicit calculation.

#### Hyperparameter tuning for visualization selection

Candidate pMiCS visualizations were generated by sequential hyperparameter tuning on a single source image. The final search used two practical stages. First, we performed coordinate one-factor-at-a-time exploration of categorical axes to identify useful regions of the search space. Second, we applied stochastic hill climbing, a local-search strategy, over neighboring configurations to refine candidate settings (*32*). At each step, the search sampled a nearby configuration, evaluated the scalar visualization objective, and retained changes that improved the current candidate while skipping duplicate configurations.

For trial *t* with hyperparameters ***θ***_*t*_, let *m*_*q,t*_ denote the score for metric *q* ∈ {edge, lum, chroma}, corresponding to SpecEdge-Dice, luminance RMS contrast, and LAB chroma entropy. Given user-specified bounds (*a*_*q*_, *b*_*q*_), each metric was clipped and scaled to [0, 1],

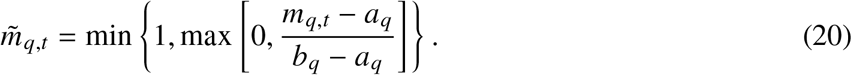

The scalar objective maximized during tuning weighted boundary fidelity twice as strongly as each color-variety metric:

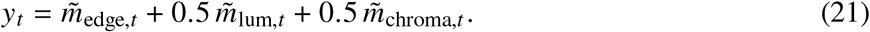

Thus SpecEdge-Dice received weight 1, while luminance RMS contrast and LAB chroma entropy each received weight 0.5. An alternative `rank composite` mode combines trial-wise ranks across the search history instead of the bounded weighted score above; it was used only when explicitly selected in the configuration. Duplicate configurations were skipped, and tuning stopped after a fixed trial budget *T*_max_.

#### Panel construction: local quality and selection strategies

Given the candidate pool, a gallery of *K* complementary views is assembled, and we compare strategies for selecting them. All scoring uses the three metrics that correlated positively with pathologists’ preferences (SpecEdge-Dice, luminance RMS contrast, and LAB chroma entropy), with weights (1, 0.5, 0.5) that assign boundary fidelity twice the weight of each color-variety metric. The geometric-fidelity metrics (trustworthiness and the global Spearman cosine) are excluded because they correlated *negatively* with pathologists’ preferences (Figure 7D).

##### Local quality maps

For each candidate rendering and its tissue mask, the three metrics are computed on overlapping patches (size 48, stride 16). To obtain pool-relative maps for *N* candidates, the *N* metric values at each tissue location are rank-normalized to [0, 1], giving candidate-specific rank maps 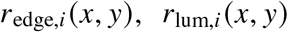, and 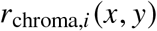, which are combined into a local-quality score

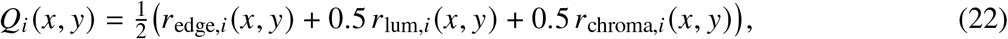

which measures the quality of candidate *i* at that location relative to the other candidates. Applying the same construction to the whole-image metrics gives candidate-specific ranks 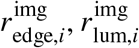, and 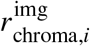 and the global composite

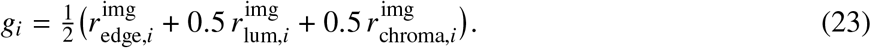

##### Selection strategies

We compare three strategies against a random baseline. All are seeded with the highest-*g*_*i*_ image and differ only in how the remaining views are added:

- **Greedy** adds the next-highest *g*_*i*_, with no interaction between choices.
- **Spatial** adds the view that most increases the running envelope *E ←* max(*E, Q*_*i*_)—the largest marginal local-quality gain relative to the current panel; it prefers views that are strong where the current panel is weak.
- **Coreset** embeds every candidate with DINOv2 (*33*) and, after the same initial selection, repeatedly adds the image farthest in embedding space from those already chosen. This *k*-center rule (*34*), introduced for core-set active learning, greedily minimizes the covering radius over the embeddings *z*,

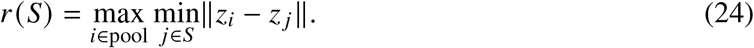

- **Random** draws views uniformly.

##### Metrics

Galleries are compared as a function of gallery size *k* using three measures:

- **Local envelope** (higher is better): the mean over tissue of the best local quality any selected view provides,

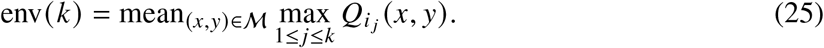

A location is covered when at least one selected view is strong there (the pixelwise maximum), so env(*k*) rises only when an added view improves locations that were previously weak.
- **Covering radius** *r* (*S)* (lower is better): how completely the selection represents the candidate pool in the DINOv2 embedding.
- **Worst image** min_*i*∈*S*_ *g*_*i*_ (higher is better): the quality of the weakest view in the panel.

Each measure coincides with one strategy’s objective by construction (**spatial** with the envelope, **coreset** with the covering radius, **greedy** with the worst image), so the measures are complementary lenses on a gallery rather than independent judges.

### Benchmark setup

We benchmarked visualization quality on eight curated MSI images and compared methods under identical preprocessing, repeated random seeds, and four blinded rankings by pathologists.

#### Images used

Table 1 lists the benchmark datasets with organ, spatial resolution, and source.

#### Preprocessing and visualization hyperparameters

Table 2 summarizes the visualization methods included in the benchmark.

**Table 2:** Visualization methods compared in the benchmark.

| Method | Type | Short description |
| --- | --- | --- |
| PCA | linear | principal-component projection to RGB |
| NMF | linear | nonnegative parts-based factorization |
| pUMAP | parametric nonlinear | neural UMAP mapping for out-of-sample projection |
| TOP3 | model-free baseline | three-channel $m/z$ selection from MSI-VISUAL (2) |
| pMiCS (parametric MiCS+LMC) | parametric nonlinear | distilled MiCS+LMC spectral-to-RGB mapping |

All MSI data were binned using an *m*/*z* bin width of 5 and normalized by total ion current (TIC). Tissue background was masked before visualization and rendered as black for all compared methods, so visual comparisons were restricted to valid tissue pixels.

For pMiCS, the parametric neural projection was implemented using the same parametric UMAP (pUMAP) projection framework used for the pUMAP baseline, but trained with the MiCS+LMC objective. pMiCS was trained for 30 epochs with coreset sampling to 1,000 landmarks. The MiCS clustering objectives were generated using 1,024, 2,048, and 4,096 clusters. Additional hyperparameters were set to *β* = 20, *k*_epoch_ = 20, batch size 1,024, temperature 100, and 10 warm-up epochs. Final RGB visualizations were generated using LAB-to-RGB color mapping. Runs used fixed random seeds, so a given configuration reproduces its output; where variability is reported, it reflects independent seeds and pixel subsamples. A one-at-a-time sensitivity analysis of these settings, rescored on the eight benchmark sections, is provided in Supplementary Figure S4.

For the pUMAP baseline, we used three embedding components, 50 neighbors, 5,000 randomly sampled pixels for visualization benchmarks, two graph-construction epochs, and two neural-network training epochs. These values were reduced from the default settings because default pUMAP training required more than five hours per image for smaller 20 *µ*m datasets and was not computationally feasible for repeated sweeps on larger MSI images in the benchmark workflow. Runtime measurements used 500 sampled pixels for pMiCS and pUMAP to compare practical train–inference behavior under a constrained interactive setting. These reduced pUMAP settings were chosen for feasibility and should not be interpreted as an exhaustive tuning of pUMAP. For PCA, the maximum number of iterations was set to 2,000.

### Metrics

Table 3 lists the primary metrics and what each metric checks.

**Table 3:** Metrics used for quantitative evaluation.

| Metric | What it checks |
| --- | --- |
| Trustworthiness | Preservation of local nearest-neighbor structure |
| Spearman correlation | Rank-order agreement of pairwise relationships |
| Luminance RMS contrast | No-reference light–dark separation over tissue pixels |
| LAB chroma entropy | No-reference chromatic diversity over tissue pixels |
| SpecEdge-Dice | Agreement of structural boundaries (edge fidelity) |

#### Ranking protocol

Pathologists independently ranked (*N* = 4) anonymized method outputs presented in randomized on-screen order. Each task showed the five methods (pMiCS, parametric UMAP, PCA, NMF, and TOP3) for one image, seed, and sampling condition, side by side; the pathologists force-ranked them from best to worst for pathology-oriented MSI review, with no ties or omissions. One seed was shown at a time. The planned design comprised ten seeds for each of eight images, or 80 sessions per sampling condition. Figure 7 summarizes only sessions with complete rankings from all pathologists: 75 random-sampling sessions and 79 superpixel-guided sessions. Mean ranks and correlations are averaged across pathologists; uncertainty in panel B is shown as the standard deviation across pathologists and sessions, and inter-pathologist agreement is summarized by Kendall’s *W*. Supplementary Figure S13 reports section-level inferential analyses of the same 154 sessions with complete rankings from all pathologists, treating pathologists and seeds as repeated measurements within sections.

#### Statistical analysis

The section-level inferential analysis in Supplementary Figure S13 uses the 154 sessions with complete rankings from pathologists. For each sampling mode, the tissue section is the independent unit; pathologists and seeds are repeated measurements within each section. Pairwise win totals are tested with a two-sided sign-flip permutation test: within each section, outcomes from all pathologists and seeds are reversed together, and all 256 section-level sign assignments are enumerated. P-values are Holm-adjusted across ten unique method pairs. Mean ranks are averaged over pathologists and seeds within each section and compared using exact one-sided Wilcoxon signed-rank tests of whether pMiCS has a lower rank than each baseline; these P-values are Holm-adjusted across four baselines.

### Dataset sources

Table 1 lists the MSI datasets displayed as biological images in the main and supplementary figures. Datasets were also used in quantitative benchmark plots, runtime analyses, or aggregate ranking and metric figures. Dataset numbers are internal identifiers used in this manuscript, and direct public dataset links are provided where available.

### Use of artificial intelligence-assisted tools

Artificial intelligence-assisted tools were used for software development and manuscript preparation. Cursor (Anysphere, Inc., CA, USA), including its Composer functionality, was used with OpenAI ChatGPT, SpaceXAI Grok, and Anthropic Claude to assist with code generation, refactoring, debugging, and documentation. PrismAI was used for writing assistance, language refinement, copyediting, proofreading, and LaTeX editing. The specific model versions available through these interfaces varied during the project and were not consistently exposed or retained by the software platforms. The AI tools did not independently define the study design, select datasets, perform the expert pathology assessment, determine the scientific conclusions, or generate experimental data. All AI-assisted code was reviewed and tested by the authors, and all AI-assisted text was critically reviewed, revised, and approved by the authors. The authors take full responsibility for the accuracy, originality, integrity, and conclusions of the submitted work. No AI-generated images were used as research data or manuscript figures.

### Ethical approval

Animal breeding and tissue harvesting for datasets D3 and D4 (Table 1) were approved by the Department of Comparative Medicine at Oslo University Hospital, Norway (approval identifier: IV2-2022).

The remaining human and animal MSI datasets were obtained from public repositories or previously published studies listed in Table 1. This study involved no new recruitment, intervention, or collection of human tissue and used no directly identifiable participant information. Ethical approval and consent for the source datasets were governed by the original studies and repositories.

## Acknowledgments

The authors thank Dr. Jorunn Stamnæs for performing the in-house MSI measurements of the mouse-brain samples (datasets D3, 4) and for technical discussions, Thomas Brüning for sectioning and preparing the mouse-brain hemispheres and control stains, Prof. Dr.-Ing. Marcin Grzegorzek, University of Lübeck, Dr. Henning Leske, and Christina Bang, Oslo University Hospital, for supporting this work.

The authors especially thank all providers of open-access datasets: Lars Gruber and Carsten Hopf (spinal cord dataset, Hochschule Mannheim, Germany), Miriam F. Rittel and Carsten Hopf (liver metastasis dataset (*35*), Hochschule Mannheim, Germany), Ethan Stancliffe (mouse-brain tumor dataset, Washington University in St. Louis, USA, (*36*)), Haruki Uchino 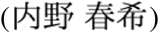 and Makoto Arita 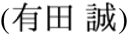 (mouse-brain dataset from the SMASH dual-polarity imaging study, RIKEN Center for Integrative Medical Sciences 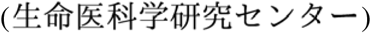, Yokohama, Japan, (*37*)), and Brittney Gorman (kidney dataset, Pacific Northwest National Laboratory, USA). These datasets were used for benchmarking, analysis, and figure preparation.

The mass spectrometry measurements of our mouse-brain samples were performed at the Proteomics Core Facility at the University of Oslo/Oslo University Hospital, which is supported by the Core Facilities Program of the South-Eastern Norway Regional Health Authority (HSØ) and NAPI (www.napi.uio.no; NFR, Norway; 295910).

## Author contributions

Conceptualization (J.G. and J.P.), Methodology (J.G. and J.P.), Software (J.G. and J.P.), Validation (J.G. and J.P.), Formal analysis (J.G. and J.P.), Investigation (J.G. and J.P.), Resources (J.P.), Data curation (J.P.), Writing–original draft (J.G. and J.P.), Writing–review and editing (J.G. and J.P.), Visualization (J.G. and J.P.), Supervision (J.P.), Project administration (J.P.), Funding acquisition (J.P.).

## Data, code, and materials availability

All publicly available MSI datasets are identified with repository records or accession numbers in Table 1.Analysis code will is made publicly available on Zenodo (https://doi.org/10.5281/zenodo.23372802). All other data needed to evaluate the conclusions are present in the main text or the Supplementary Materials. This study did not generate new materials.

## Supplementary Materials

File 1. Figures S1 to S20

File 2. Further explanations of equations.

## Supplementary Materials 1

### Supplementary Figures

#### Qualitative comparison of molecular edge detectors

SoLaCE is compared with two baseline edge detectors across diverse MSI tissues. The panels support visual assessment of boundary coherence, anatomical detail, and sensitivity to isolated high-intensity responses.

**Figure S1:**
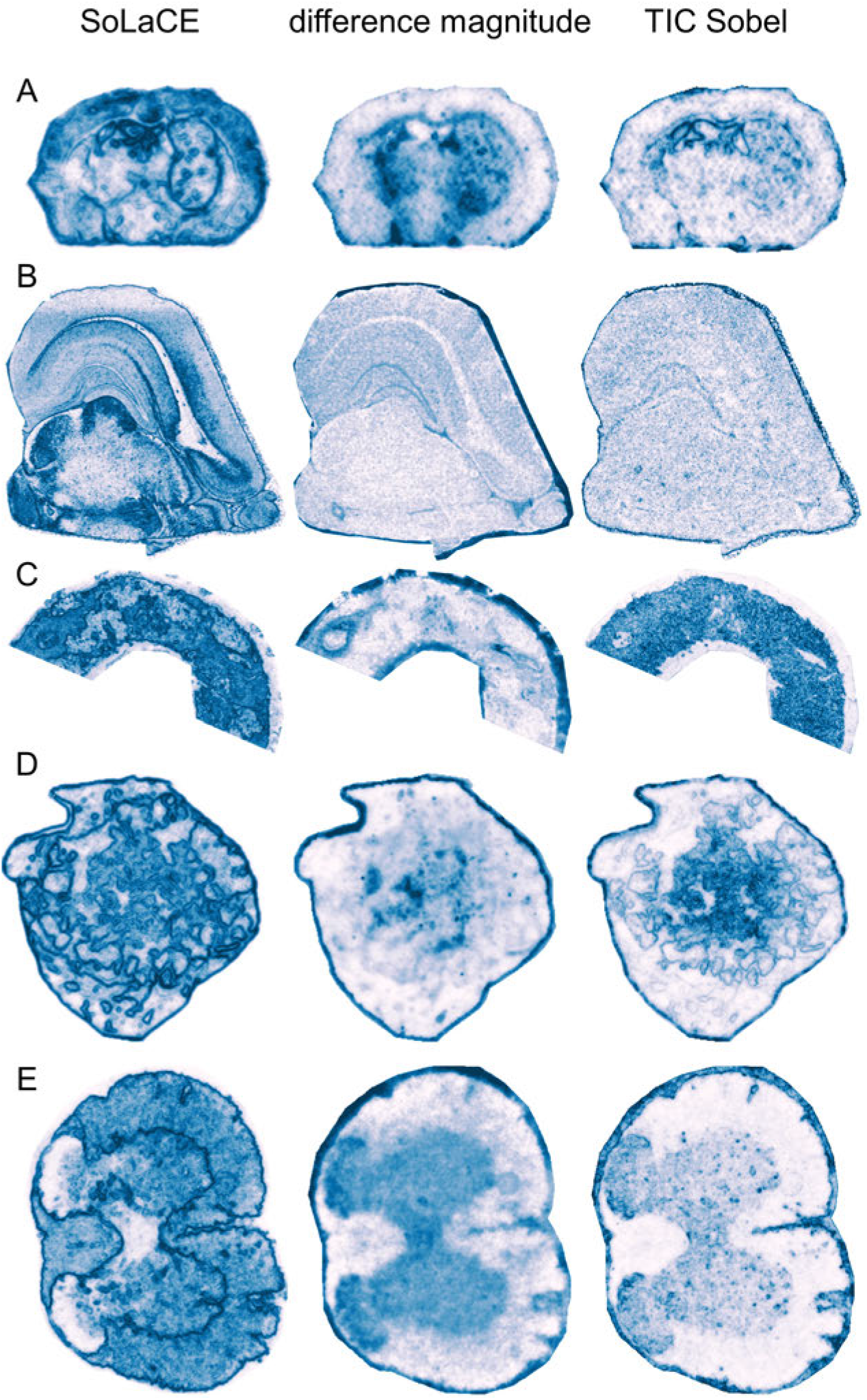
Qualitative comparison of SoLaCE with baseline edge detectors. (**A**–**E**) Side-by-side edge maps for five MSI sections generated using SoLaCE, difference magnitude, and TIC-Sobel. These panels constitute the image set included in the qualitative review by a single expert pathologist described in Materials and Methods.

**Figure S2:**
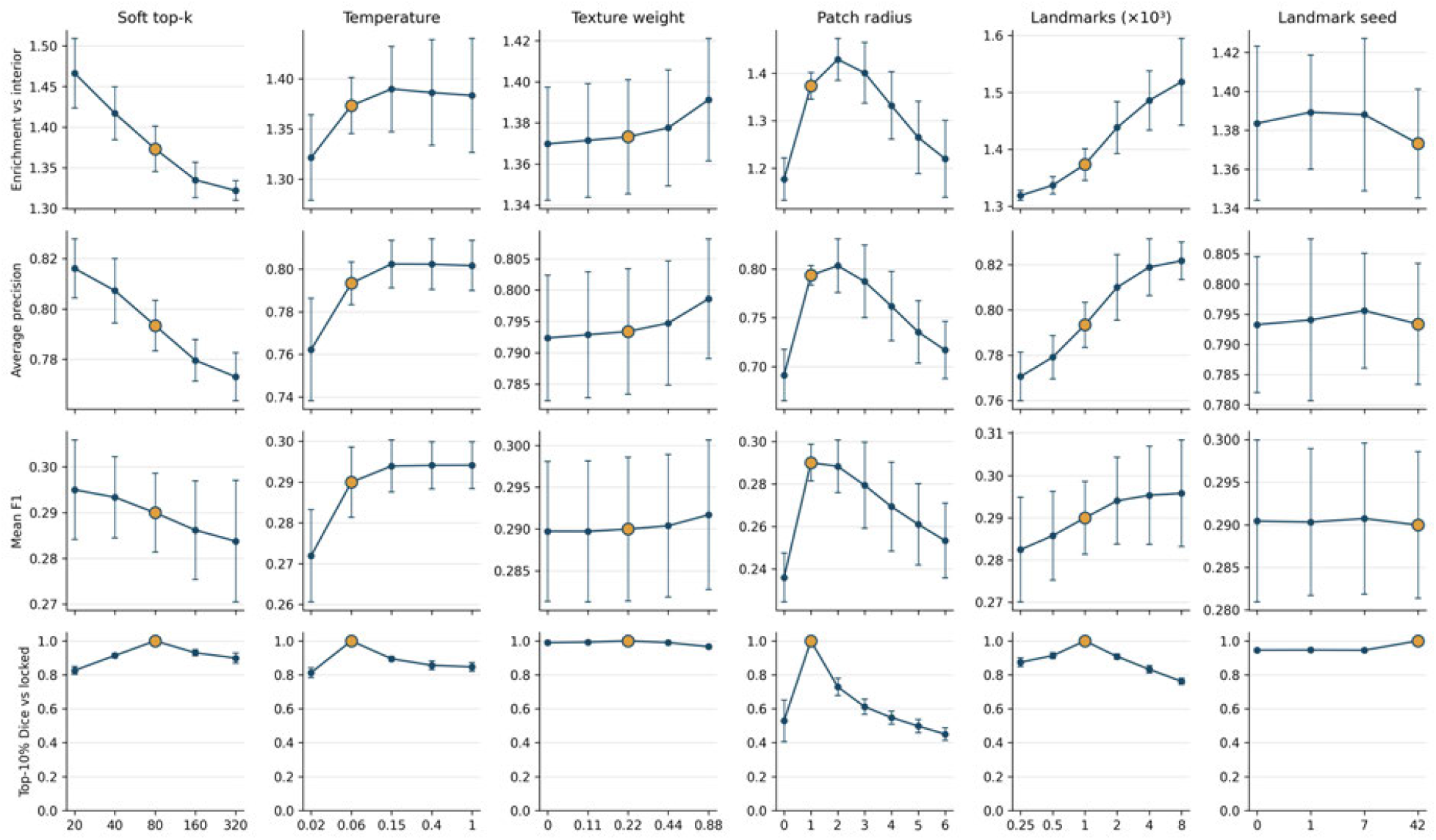
Sensitivity of SoLaCE to its hyperparameters. Each setting is varied one at a time while the others are held at the locked configuration (soft top-*k* = 80, temperature *τ* = 0.06, texture weight 0.22, patch radius 1, 1,000 landmarks, seed 42; gold marker). Performance is evaluated on four mouse-brain hemispheres against the same expert anatomical annotations used in the boundary benchmark. Top three rows: border-to-interior enrichment, average precision, and mean F1 (density-matched at the top 5–20% of tissue). Bottom row: top-10% Dice overlap between each resulting edge map and the locked map (rank-normalized over tissue), which quantifies agreement between edge maps as each setting changes (the locked point is 1 by definition). Each point is the mean across four sections; error bars show the standard deviation. Ranges: soft top-*k*, 20–320; temperature, 0.02–1; texture weight, 0–0.88; patch radius, 0–6; landmark count, 250–8,000; and landmark seed, 0, 1, 7, and 42. Performance varies smoothly and modestly with each setting; no choice causes a collapse except complete removal of spatial averaging (patch radius 0). The edge maps are highly self-consistent (top-10% Dice > 0.8), except with respect to patch radius, and are essentially unchanged across landmark seeds. The locked configuration is a mid-range choice rather than a tuned optimum: a smaller soft top-*k*, more landmarks, or a patch radius of 2 scores marginally higher on the anatomical metrics.

#### SoLaCE component ablation

The landmark step and its softmax control how sharply SoLaCE resolves fine structure; removing the landmark step and contrasting spectra directly reduces anatomical detail.

**Figure S3:**
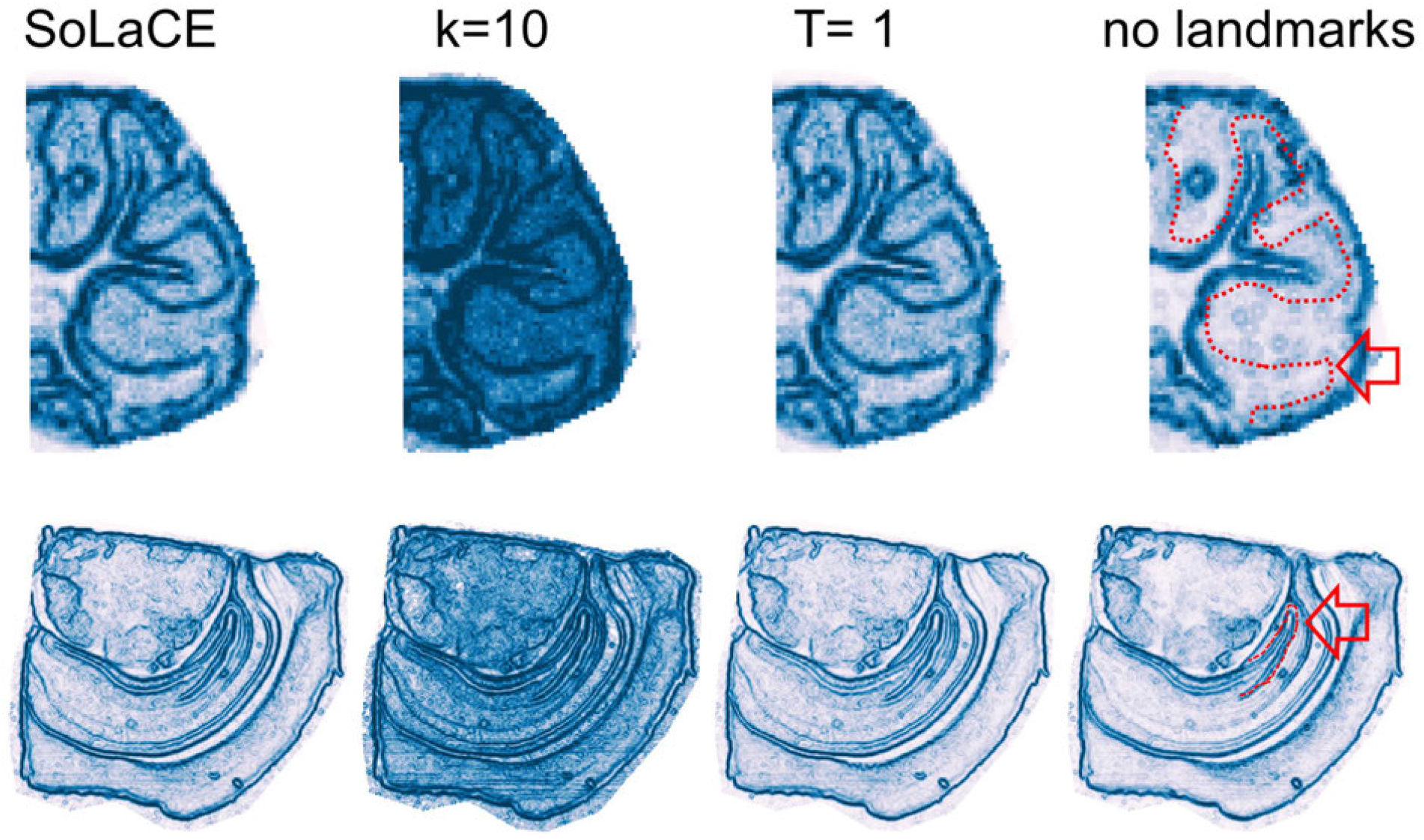
SoLaCE component ablation on two sections. The examples are a mouse-brain section from a SMASH dual-polarity imaging study (*37*) (first row) and one of the internal mouse-brain hemispheres (*1*) (second row). Columns show SoLaCE with the default softmax over the top *K* = 80 of the 1,000 landmark similarities; a smaller top-*K* (*K* = 10), which produces denser edges; a softmax temperature of *T* = 1; and complete removal of the landmark step, with spectra contrasted directly, which produces sparser boundaries. Red outlines and arrows in the *no landmarks* column mark anatomical structures lost without the landmark step. The top-*K* landmark signature and its softmax temperature together control the sharpness with which fine structure is resolved.

#### Sensitivity of pMiCS to its hyperparameters

Each pMiCS setting is varied one at a time and the resulting visualization is rescored on the eight benchmark sections, showing that the fixed configuration is a mid-range choice rather than one tuned to the benchmark.

**Figure S4:**
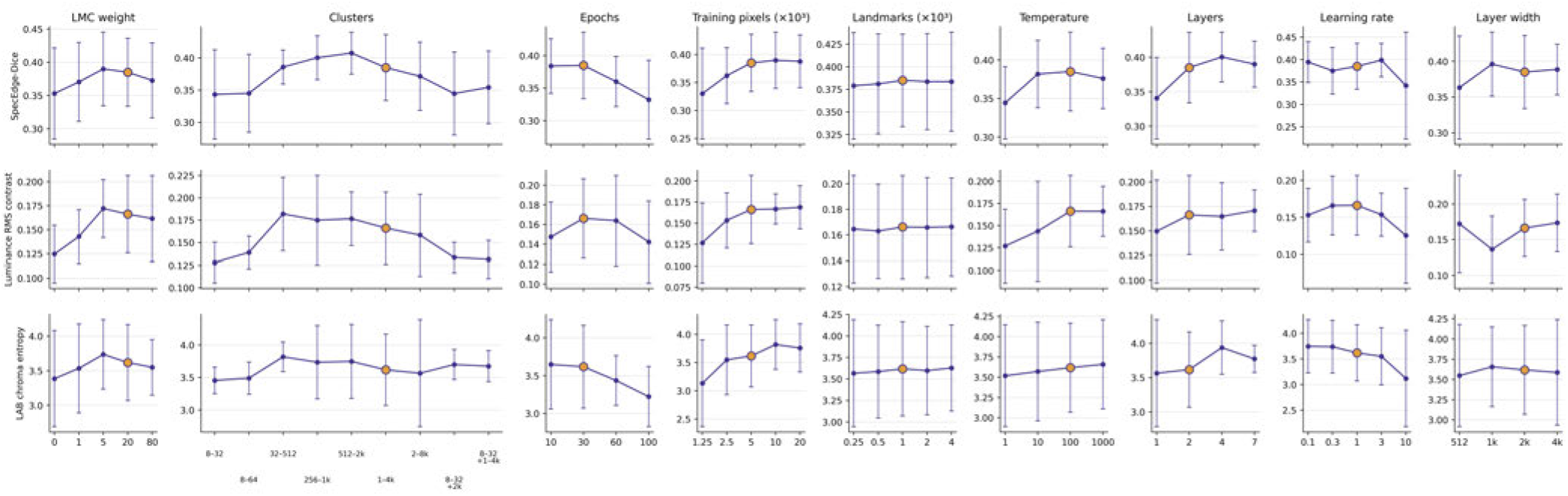
Sensitivity of pMiCS to its hyperparameters. Each setting is varied one at a time while the others are held at the locked configuration (LMC weight *β* = 20; MiCS cluster resolutions 1024/2048/4096; 30 epochs; 5,000 training pixels; 1,000 landmarks; temperature 100; 2 layers; learning rate 1; layer width 2048; gold marker), and each resulting visualization is rescored on the same eight benchmark sections. Rows report the three pathologist-aligned metrics: SpecEdge-Dice, luminance RMS contrast, and LAB chroma entropy. Each point is the mean over the eight sections and error bars are the standard deviation. The Clusters axis varies the set of *k*-means cluster counts (heads) combined by MiCS, not a single clustering with an intermediate *k*: a label such as “8–32” denotes three separate fits, at 8, 16, and 32 clusters, with the label showing only the smallest and largest head (8– 32= {8, 16, 32}; 8–64= {8, 16, 32, 64}; 32–512= {32, 64, 128, 256, 512}; 256–1k= {256, 512, 1024} ; 512–2k= {512, 1024, 2048}; 1–4k= {1024, 2048, 4096}, locked; 2–8k= {2048, 4096, 8192}; “8–32+2k”= {8, 16, 32, 2048}; “8–32 1k–4k”= {8, 16, 32, 1024, 4096}). Performance varies smoothly and modestly with each setting, and the locked configuration is a mid-range choice rather than one selected to maximize any metric.

**Figure S5:**
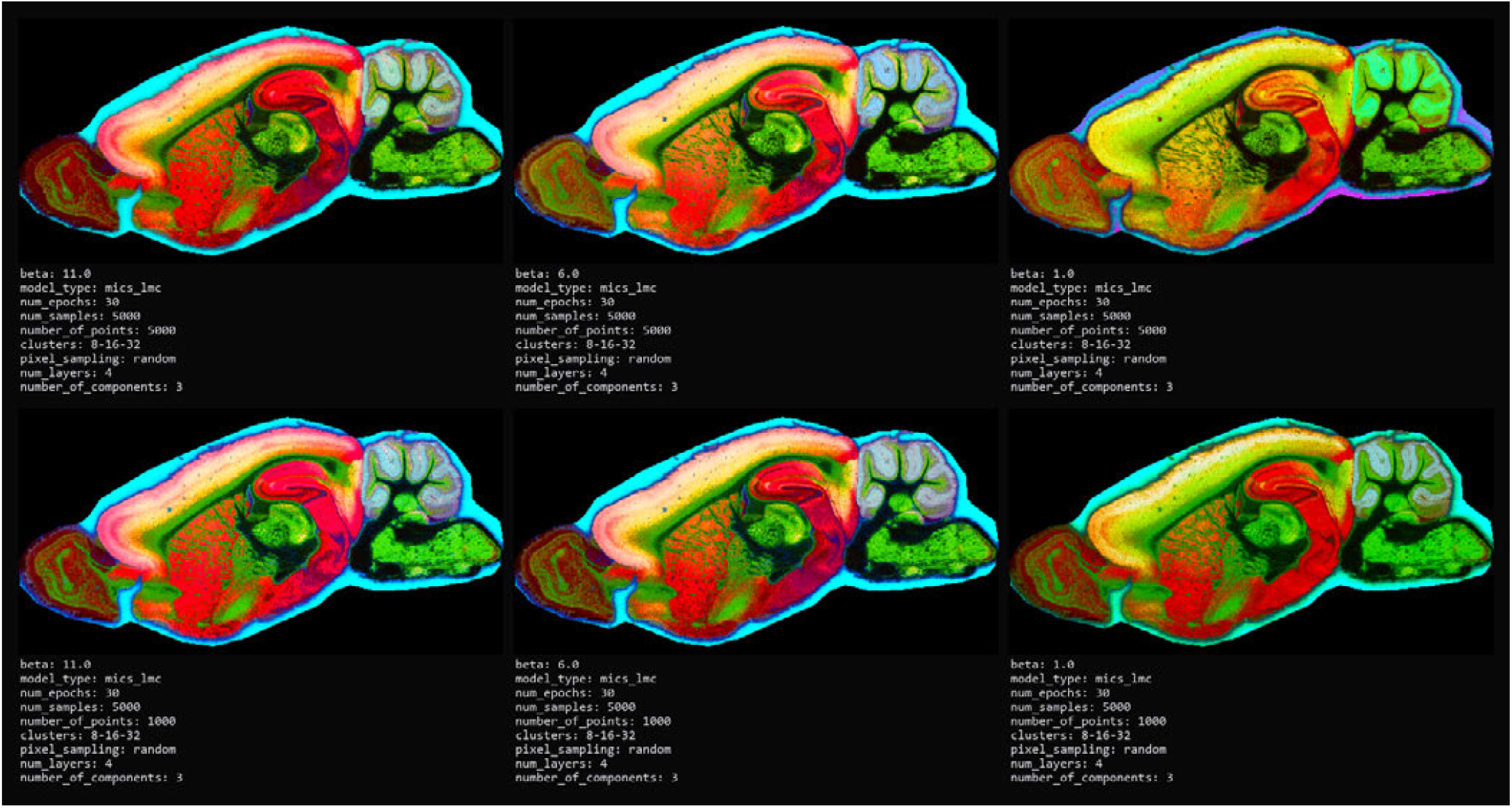
Qualitative sensitivity of pMiCS to *β* and the number of points. Columns show *β* = 11, 6, and 1 from left to right; rows show `number of points`=5,000 (top) and 1,000 (bottom), while `num samples` remains 5,000 in all panels. In these examples, higher *β* appears to increase local contrast in the cerebellar region while reducing the prominence of some hippocampal laminar and connecting patterns. These are qualitative rendering differences; possible Purkinje-cell-associated structures are not identified or validated by this figure. Dataset from (*37*).

#### Gallery construction strategies

Supplementary Figure S6 extends the two metastasis examples in Figure 8 to five MSI sections, comparing greedy, spatial, and coreset selection with a random baseline. Greedy selection prioritizes whole-image quality, spatial selection adds views that improve the local-quality envelope, and coreset selection prioritizes coverage of the candidate pool in DINOv2 embedding space. The local envelope, covering radius, and weakest included whole-image score quantify these complementary objectives, as defined in the Panel construction subsection of Materials and Methods.

The selected views illustrate why panel size alone does not establish complementarity. Greedy selection can include near-duplicate high-scoring views, whereas coreset selection can include near-uniform outliers that add embedding diversity but little tissue detail. Spatial selection prioritizes local complementarity by improving locations that the existing panel represents weakly. Because each evaluation measure corresponds to a selection objective, these comparisons characterize panel construction rather than provide independent biological or diagnostic validation.

**Figure S6:**
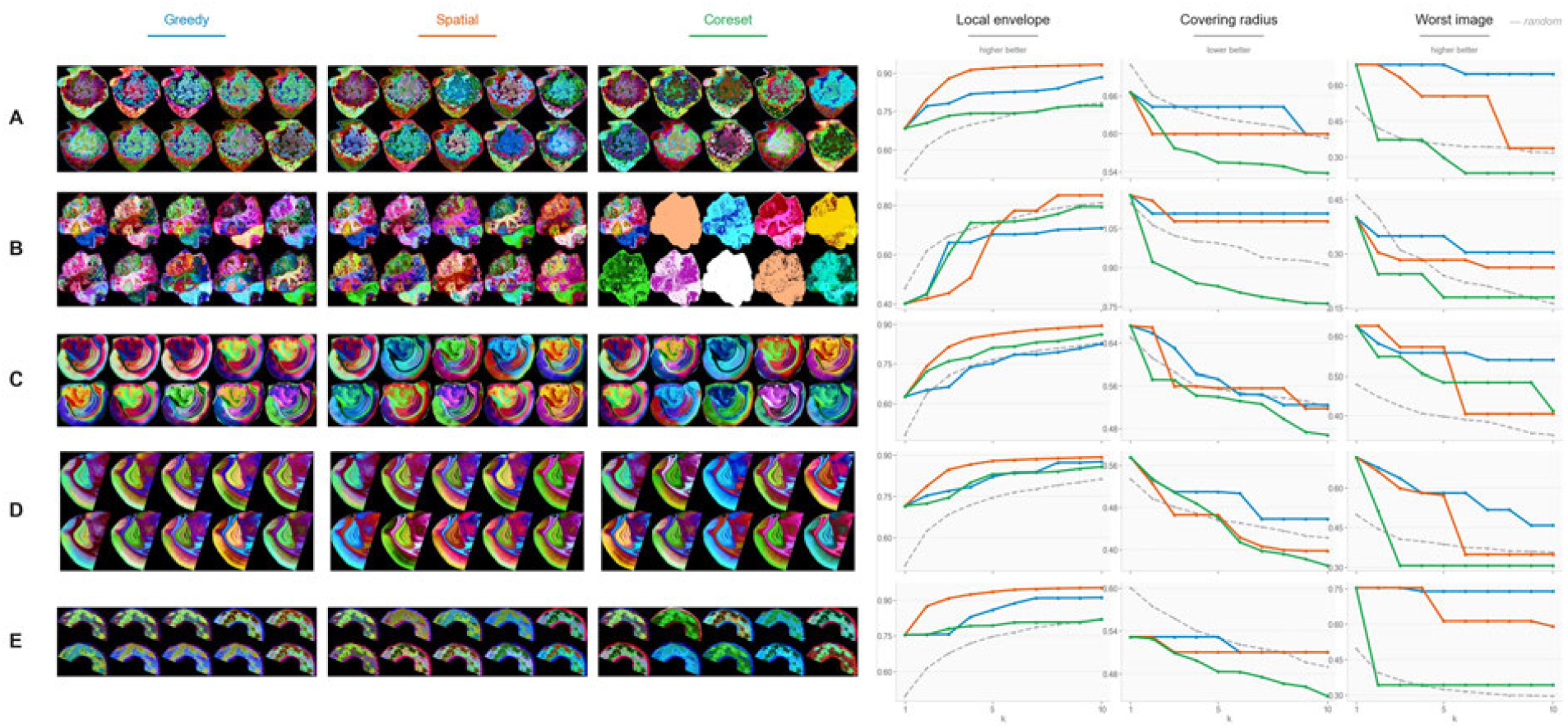
Comparison of gallery-construction strategies across five MSI sections (rows A–E). Left three columns: the ten selected pMiCS views for the greedy (blue), spatial (orange), and coreset (green) strategies. Right three columns: the local envelope (higher is better), covering radius (lower is better), and worst image (higher is better) as functions of gallery size *k*, for the three strategies and a random baseline (gray dashed). Each strategy leads on the measure it targets by construction, and the structured strategies generally improve on random selection on those respective measures. Greedy selects near-duplicate views among its first three (rows C–E), whereas coreset selects near-uniform outlier views (row B); spatial selection favors marginal local-quality gains but does not lead on every measure or at every gallery size.

#### Concept-defining *m*/*z*

For each of the four concepts shown as ion images in Figure 5B, Table S1 lists the full ordered set of defining *m*/*z*.

**Table S1:** Concept-defining *m/z* for the concepts in Figure 5B. For each concept, the table lists the *m/z* values retained as spatially enriched (enrichment ≥1) from the 50%-loading candidate set, in descending order of enrichment. No molecular identities are assigned.

| Concept | Region | $m/z$ (enrichment order) |
| --- | --- | --- |
| c50 | normal liver | 844.6, 1604.2, 1539.2, 1607.2, 1606.2, 845.6, 1538.2, 1564.2, 1581.2, 1580.2, 1565.2, 1611.2, 781.6, 1613.2, 1566.2, 1567.2, 1609.2, 1583.2, 1612.2, 797.6, 1582.2, 1555.0, 1608.2, 1610.2, 1592.2, 1540.2, 738.4, 1595.2, 1541.2, 1554.0, 821.6, 1594.2, 1568.2, 1596.2, 1597.2, 1557.2, 1587.2, 1584.2, 1569.2, 1556.2, 1519.2, 1585.2, 1586.2, 1512.2, 1547.2, 749.6, 766.4, 763.4, 847.6, 1518.2, 759.6, 1520.2, 1546.2, 1500.2, 1513.2, 1570.2, 1571.2, 1545.2, 1542.2, 1572.2, 785.6, 834.6, 1544.2, 688.4, 1558.2, 1514.2, 761.6, 1516.2, 1528.2, 1543.2 |
| c20 | cellular centers (metastasis) | 798.6, 739.4, 827.6, 826.6, 810.6, 824.6 |
| c8 | connective-tissue matrix (metastasis) | 713.4, 742.6, 725.6, 726.6, 682.4, 741.6, 743.6, 683.4, 848.6, 849.6, 851.6, 810.6, 823.6, 827.6, 826.6 |
| c65 | proliferative glandular centers (metastasis) | 799.6, 826.6 |

#### Concept-deletion faithfulness

This test evaluates whether the concepts identified by the optimization are more relevant to the visualization than arbitrary or abundance-ranked concepts, thereby assessing model relevance rather than mere spectral presence.

**Figure S7:**
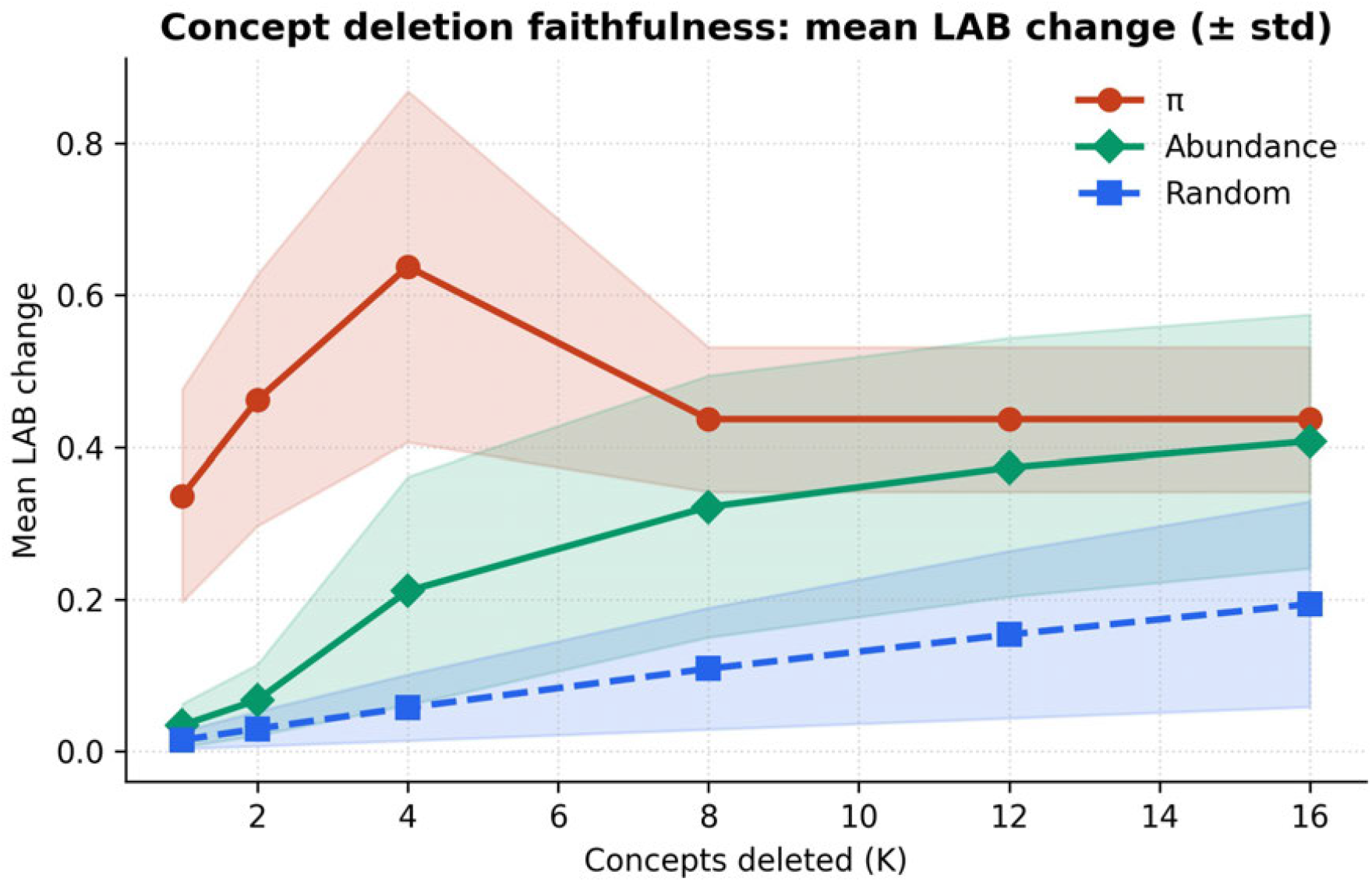
The concepts identified by the optimization drive the visualization. Mean LAB color change when the top-*K* concepts are deleted and the mixture re-optimized on the remainder, ranked by the optimized usage (*π*), by NMF abundance, and at random (bands show±one standard deviation). The optimization ranking disrupts the rendering more than both baselines at every *K*: more than random throughout, and more than abundance with the largest gap at small *K*, narrowing as *K* grows and nearly closing by *K* = 16. The small-*K* regime, where a few concepts carry most of the color, is the most informative.

#### Reconstruction fidelity of the concept representation

The sparse concept mixture reproduces the model’s rendered color closely even where the shared NMF basis reconstructs the full spectrum less tightly, because the pMiCS output depends only on the spectral variation the model uses.

**Figure S8:**
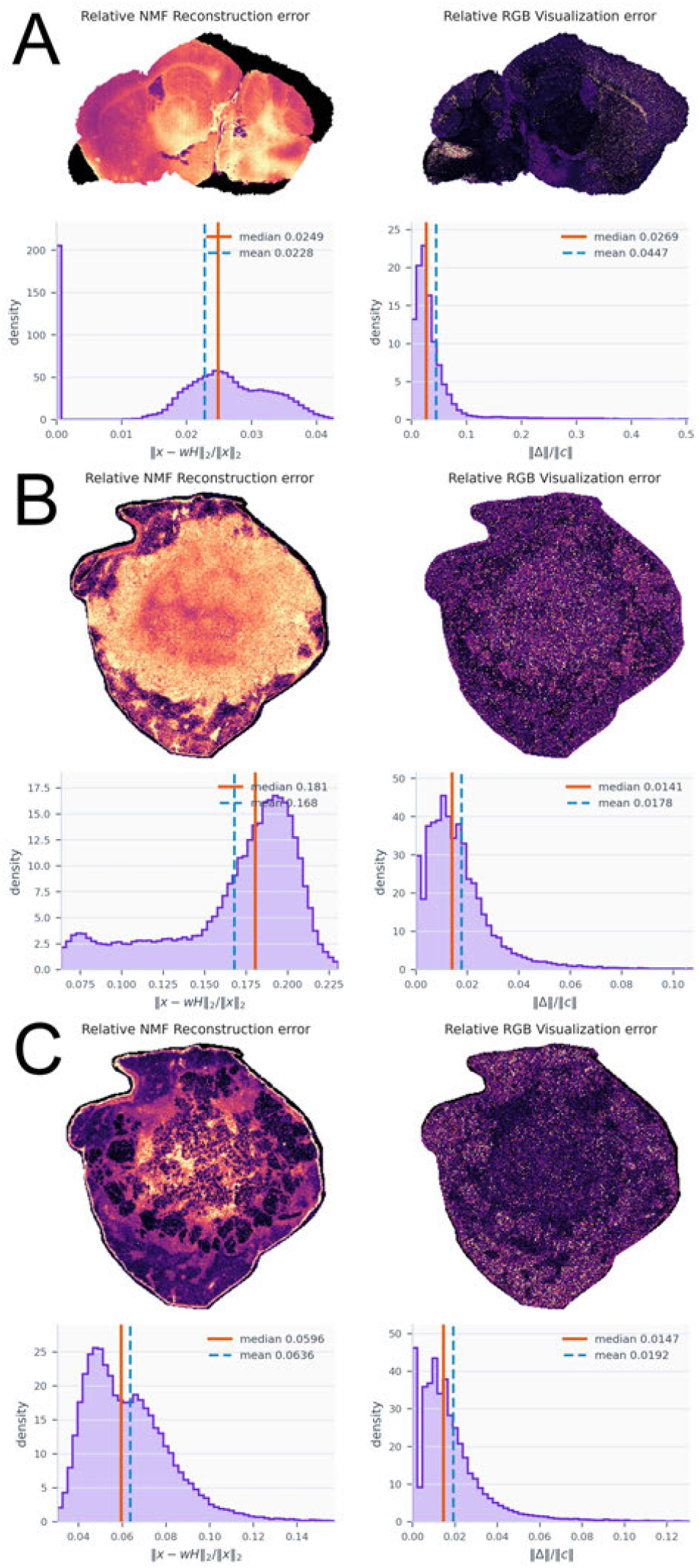
Reconstruction fidelity of the concept representation. *Left:* relative NMF spectral reconstruction error per pixel, ∥*x* − *WH*∥_2_ ∥*x*∥_2_ (spatial map and histogram). *Right:* relative error of the rendered CIELAB color, ∥Δ∥*/*∥*c*∥. Median errors (spectral / color): (**A**) mouse brain 0.025/0.027; (**B**) colorectal metastasis 0.181/0.014; (**C**) colorectal metastasis 0.060/0.015. The rendered color is reproduced to within a few percent throughout, even where the shared NMF basis reconstructs the full spectrum less accurately, because the output depends only on the spectral variation used by the model.

#### Usage–abundance discrepancy and pixel mixedness

The discrepancy *d*_*p*_ (Jensen–Shannon divergence between model usage *π*_*p*_ and the scale-adjusted molecular abundance 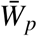) tracks where the rendering reweights the molecular signal, and is related to, but not reducible to, how mixed the pixel is.

**Figure S9:**
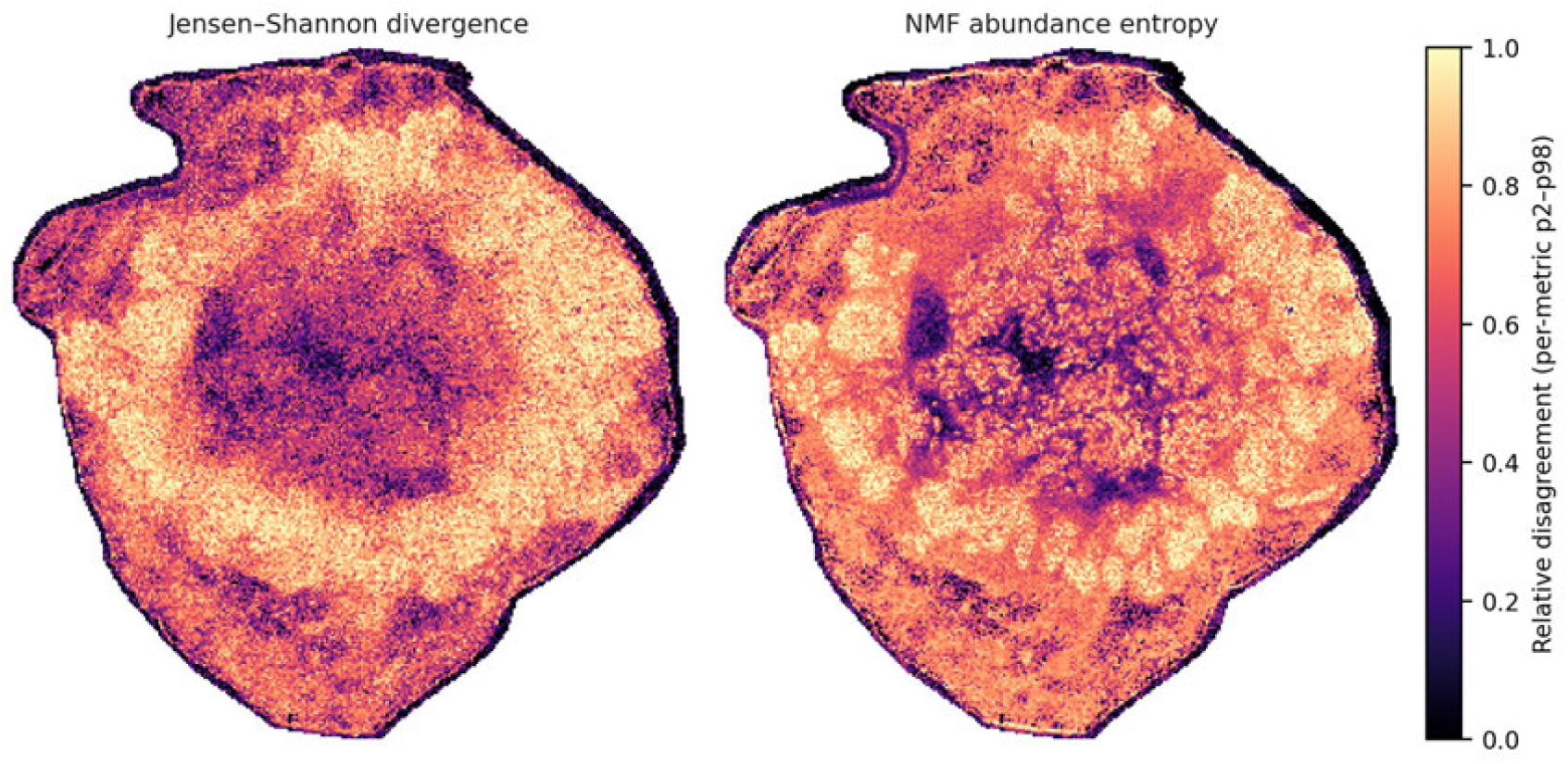
Usage–abundance discrepancy *d*_*p*_ compared with mixedness. Maps for a colorectal-adenocarcinoma liver metastasis. Left: the per-pixel Jensen–Shannon divergence *d*_*p*_ between the model’s concept usage *π*_*p*_ and the scale-adjusted NMF concept contribution 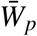 . Right: the Shannon entropy 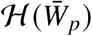 of the molecular mixture (both on a relative, percentile-normalized scale). The two maps share broad structure—more mixed regions tend to have higher discrepancy—but differ in detail, so *d*_*p*_ tracks mixedness only partly and is not a second entropy map.

**Figure S10:**
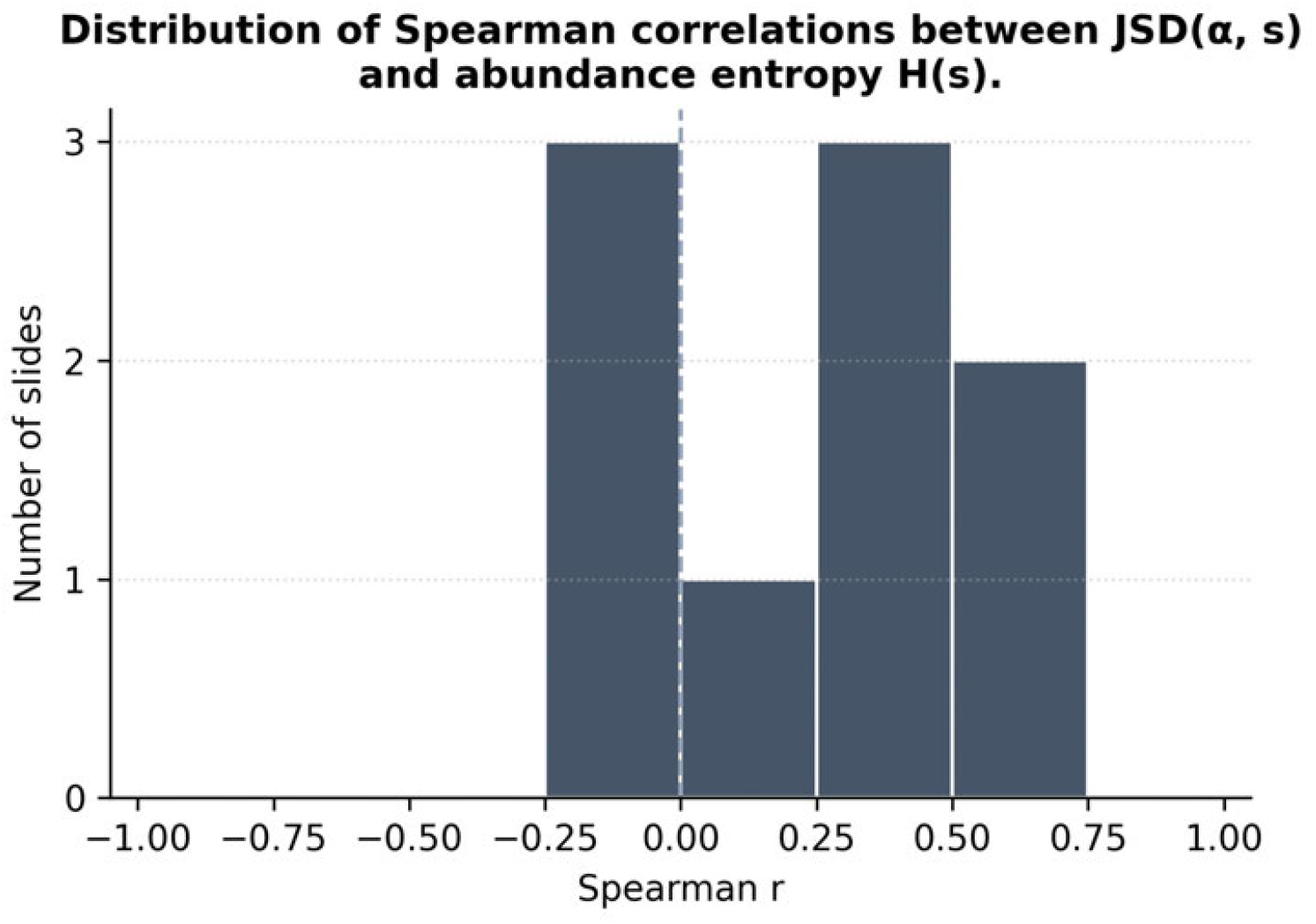
*d*_*p*_ correlates with, but is not reducible to, pixel mixedness. Distribution across sections of the Spearman correlation between the discrepancy *d*_*p*_ and the abundance entropy 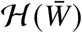. The correlations are positive but moderate (median 0.255) and are negative in a minority of sections; thus, more mixed pixels tend to have higher discrepancy, but *d*_*p*_ does not reduce to an entropy map.

#### Boundary and color metrics are necessary but not sufficient

Two controlled interventions show that the expert-aligned boundary/color metrics and the geometric structure-preservation metrics measure distinct, sometimes competing axes.

**Figure S11:**
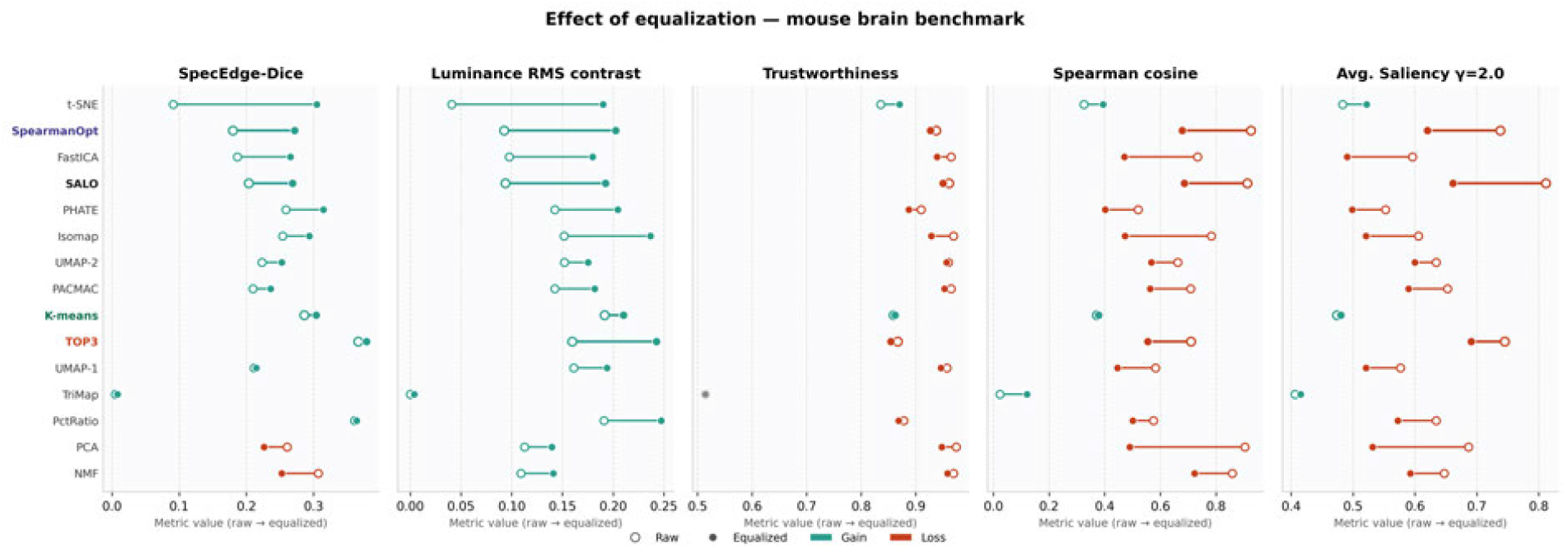
Histogram equalization improves contrast- and edge-based metrics while degrading geometric ones. Global per-channel histogram equalization was applied to each RGB visualization across the four mouse-brain hemispheres. For each method, each metric is shown before (open circle, raw) and after (filled circle, equalized); teal indicates a gain and red a loss. Equalization raises SpecEdge-Dice and luminance RMS contrast for almost all methods, while the structure-preservation metrics—trustworthiness, Spearman cosine, and MSI-VISUAL average image saliency (*2*)—all decrease. Thus, the operation that improves the contrast and edge metrics tends to reduce the geometric metrics, indicating that these are distinct and sometimes competing axes.

**Figure S12:**
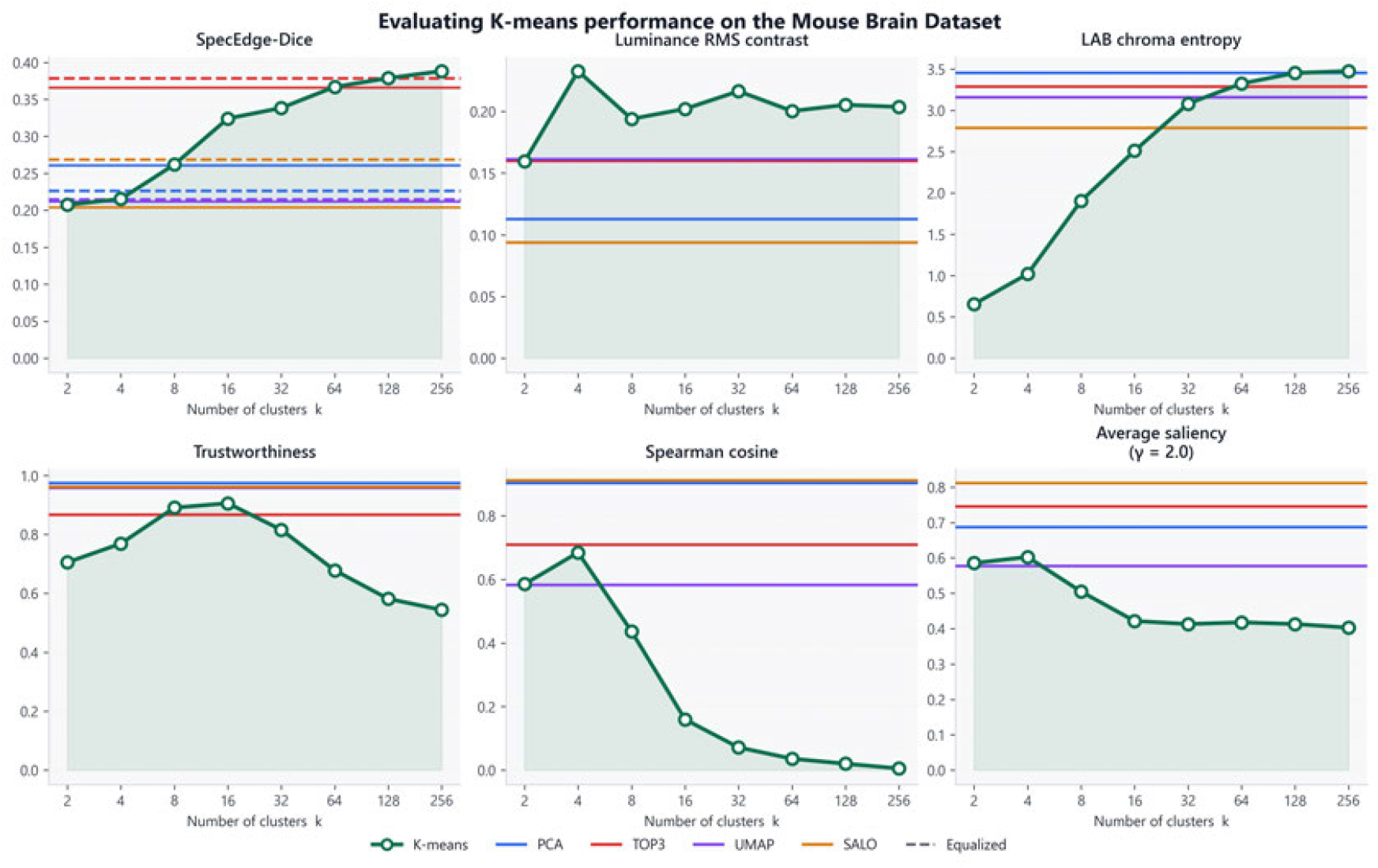
Boundary and color metrics are necessary but not sufficient. *k*-means segmentation is rendered as a visualization (one arbitrary RGB color per cluster) on the mouse-brain benchmark and evaluated across the number of clusters *k* (green), with dimensionality-reduction baselines shown as horizontal lines. As *k* increases, SpecEdge-Dice and LAB chroma entropy rise above all baselines, while the Spearman cosine correlation approaches zero and average saliency declines: the segmentation gains boundaries and color variety but loses information about region similarity. High boundary- and color-metric scores can therefore coexist with a visualization that is uninformative about molecular structure, which is why the metrics must be applied jointly.

#### Section-level inferential analysis of pathologist rankings

The four-pathologist comparisons are assessed separately by sampling mode, treating each tissue section (not each seed or pathologist) as the independent unit.

**Figure S13:**
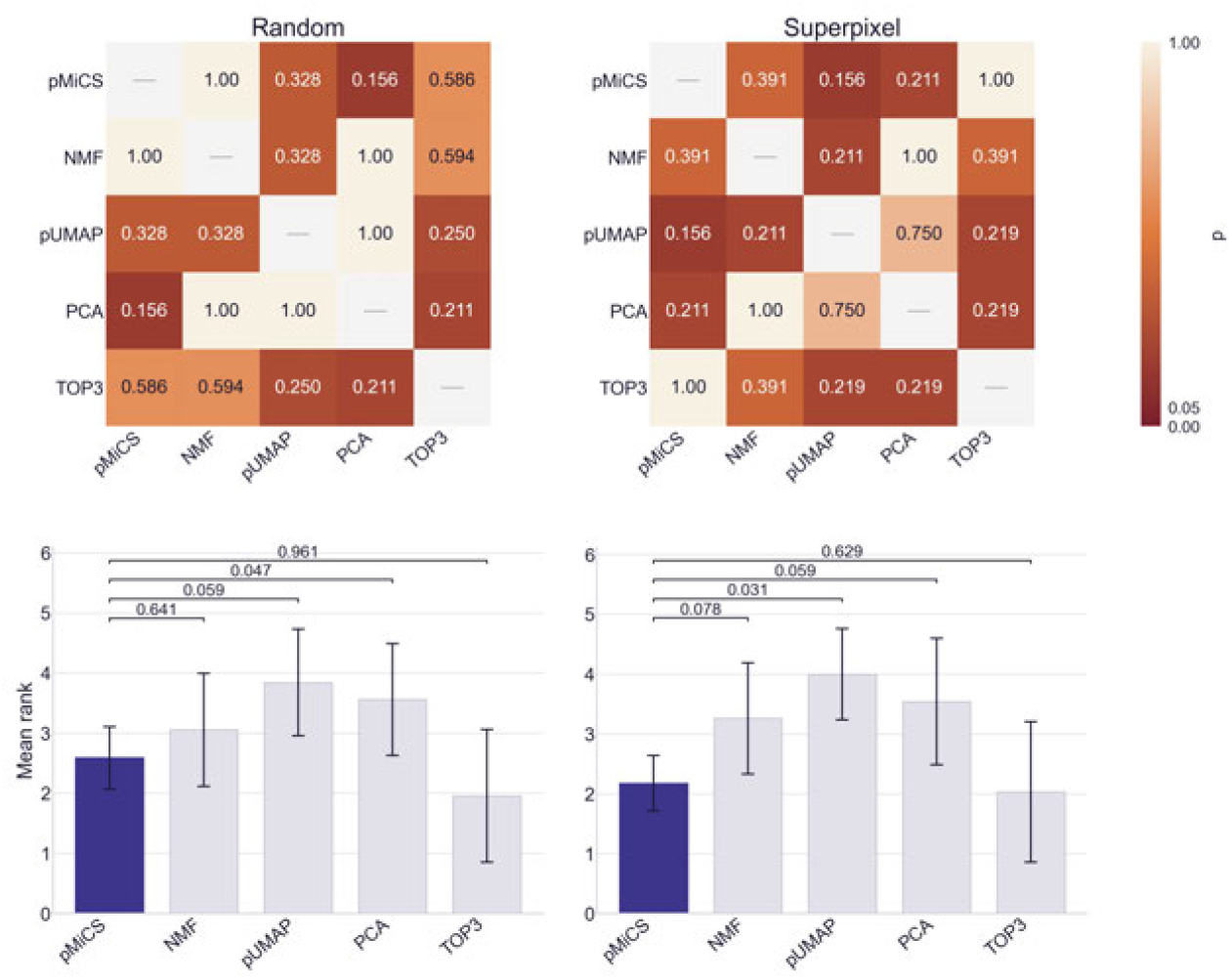
Statistical significance of the four-pathologist ranking analysis. Ranks from four independent pathologists are analyzed separately for random and superpixel sampling, using the 154 seed-level sessions ranked completely by the pathologists; the seeds and pathologists on a section are treated as repeated measurements for that section. *Top:* pairwise significance of the head-to-head win totals; each cell shows the Holm-adjusted *p*-value (color-coded) from a two-sided sign-flip permutation test (*38*). A win is a pathologist’s seed-level ranking in which the row method received a better rank than the column method, aggregated across the eight sections. Within each section, all pathologist and seed outcomes are reversed together; all 28 = 256 sign assignments of the eight sections are enumerated, and *p* is the fraction of assignments whose win total is at least as far from an even split as the observed total. Holm adjustment (*39*) is applied across the ten unique method pairs, so both directions of a pair share a *p*-value. With eight sections, the smallest attainable two-sided *p* is 2/256, which Holm adjustment over ten pairs raises to 0.078; no win-count test reaches Holm *p* < 0.05, and the smallest values are for pMiCS versus PCA (random, *p* = 0.156) and pMiCS versus pUMAP (superpixel, *p* = 0.156). *Bottom:* mean-rank bars; ranks are averaged over pathologists and seeds within each section, and each bar is the mean of the eight section means, with whiskers showing the standard deviation. Each bracket represents an exact one-sided Wilcoxon signed-rank test (*40*) of whether pMiCS has a lower section-mean rank than the corresponding baseline, Holm-adjusted across the four baselines. pMiCS has a significantly lower mean rank than PCA under random sampling (section-mean ranks 2.59 versus 3.56, Holm *p* = 0.047) and than pUMAP under superpixel sampling (2.17 versus 4.00, *p* = 0.031). The remaining comparisons with PCA, pUMAP, and NMF point in the same direction (superpixel PCA 3.54, *p* = 0.059; random pUMAP 3.84, *p* = 0.059; superpixel NMF 3.26, *p* = 0.078; random NMF 3.06, *p* = 0.641). TOP3 has a slightly lower mean rank than pMiCS (1.95 versus 2.59 random, 2.03 versus 2.17 superpixel); the difference is not significant in a two-sided test (*p* = 0.109 and *p* = 0.844). With eight sections as the independent unit, consistent directional differences may not reach the adjusted significance threshold.

#### Out-of-sample projection across brain sections

This analysis evaluates whether parametric mappings trained on sampled pixels can be reused for full-image and cross-image projection. Comparisons with pUMAP and PCA show both scalability and the conditions required for consistent color semantics.

**Figure S14:**
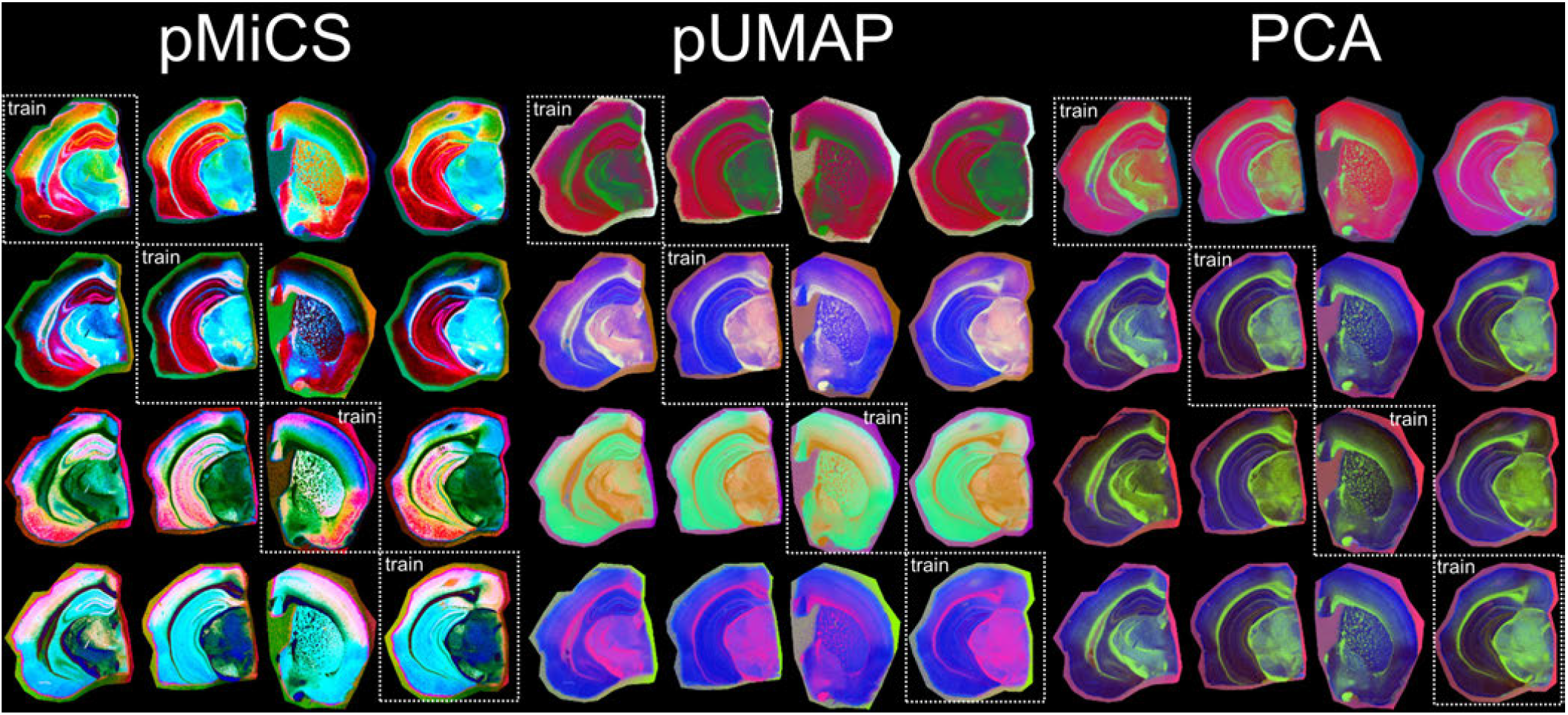
Out-of-sample projection and cross-image consistency of parametric MSI visualizations. The figure compares pMiCS, pUMAP, and PCA across four in-house mouse-brain hemisphere MSI datasets acquired at 20 *µ*m spatial resolution. For each row, pMiCS and pUMAP were trained on 5,000 valid pixels from the source image indicated by the dotted box, corresponding on average to approximately 7% of valid pixels. The trained models were then used to project the remaining pixels from the source image and all valid pixels from the other three images without re-optimizing the embedding. PCA is shown as a linear reference projection. Within each row, corresponding colors across the projected images illustrate reuse of the learned parametric mapping for scalable full-image rendering and cross-image comparison when preprocessing and feature alignment are matched.

#### Runtime of parametric visualization methods

Training and inference times quantify the computational requirements of the compared parametric workflows. The reduced pUMAP settings are reported explicitly to keep the practical comparison transparent.

**Figure S15:**
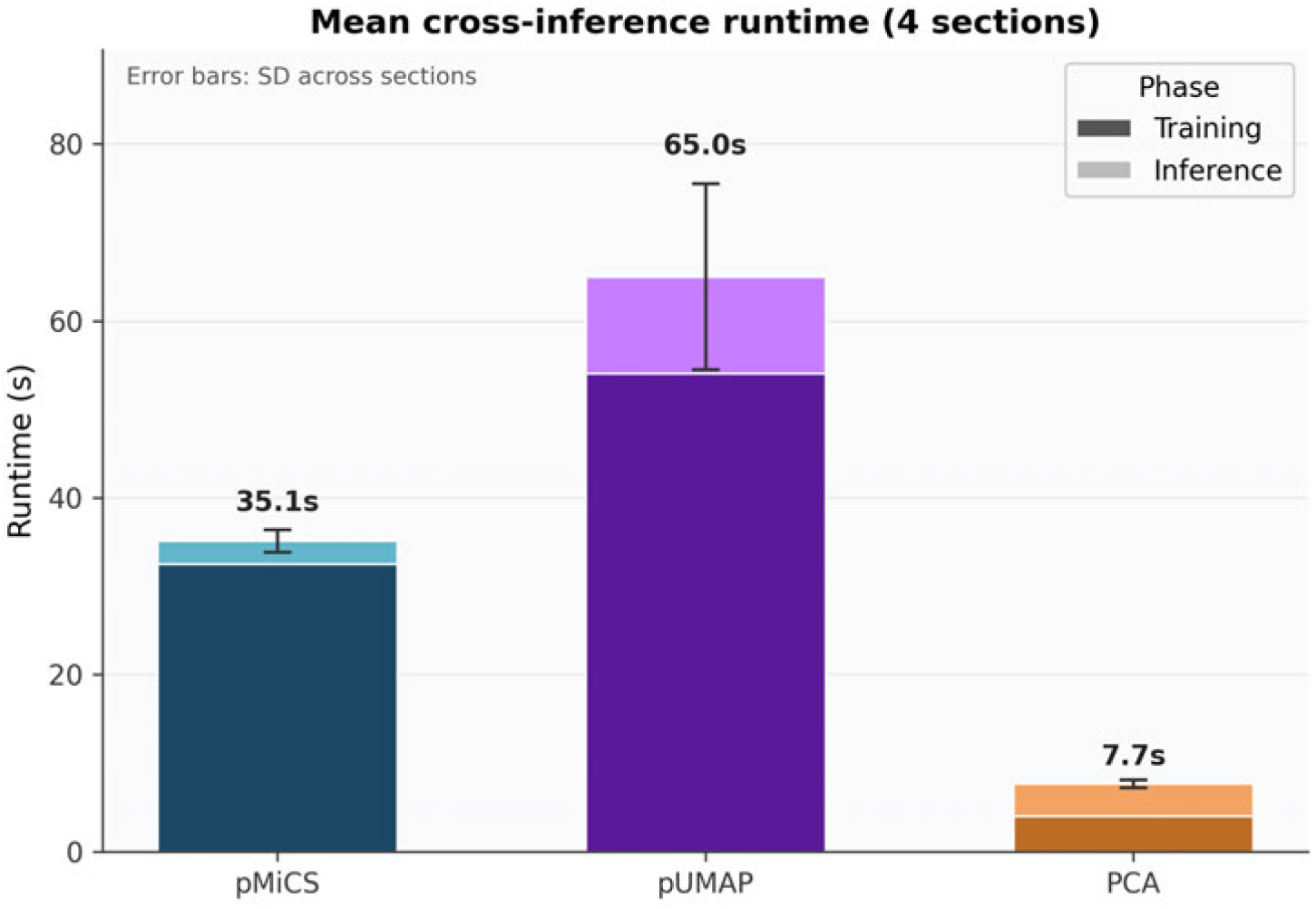
Runtime comparison for MSI visualization methods on brain images. Average training and inference times are shown for pMiCS, pUMAP, and PCA on the brain benchmark dataset. pMiCS and pUMAP were trained on 500 sampled pixels and run on GPU; PCA was run on CPU using the scikit-learn implementation. pMiCS achieved the lowest average training and inference time under these practical benchmark settings. pUMAP was evaluated with reduced settings because default pUMAP training required approximately five hours per image and was not practical for repeated visualization sweeps; therefore, this comparison should not be interpreted as an exhaustive optimization of pUMAP runtime.

#### Additional concept-based interpretation example

This example extends concept-based interpretation to a human kidney needle biopsy. Selected image regions are linked to sparse spectral concepts to distinguish molecular abundance from the concepts used by pMiCS to generate visual contrast.

**Figure S16:**
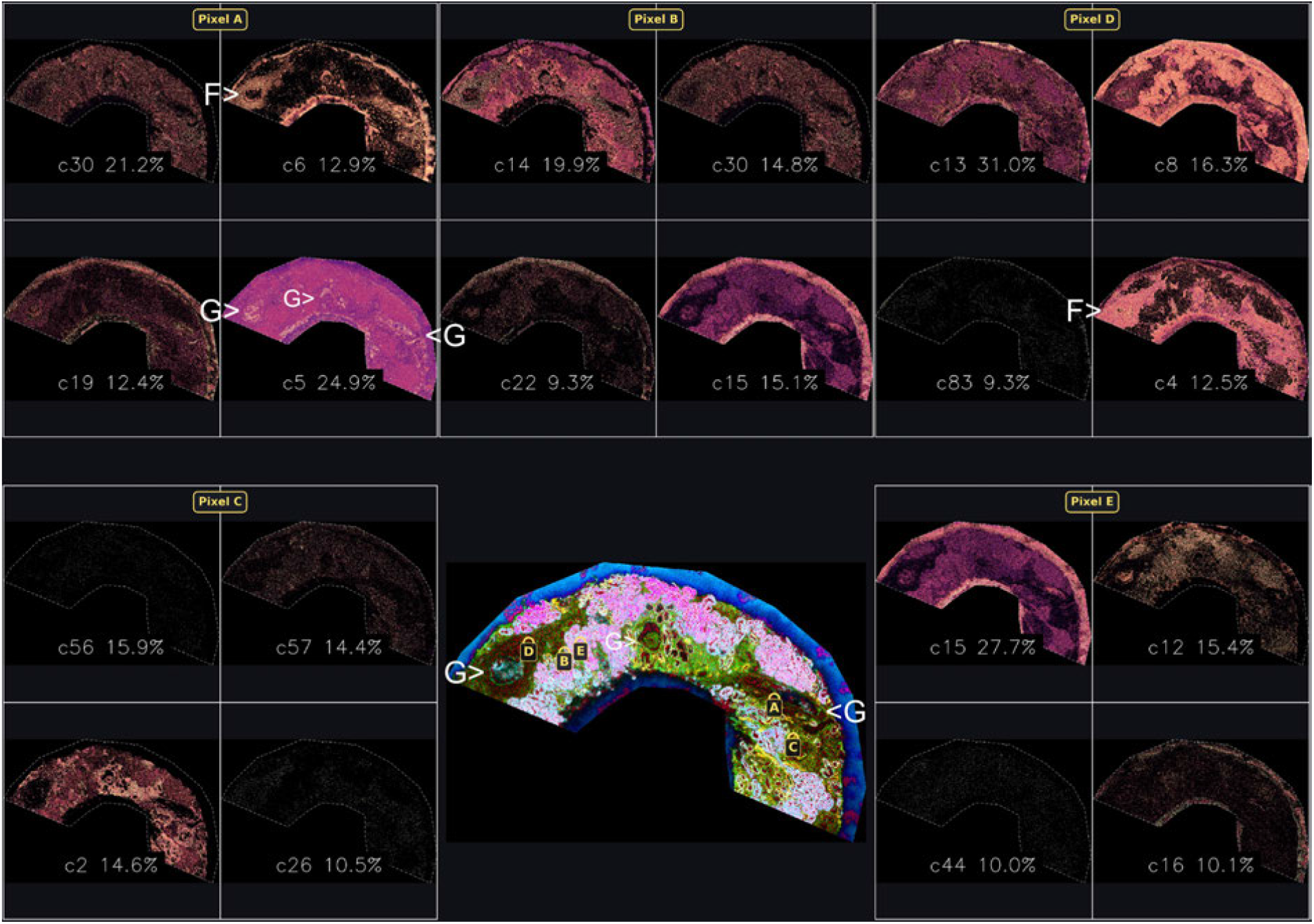
Concept-based interpretation of a pMiCS visualization of a human kidney needle biopsy with severe glomerular and interstitial pathology. The central panel is a pMiCS rendering of the biopsy (three glomeruli, G); pixels A–E sample glomerular-like, tubular, and tubulointerstitial regions, each explained by four NMF concepts (spatial maps, with concept labels and optimized explanation percentages). Glomerular-like structures, tubular fields, and neighboring interstitial regions depend on distinct but partially overlapping concept mixtures, so the contrast is not produced by a single kidney-wide component. Percentages are relative explanation weights, not molecular abundances or histopathological diagnoses.

#### Spatially complementary panel construction

This example shows how candidate pMiCS views are selected to increase local coverage of pathology-relevant structure. Each added visualization contributes information not already emphasized by the current panel.

**Figure S17:**
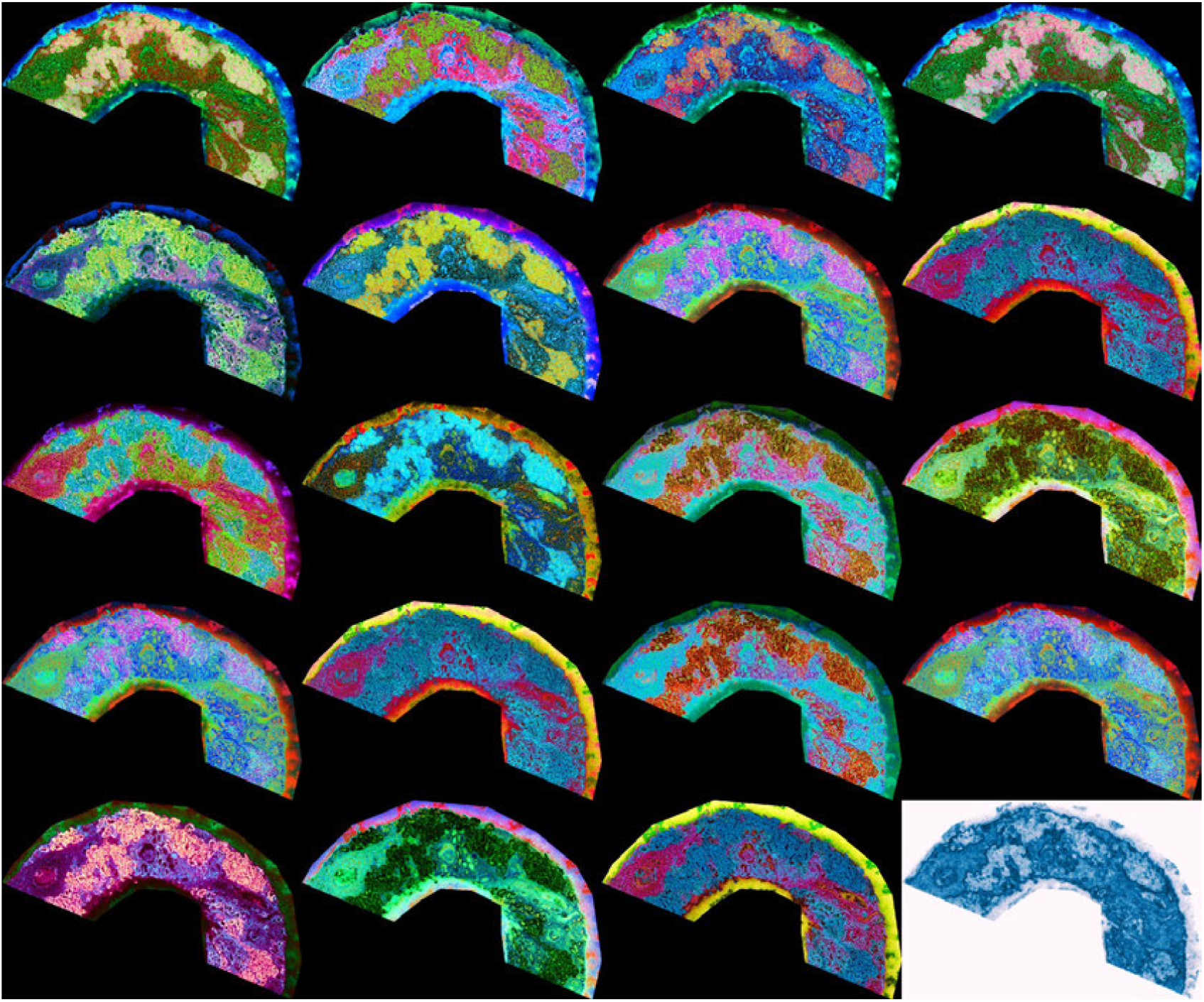
Spatially complementary pathology-oriented MSI panel construction 5 *µ*m spatial resolution. Candidate pMiCS visualizations were generated by hyperparameter tuning, including variation in the MiCS cluster combinations. From 300 candidates, 19 pMiCS views were selected together with the corresponding SoLaCE edge map. Local patch scores were aggregated using a running maximum, and each successive view was chosen to maximize the gain in patch coverage. The final white-background image shows the SoLaCE molecular edge map.

#### Rank-based panel construction

This example prioritizes visualizations with strong overall quality rather than local complementarity. It provides a compact alternative when the number of views must be limited for rapid review.

**Figure S18:**
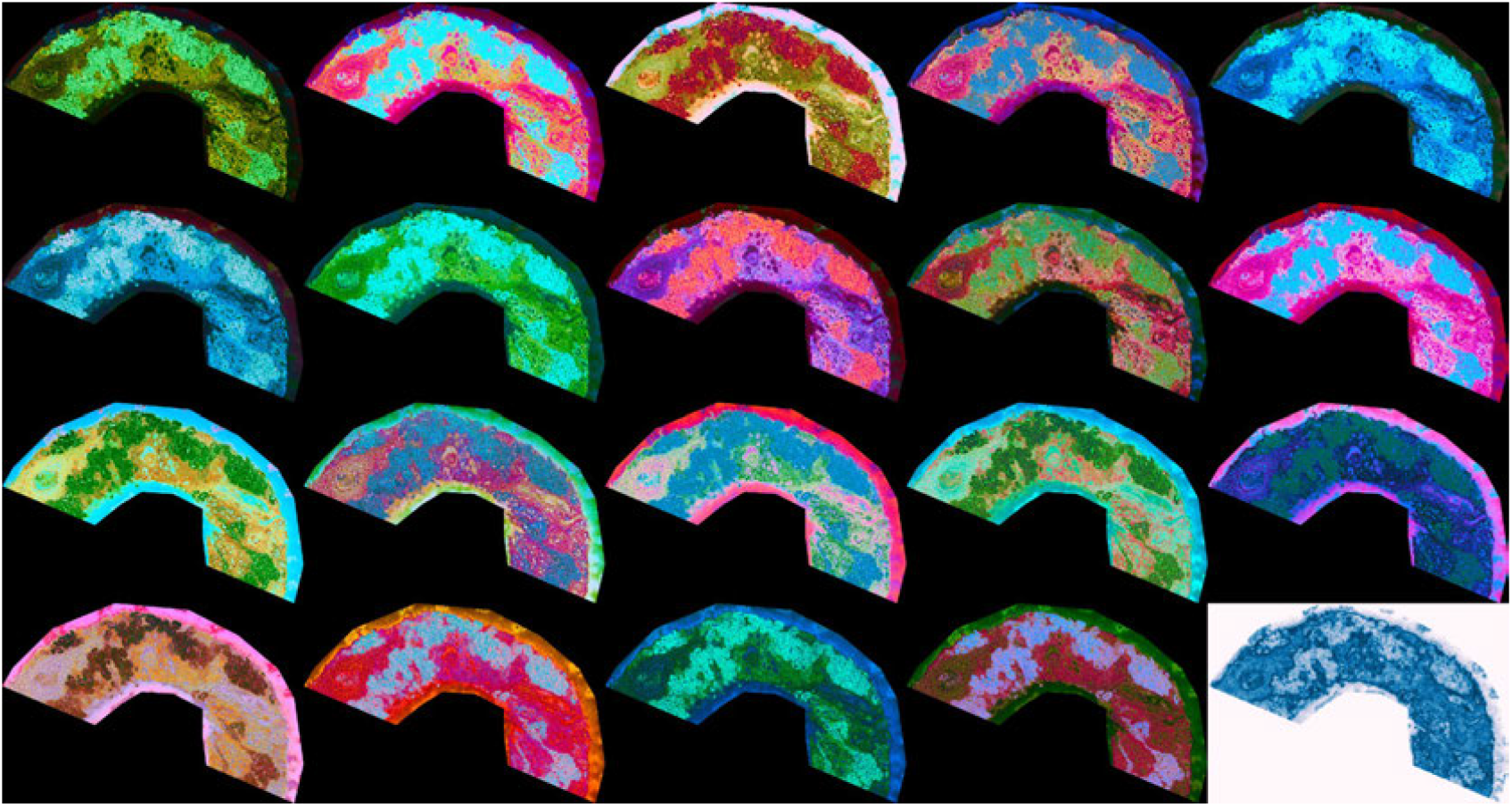
Rank-based pathology-oriented MSI panel construction 5 *µ*m spatial resolution. Candidate pMiCS visualizations were generated by sweeping MiCS hyperparameters, including the cluster combinations. Each candidate was evaluated using color-variety and edge-agreement criteria, and the highest-ranking views were selected to form a compact pathology-oriented panel.

#### Enlarged H&E image of adenocarcinoma metastasis ID9 (*35*)

**Figure S19:**
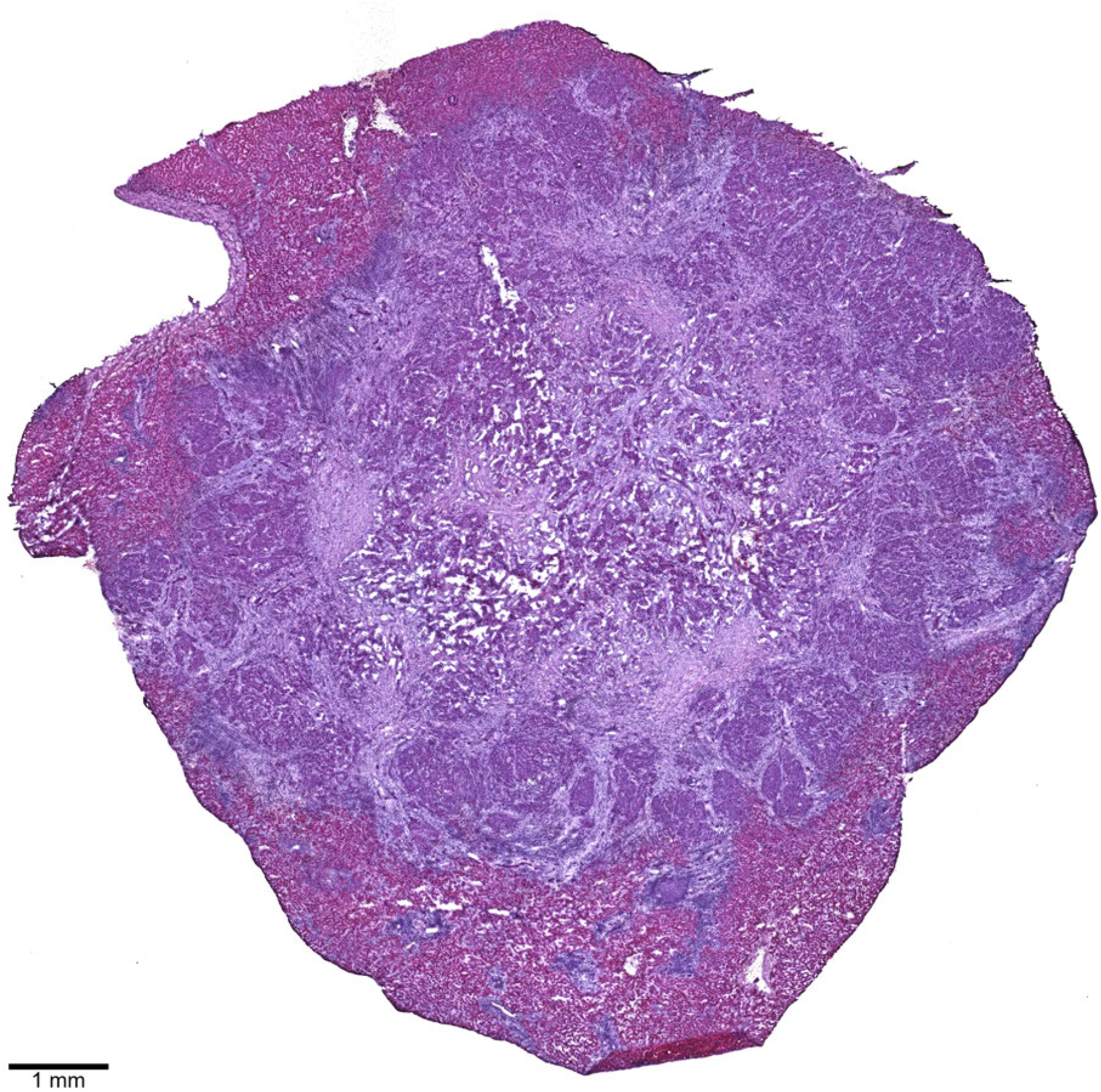
Enlarged H&E image of the colorectal adenocarcinoma metastasis (ID9). The section shows a central metastasis with heterogeneous structural compartments, including a peripheral ring of proliferative-appearing tumor nodules, a central matrix-rich region, and surrounding liver tissue containing finger-like extensions of the metastasis.

#### Adenocarcinoma metastasis visualizations - ID18a (*35*)

This independent section contains a large colorectal adenocarcinoma metastasis that occupies most of the sample. A small compartment of residual liver remains at the lower right and appears darker in the H&E image. Within the metastasis, the histology includes compact tumor regions and prominent gland-forming areas, providing a different tissue architecture from the rounded lesion shown in the main text. SoLaCE delineates the tumor–liver interface and a network of intratumoral transitions associated with the compact and glandular organization. Across the ranked pMiCS views, the residual liver repeatedly forms a molecularly distinct compartment, while the metastatic tissue is subdivided into several spectral phenotypes associated with solid, glandular, and intervening stromal-appearing regions. Positive- and negative-ion modes emphasize these compartments differently, indicating complementary molecular contrast rather than a uniform adenocarcinoma mass. This second example shows that rank-based VPP selection can be applied to metastases with different proportions of liver tissue and different growth patterns while identifying regions for targeted molecular comparison.

**Figure S20:**
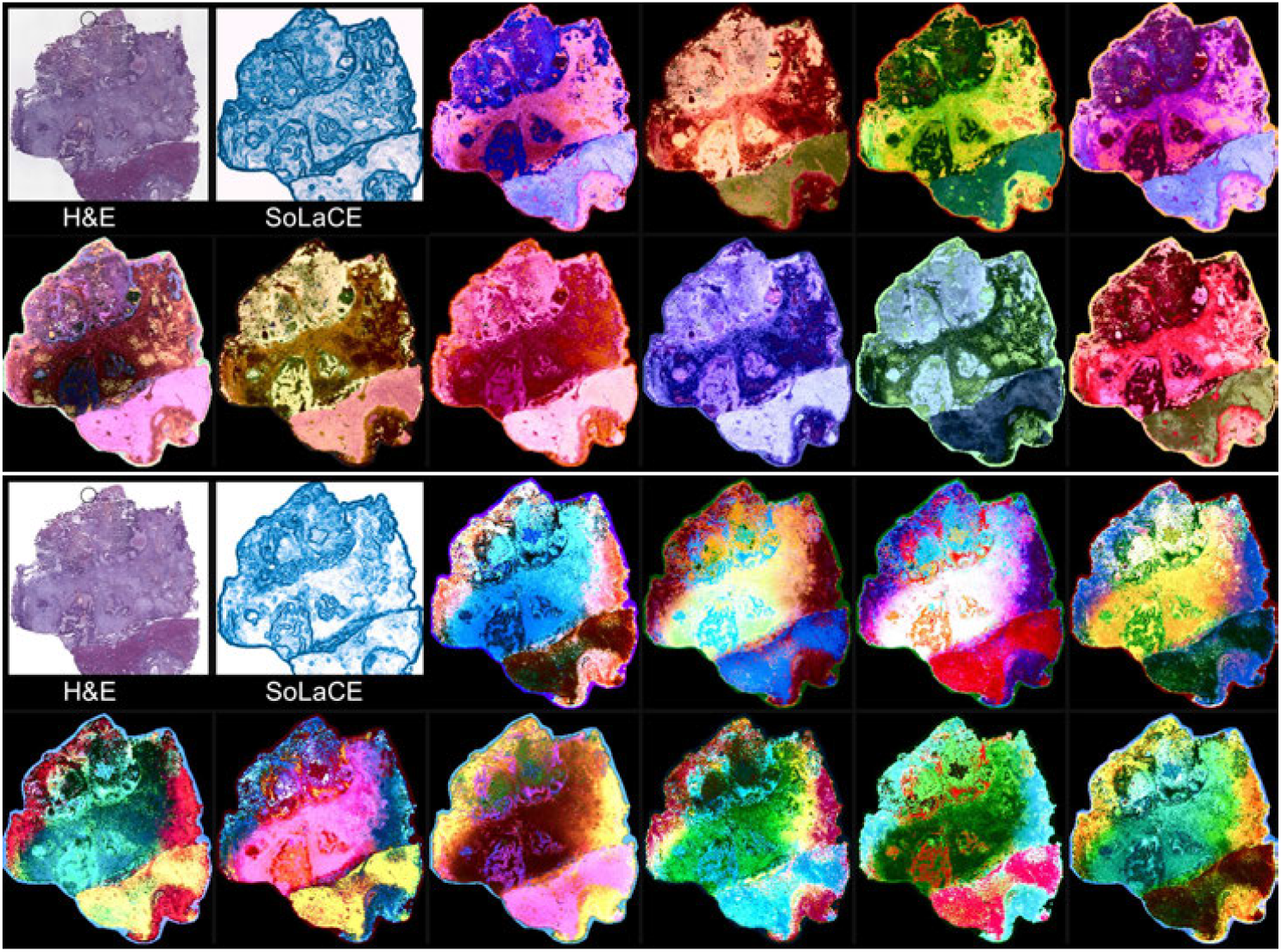
Ranked pathology-oriented VPP for a large colorectal adenocarcinoma liver metastasis (ID18a) with residual liver and heterogeneous growth patterns. Positive-ion-mode (top) and negativeion-mode (bottom) panels each show the H&E image, the corresponding SoLaCE molecular boundary map, and the highest-scoring pMiCS views. The darker compartment at the lower right of the H&E section represents residual liver, whereas most of the sample is occupied by metastatic adenocarcinoma. Within the tumor, compact regions coexist with gland-forming areas and intervening stromal-appearing tissue. SoLaCE highlights the interface between residual liver and metastasis and resolves additional boundaries within the tumor. Across the pMiCS views, the residual liver remains a recurrent molecular compartment, while compact and glandular tumor regions receive different spectral contrasts. The positive-and negative-ion modes emphasize these features differently, supporting complementary biochemical organization within the same histological lesion. Recurring patterns identify robust regions for comparison, whereas view-specific differences nominate compact, glandular, stromal-appearing, and residual-liver regions for targeted *m/z* mapping and molecular validation. Colors represent visualization-derived spectral phenotypes rather than definitive cell-type assignments.

## Supplementary Materials 2

### Overview

This file explains the mathematical quantities used in the manuscript for parametric MiCS+LMC visualization, concept-based interpretation, Soft Landmark Contrast Edges (SoLaCE), color-variety metrics, SpecEdge-Dice, an illustrative contrast-preservation example, hyperparameter tuning, and panel construction. It distinguishes model-output faithfulness from biological validation, and candidate-tuning scores from the pool-relative scores used to select complementary views.

### Parametric MiCS+LMC objective

The pMiCS model learns a neural mapping from each MSI spectrum to a three-dimensional visualization coordinate. The coordinate is subsequently mapped to CIELAB and converted to RGB for display; the neural output and displayed RGB color should therefore not be treated as interchangeable. The training objective combines a local, multiscale cluster-preservation term with a global landmark-distance preservation term:

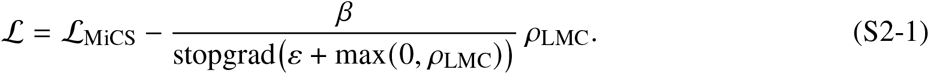

Here, *L*_MiCS_ is the Multi-resolution Cluster Supervision loss. It encourages the embedding to preserve cluster assignments generated at multiple resolutions in the original spectral space. The term *ρ*_LMC_ is the Landmark Mantel Correlation, which measures how well distances from pixels to a selected landmark set agree between the original high-dimensional MSI space and the low-dimensional embedding. Higher values of *ρ*_LMC_ indicate better agreement, so the term is subtracted from the loss.

In the displayed objective, *β* controls the scale of the LMC term and *ε* prevents division by zero. The max (0, *ρ*_LMC_) term keeps the normalization factor at least *ε;* when *ρ*_LMC_ < 0, the denominator uses *ε* while the signed numerator makes the LMC contribution increase the loss. The stopgrad operator treats the denominator as constant during backpropagation, so gradients flow through the numerator but not through the normalization factor.

In the implementation, the LMC contribution is scheduled during training, including the warm-up described in Materials and Methods. A frozen model can project unsampled pixels and other sections when preprocessing and spectral features are matched; this is not a guarantee of transfer across instruments or laboratories. Supplementary Figures S4 and S14 document sensitivity and out-of-sample projection, respectively.

Figure S2-1 gives a conceptual overview of pMiCS training.

**Figure S2-1:**
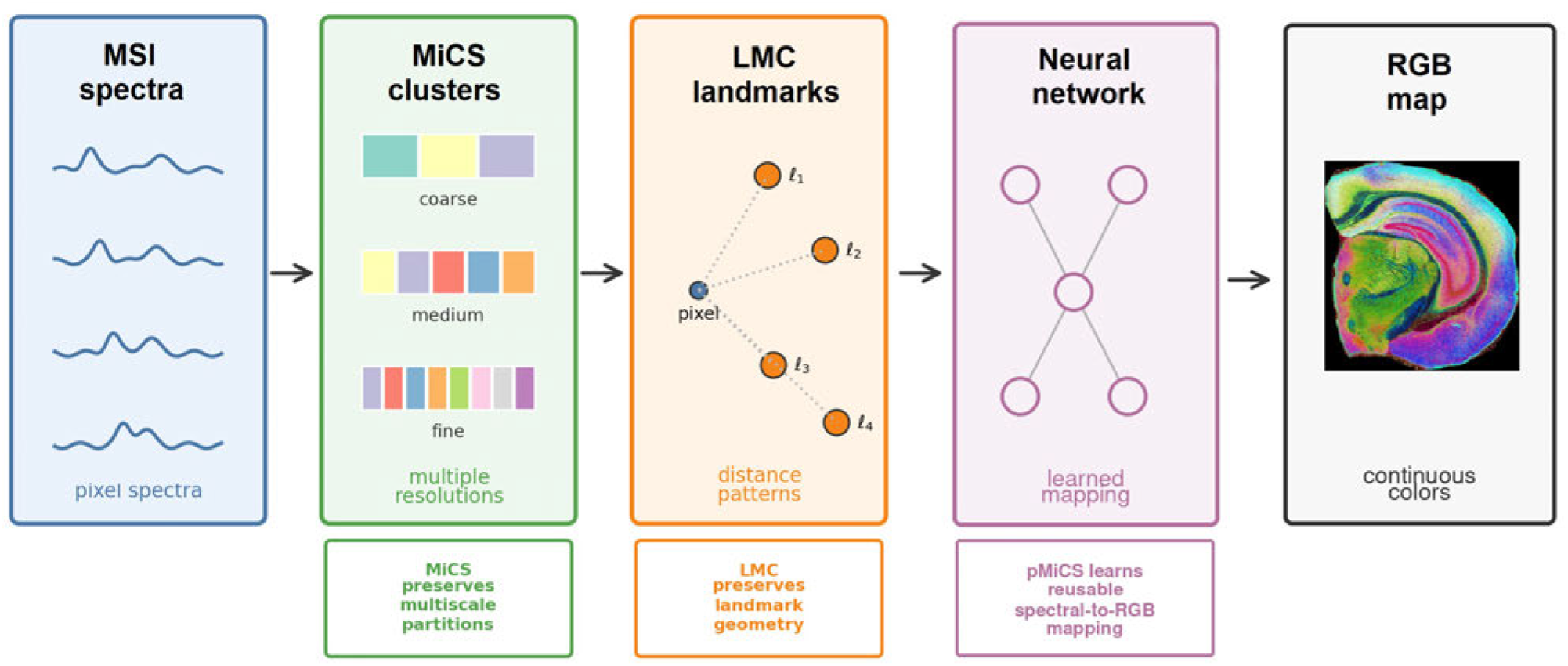
Conceptual overview of pMiCS training. Multi-resolution cluster supervision encourages preservation of local and regional tissue organization, while Landmark Mantel Correlation preserves global relationships to representative landmark spectra. A neural network distills these constraints into a reusable spectral-to-RGB mapping.

### Concept-based interpretation

The concept-based interpretation procedure seeks a sparse mixture of spectral concepts that approximately reproduces the output of a frozen pMiCS model. It distinguishes NMF spectral contributions from an optimized, model-output-equivalent explanation. Multiple mixtures can reproduce the same three-dimensional output, so the explanation is not a unique or causal attribution and does not establish molecular identity, cell type, or diagnostic significance.

The MSI data matrix is first approximated by nonnegative matrix factorization:

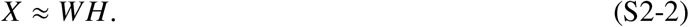

Here, 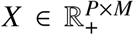 contains preprocessed spectra, 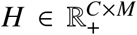 contains basis spectra, and 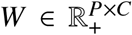 contains pixel activations. There are *P* pixels, *M* spectral bins, and *C* = 128 concepts. A concept is a weighted spectrum with a spatial activation map, not a single identified molecule. Its dominant *m/z* peaks can be inspected in Figure 5B and Table S1.

*F*or a pixel *p*, the trained pMiCS network maps the original spectrum *x*_*p*_ to a visualization coordinate:

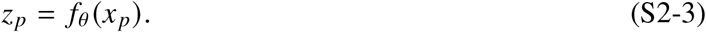

The vector *z*_*p*_ is the model coordinate used by the explanation loss; rendering it as RGB is a subsequent step. Sparse concept weights are obtained by applying a top-softmax operator to trainable logits *α*_*p*_:

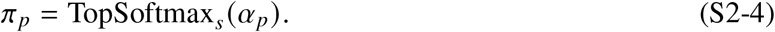

The operator applies softmax only to the *s* largest logits and assigns exactly zero weight to all other concepts. Thus *π*_*pk*_ ≥ 0 and ∑_*k*_ *π*_*pk*_ = 1, with at most *s* nonzero entries. This encourages concise explanations rather than diffuse mixtures.

Using the reconstruction notation of Materials and Methods, the sparse mixture forms an explanation spectrum from the concept basis:

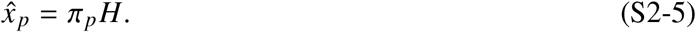

This reconstructed spectrum is then passed through the same trained pMiCS mapping:

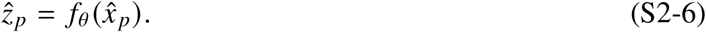

The basis scaling and input preprocessing must be held fixed between explanation fitting and evaluation. Closeness of 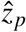 to *z*_*p*_ is a model-output criterion, not evidence that 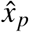 reconstructs every feature of *x*_*p*_. The loss is

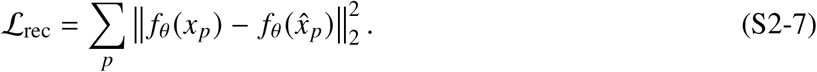

A sparsity-promoting entropy penalty is added to favor concise explanations:

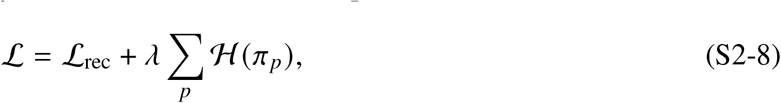

where the entropy of the concept mixture is

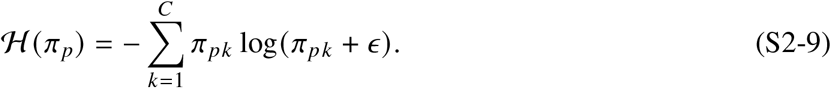

Here, *λ* controls the strength of the entropy penalty and *ϵ* prevents taking the logarithm of zero. Lower entropy means that the explanation uses fewer concepts more decisively.

To distinguish raw NMF coefficients from scale-adjusted spectral contributions, define *l*_*k*_ = ∥*h*_*k*_ ∥_1_ for each nonzero concept, 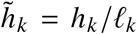, and 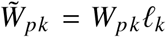 . Let 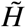 contain the rows 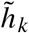; then 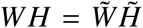, because rescaling a basis requires the inverse rescaling of its coefficients to preserve the spectrum. The normalized contribution distribution is

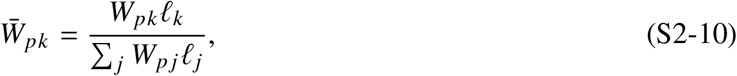

defined for pixels with positive total concept contribution. The manuscript compares this distribution with the fitted explanation weights using Jensen–Shannon divergence (*27*),

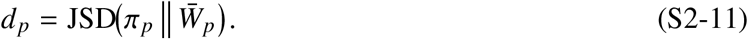

With 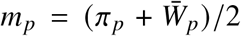, this is 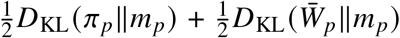 and lies in [0, 1] with base-two logarithms. A large *d*_*p*_ indicates disagreement between the fitted explanation weights and the scale-adjusted NMF contributions; it does not establish diagnostically useful contrast. Its interpretation is conditional on the basis scaling used to fit *π*_*p*_: normalized coefficients alone do not make an explanation invariant to basis rescaling. The distinction between the reconstruction basis and the contribution-normalization step must be retained when reproducing the analysis.

The concept-deletion test compares optimized-weight, abundance, and random rankings after identical deletion and re-optimization (Supplementary Figure S7). This assesses faithfulness to the frozen model under the tested perturbations, not biological truth or uniqueness. Spectral and rendered-color reconstruction fidelity are reported separately in Supplementary Figure S8. Supplementary Figures S9 and S10 compare *d*_*p*_ with abundance entropy; discrepancy and pixel mixedness are related but distinct quantities.

### Soft Landmark Contrast Edges

SoLaCE estimates molecular edges directly from high-dimensional MSI spectra rather than from a single ion image or an RGB visualization. Each pixel *i* has a spectrum *x*_*i*_. As specified in Materials and Methods, tissue spectra are TIC-normalized, projected onto 64 PCA components fitted on tissue pixels, and *l*_2_-normalized to obtain *z*_*i*_. A set of *L* = 1,000 landmarks *L*_land_ = {*l*_1_, …, *l*_*L*_*}* is sampled uniformly from valid tissue pixels. The default encoding retains *K* = 80 similarities with softmax temperature *τ* = 0.06; patch averaging uses a 3 × 3 neighborhood and texture suppression uses *α* = 0.22. Supplementary Figures S2 and S3 examine sensitivity and component ablation.

Figure S2-2 summarizes the SoLaCE workflow.

#### Cosine similarity to landmarks

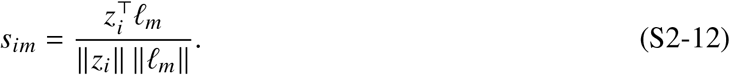

This equation computes the cosine similarity between pixel *i* and landmark *m*. Cosine similarity measures spectral direction rather than absolute magnitude, making it useful when the pattern of relative molecular abundances is more important than total intensity. A value near 1 indicates that the pixel and landmark have very similar projected spectral profiles; a value near 0 indicates weak similarity; negative values indicate opposite directions in the projected space.

**Figure S2-2:**
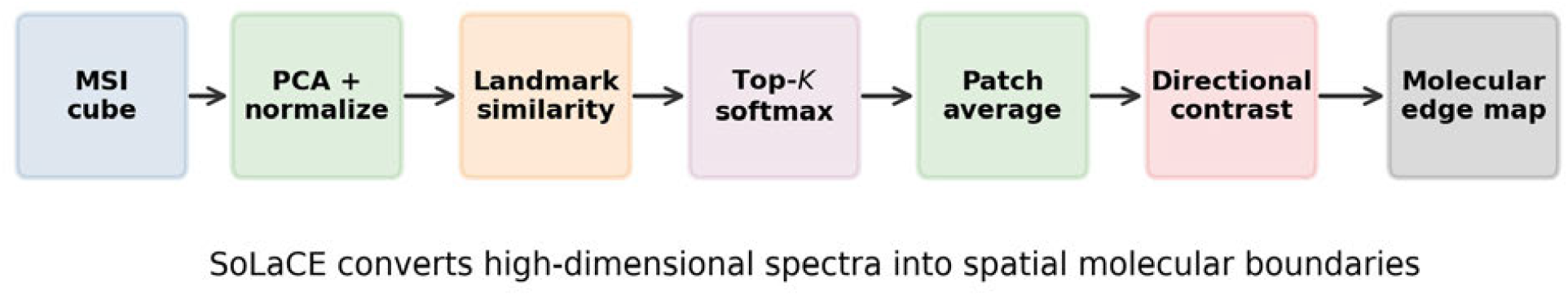
Workflow for Soft Landmark Contrast Edges. High-dimensional spectra are converted into sparse landmark-affinity signatures, smoothed locally, and compared across spatial directions to recover molecular boundaries directly from MSI data.

#### Top-*K* landmark neighborhood

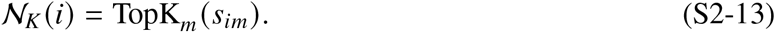

For each pixel, only the *K* most similar landmarks are retained. This creates a sparse representation that focuses on the most relevant reference spectra for the pixel and reduces noise from unrelated landmarks. In the manuscript, *K* = 80 is used.

#### Sparse soft landmark signature

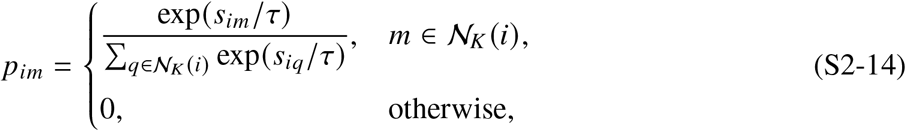

The top-*K* similarities are converted into a probability-like soft signature using a softmax. The value *p*_*im*_ is large when landmark *m* is among the closest spectral references for pixel *i*. Landmarks outside the top-*K* set receive zero weight. The temperature *τ* controls sharpness: smaller *τ* concentrates weight on the most similar landmarks, whereas larger *τ* spreads weight more evenly across the top-*K* landmarks.

This representation converts each pixel spectrum into a landmark-affinity profile. Instead of comparing thousands of mass channels directly, SoLaCE compares compact soft signatures that summarize the pixel’s spectral identity relative to representative landmark spectra.

Figure S2-3 illustrates the soft landmark signature.

#### Local patch averaging

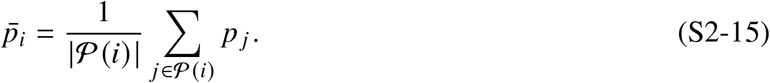

The vector *p*_*i*_ can vary from pixel to pixel because of noise, sampling variation, or small local heterogeneity. Therefore, signatures are averaged over a local patch *P(i)*, typically a 3 × 3 spatial neighborhood centered on pixel *i*. The averaged signature 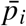 is a local molecular-context descriptor. This smoothing step improves robustness while still preserving boundaries between larger tissue states.

**Figure S2-3:**
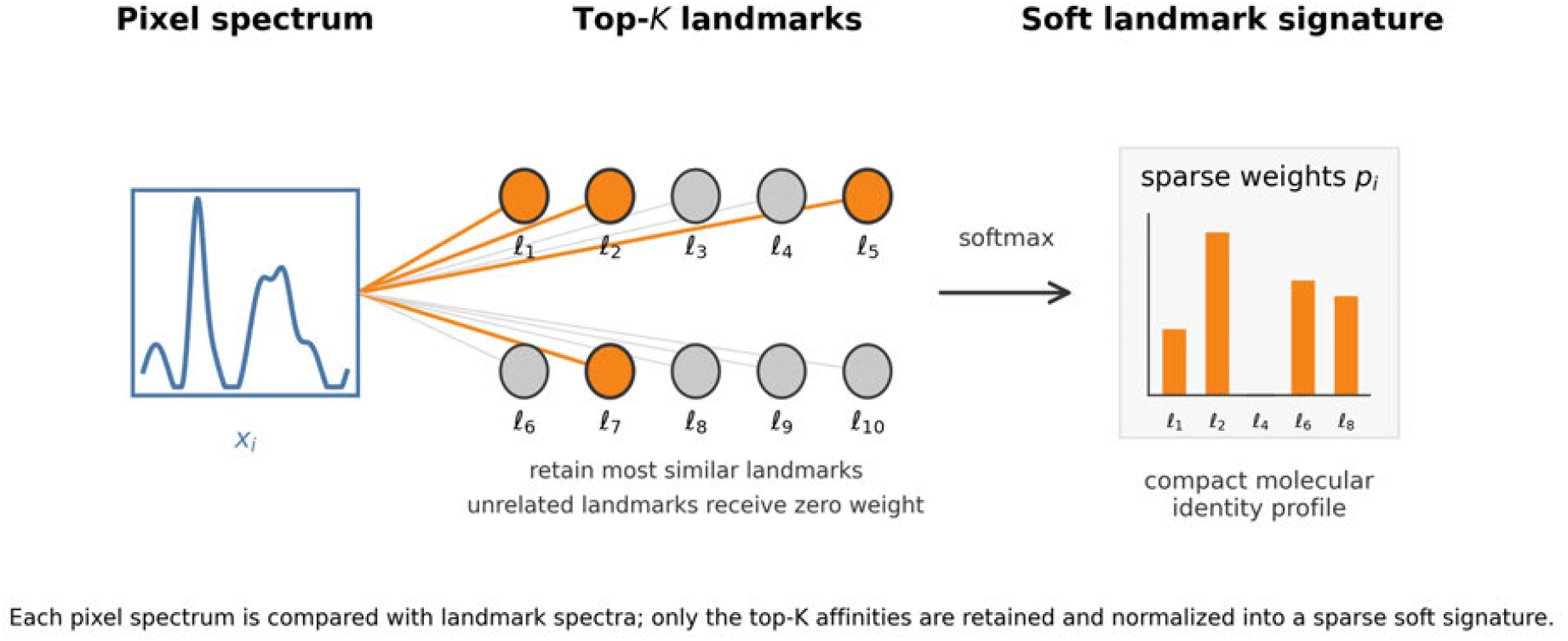
Soft landmark encoding of an MSI pixel. Each pixel spectrum is compared with landmark spectra, only the top-*K* most similar landmarks are retained, and the retained similarities are converted into a sparse softmax signature. This signature summarizes molecular identity while reducing noise from unrelated landmarks.

#### Across-boundary contrast

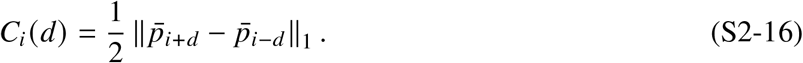

For a spatial direction *d*, this equation compares the averaged landmark signatures on opposite sides of pixel *i*. If the molecular context on one side differs strongly from the molecular context on the other side, the *L*_1_ distance is large and the pixel is likely to lie on a boundary. The factor 1/2 rescales the contrast because the *L*_1_ distance between two probability vectors can reach 2.

#### Local texture penalty

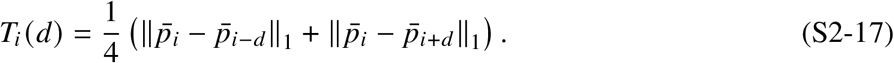

A high contrast score can arise from a true boundary, but it can also arise from local texture or noisy internal variation inside one tissue region. The texture term measures how different the center pixel is from each side individually. If the center differs strongly from both sides, the response may reflect local texture rather than a clean boundary between two coherent regions. The factor 1/4 keeps the penalty on a comparable scale to the across-boundary contrast.

#### Directional edge score

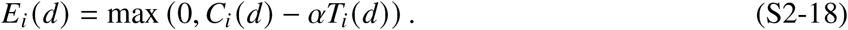

The directional edge score subtracts the texture penalty from the across-boundary contrast. The parameter *α* controls how strongly local texture is suppressed. The maximum with zero ensures that negative scores are clipped to zero, so only positive evidence for a boundary is retained.

#### Final SoLaCE edge strength

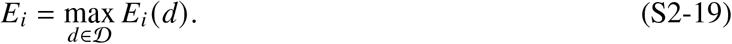

Edges are evaluated along the two undirected spatial axes *D* = {(1, 0, 0, 1)}, covering the four-connected neighbors on either side of each axis. The final edge strength at pixel *i* is the strongest directional response. This produces an edge map that highlights the most prominent molecular transition passing through each pixel. Percentile normalization is then used for visualization.

Figure S2-4 illustrates the across-boundary contrast and texture-suppression terms.

**Figure S2-4:**
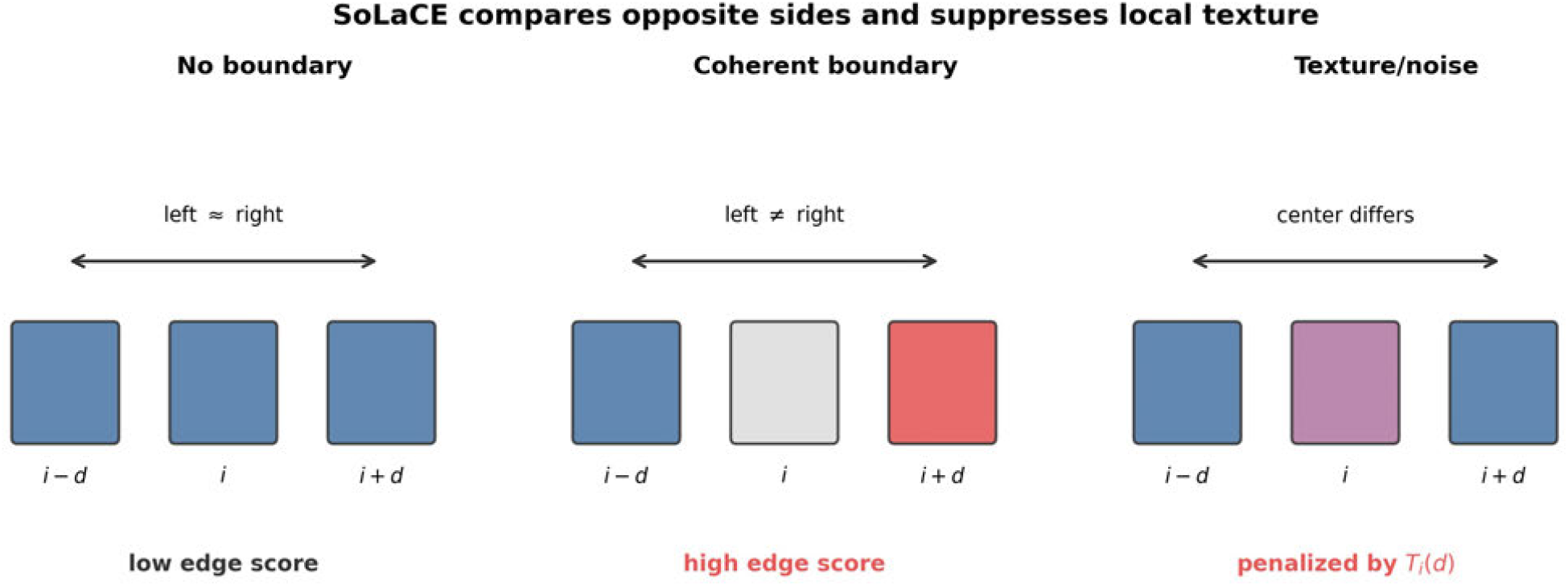
Directional SoLaCE edge scoring. Across-boundary contrast compares molecular signatures on opposite sides of a pixel, while the texture penalty reduces responses associated with local heterogeneity around the center pixel. Positive directional scores are retained and the maximum across directions forms the final edge map.

### Color-variety metrics

The manuscript uses two no-reference color-variety metrics, computed only over valid tissue pixels. They do not compare an RGB visualization to the original spectra or independently validated histology. They describe visual contrast and color diversity, not biological correctness. Empty tissue masks yield undefined scores.

#### Luminance RMS contrast

RGB values scaled to [0, 1] are first converted to Rec. 601 luminance,

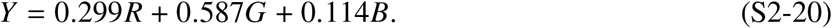

Luminance RMS contrast is the standard deviation of *Y* over valid tissue pixels. It measures global light–dark separation. Low values indicate a flat image with little intensity contrast; high values indicate stronger visual separation between bright and dark regions.

#### LAB chroma entropy

Quantized RGB values are converted to OpenCV-style *L*^∗^*a*^∗^*b*^∗^ color space. A 16 × 16 joint histogram is formed over the chromatic channels *a* and *b*, and Shannon entropy over occupied bins is computed as

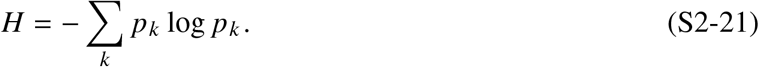

Here, *p*_*k*_ is the fraction of valid tissue pixels in chroma bin *k*. Higher entropy indicates use of more chromatic categories, but does not establish that those categories correspond to distinct biological states. Histogram binning and the logarithm base must be held fixed for comparisons.

## SpecEdge-Dice

SpecEdge-Dice compares molecular edges from the original MSI spectra with edges from a candidate RGB visualization:

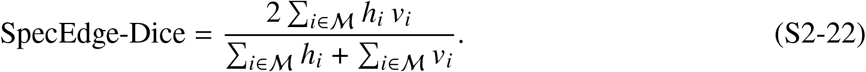

The set M is the valid tissue mask. The value *h*_*i*_ is the SoLaCE-derived reference edge strength and *v*_*i*_ is the visualization-derived edge strength at the same spatial pixel. Their pointwise product measures overlap, and the denominator is their total edge mass. For nonnegative edge strengths scaled to [0, 1] and a positive denominator, the score lies in [0, 1] . Unlike binary Dice, identical soft maps need not score 1: the numerator contains squared strengths when *h* = *v*. Consequently, the score depends on both edge location and edge strength.

The edge-extraction and normalization procedures must be fixed across candidates. Percentile normalization for display is not, by itself, a complete specification of metric preprocessing: the percentile bounds, clipping, and treatment of tissue-mask boundaries also determine the numerical score. If both maps have zero edge mass, the displayed formula is undefined; this case requires exclusion or an explicitly documented convention, rather than an assumed value. Empty tissue masks likewise do not provide a valid score.

SpecEdge-Dice measures agreement with the chosen spectral reference; it does not constitute validation against independent anatomy or molecular identity. Shifted, omitted, or extra visualization edges can reduce overlap, but the score alone cannot identify their biological cause. The equalization and *k*-means controls (Supplementary Figures S11 and S12) show why edge and color scores must be considered alongside geometric diagnostics and expert review. The tuning objective below uses only the edge/color subset; this practical choice does not make it a complete validation of visualization quality.

Figure S2-5 illustrates the overlap between visualization edges and SoLaCE boundaries used by SpecEdge-Dice.

## Illustrative contrast-preservation example

Consider scalar spectral values *x*_*i*_ sampled along a spatial tissue line, with a transition between two regions. Let the rendering compress contrast without rearranging pixels:

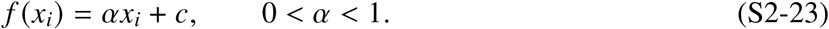

**Figure S2-5:**
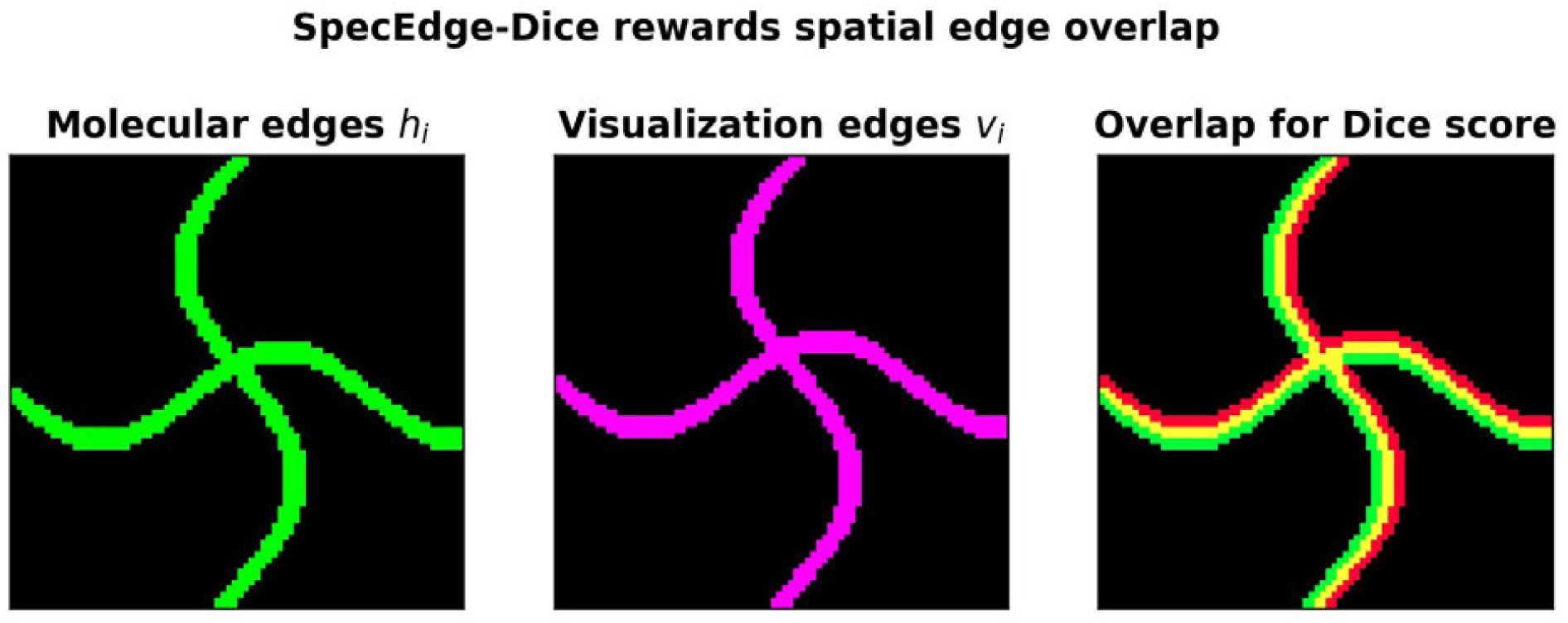
SpecEdge-Dice edge-overlap concept. SpecEdge-Dice evaluates whether RGB visualization edges coincide with molecular edges extracted from the original spectra. High scores occur when molecular and visualization-derived edge maps overlap; low scores occur when boundaries are shifted, blurred, omitted, or introduced artificially.

All pairwise distance rankings are preserved because |*f*(*x*_*i*_) – *f*(*x* _*j*_) = *α*|*x*_*i*_ − *x* _*j*_| . With consistent tie-breaking, nearest-neighbor rankings and trustworthiness are unchanged, although the rendered transition has less contrast.

For a conceptual gradient-based detector, choose a fixed common scale *A* > 0 so that

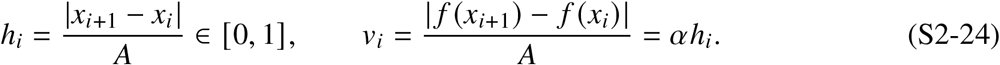

For a nonzero edge map, substitution into the SpecEdge-Dice formula gives

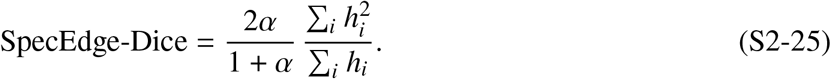

For a binary reference map, the second factor is 1, so the score decreases as contrast is compressed even though neighbor rankings do not change. For soft maps, the second factor additionally reflects edge strength. This is an illustrative calculation, not a rerun or validation of the benchmark’s SoLaCE pipeline.

Independent normalization of each edge map can remove the uniform scaling effect; no universal decrease in the implemented metric is asserted. The example isolates why neighborhood rankings do not guarantee displayed contrast and why edge normalization must be reported. The schematic below illustrates the broader distinction conceptually, rather than plotting this calculation.

## Hyperparameter tuning objective

Candidate pMiCS visualizations are scored using three metrics: SpecEdge-Dice, luminance RMS contrast, and LAB chroma entropy. Because these metrics have different units and ranges, each metric is clipped and scaled to the interval [0, 1]:

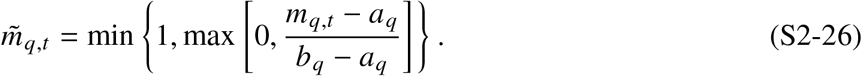

**Figure S2-6:**
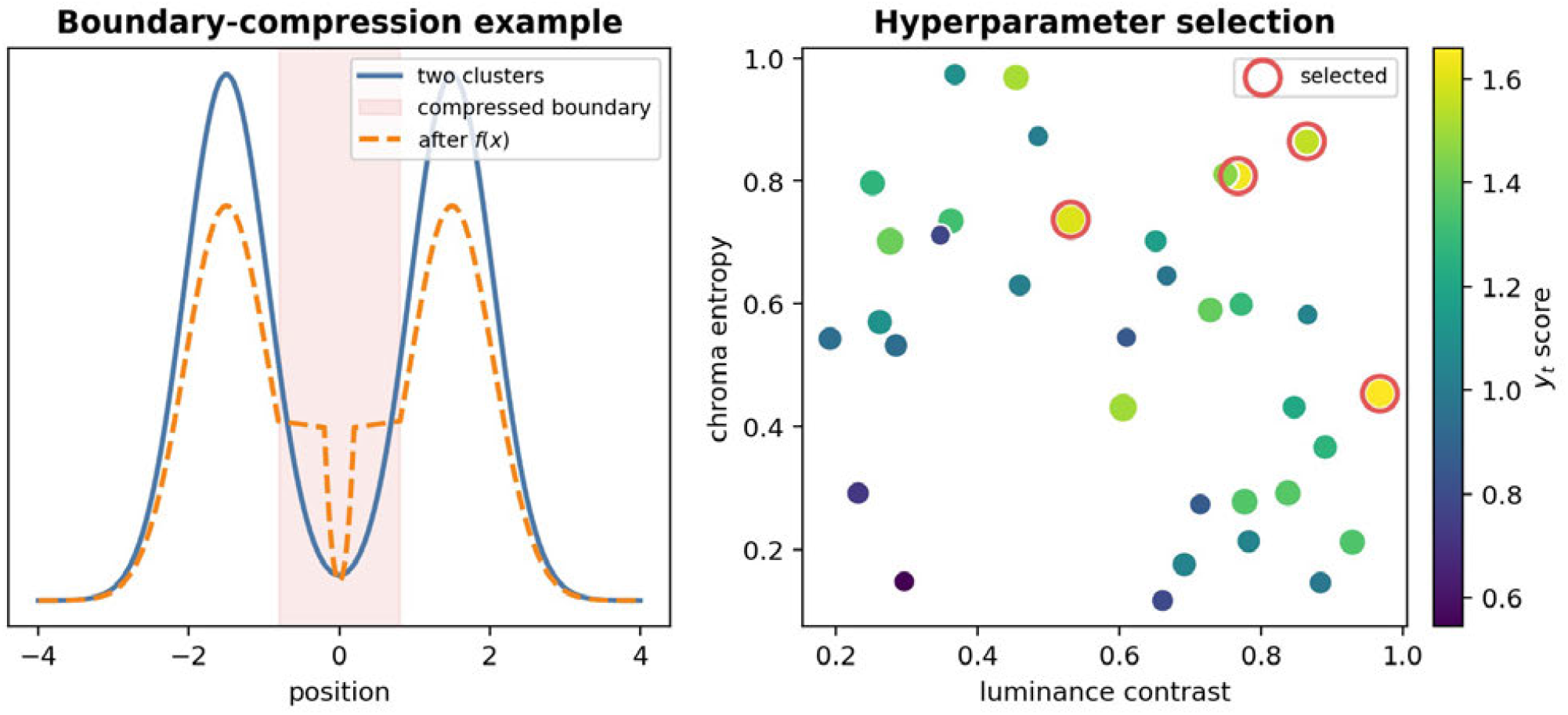
Conceptual illustration of boundary fidelity and visualization selection. Left: preserving neighborhoods alone does not guarantee a visually informative spatial transition; the sketch is conceptual, not a quantitative result of the affine contrast example. Right: candidate visualizations can be represented in metric space and selected according to the combined tuning objective *y*_*t*_. Neither panel establishes biological or diagnostic validity.

Here, *m*_*q,t*_ is the raw score of metric *q* for trial *t*, and (*a*_*q*_, *b*_*q*_) are user-specified lower and upper bounds for that metric. Values below *a*_*q*_ become 0, values above *b*_*q*_ become 1, and values between the bounds are linearly scaled.

The final scalar score for trial *t* is

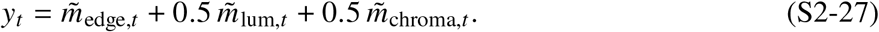

This objective gives spectral-reference edge agreement twice the weight of each color-variety metric: weights are 1, 0.5, and 0.5, respectively, and *y*_*t*_ ∈ [0, 2] . The bounds must satisfy *b*_*q*_ > *a*_*q*_ and remain fixed across compared trials. The score is a practical selection criterion, not a biological or diagnostic validity measure; geometric preservation is not explicitly optimized by this scalar score.

As described in Materials and Methods, candidate search first uses coordinate one-factor-at-a-time exploration and then stochastic hill climbing over neighboring configurations. Duplicate configurations are skipped and a fixed trial budget limits the search. The alternative `rank composite` mode is used only when explicitly selected; it is not identical to the bounded weighted score above.

The right panel of Figure S2-6 illustrates this selection concept: candidate visualizations are arranged by color-variety metrics, colored by the combined score, and selected according to the objective.

## Panel construction and complementarity

Candidate tuning scores a configuration individually. Panel construction instead selects a set of views from a fixed candidate pool, using pool-relative ranks and, for the spatial strategy, marginal coverage gains. These are distinct operations even though they use the same three metrics and weights.

### Local and whole-image quality

As specified in Materials and Methods, the three metrics are computed on overlapping patches of size 48 and stride 16. At each valid tissue location, candidate metric values are rank-normalized across the pool to [0, 1]. For candidate *i*, the resulting local quality is

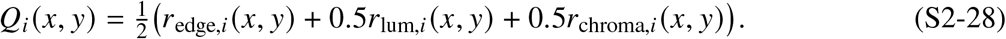

The same construction applied to whole-image metric ranks gives

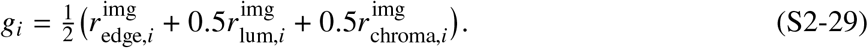

Both scores lie in [0, 1] and are relative to the candidate pool. Changing that pool can change the ranks and scores; they are not absolute measures of tissue quality and are not the bounded trial score *y*_*t*_.

### Selection strategies

All strategies start with the highest-*g*_*i*_ view. Greedy selection then adds views in descending *g*_*i*_ order. Spatial selection adds the view with the largest increase in the running local-quality envelope,

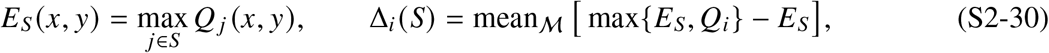

where *S* is the set already selected. A view is useful under this rule when it improves tissue locations that the current panel represents weakly.

Coreset selection uses DINOv2 image embeddings *z*_*i*_ and repeatedly adds the view farthest from the selected set,

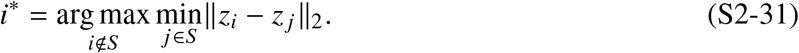

The random baseline draws the remaining views uniformly. Embedding diversity need not imply useful molecular contrast: a near-uniform rendering may be distant from other views without adding tissue detail.

### Gallery evaluation

The three measures reported in Figure 8 and Supplementary Figure S6 are the mean local envelope (higher is better), the embedding covering radius (lower is better), and the weakest included whole-image score (higher is better):

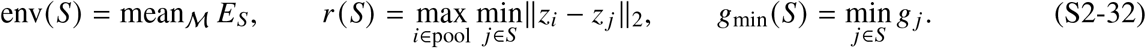

Each measure matches one strategy’s objective by construction: spatial selection targets the envelope, coreset selection targets embedding coverage, and greedy selection prioritizes individual view quality. They are complementary descriptions, not independent biological validation. Example spatial and greedy panels are shown in Supplementary Figures S17 and S18.

## Summary

The equations link five operations: learning a reusable pMiCS embedding, fitting sparse model-output-equivalent concept explanations, extracting spectral-reference edges with SoLaCE, scoring candidate visualizations, and selecting complementary panels. These operations support exploratory pathology-oriented review; model faithfulness, spectral association, and confirmed biological identity remain distinct levels of evidence. Transfer to other sections requires matched preprocessing and features, and clinical or cross-laboratory deployment requires validation beyond the analyses presented here.

## References and Notes

1. J. Gildenblat, J. Stamnas, J. Pahnke, Mass spectrometry imaging-based explainable machine learning reveals the biochemical landscapes of the mouse brain. Free Neuropathology 7, 9 (2026), doi:10.17879/freeneuropathology-2026-9413, https://doi.org/10.17879/freeneuropathology-2026-9413.

2. J. Gildenblat, J. Pahnke, Truthful visualizations for mass spectrometry imaging enable high spatial resolution interactive m/z mapping and exploration. Science Advances (2026), doi:10.1126/sciadv.aed3650.

3. W. Gardner, et al., Self-Organizing Map and Relational Perspective Mapping for the Accurate Visualization of High-Dimensional Hyperspectral Data. Anal Chem 92 (15), 10450–10459 (2020), doi: 10.1021/acs.analchem.0c00986, https://www.ncbi.nlm.nih.gov/pubmed/32614172.

4. J. A. Lee, M. Verleysen, Quality assessment of dimensionality reduction: Rank-based criteria. Neurocomputing 72 (7-9), 1431–1443 (2009), doi:10.1016/j.neucom.2008.06.011.

5. T. Sainburg, L. McInnes, T. Q. Gentner, Parametric UMAP embeddings for representation and semi-supervised learning. arXiv.org (2021), doi:10.48550/arXiv.2009.12981, https://arxiv.org/abs/2009.12981.

6. T. Alexandrov, et al., Spatial Segmentation of Imaging Mass Spectrometry Data with Edge-Preserving Image Denoising and Clustering. Journal of Proteome Research 9 (12), 6535–6546 (2010), doi: 10.1021/pr100734z.

7. T. Alexandrov, J. H. Kobarg, Efficient Spatial Segmentation of Large Imaging Mass Spectrometry Datasets with Spatially Aware Clustering. Bioinformatics 27 (13), i230–i238 (2011), doi:10.1093/bioinformatics/btr246.

8. H. Hu, R. Yin, H. M. Brown, J. Laskin, Spatial Segmentation of Mass Spectrometry Imaging Data by Combining Multivariate Clustering and Univariate Thresholding. Anal Chem 93 (7), 3477–3485 (2021), doi:10.1021/acs.analchem.0c04798, https://www.ncbi.nlm.nih.gov/pubmed/33570915.

9. K. A. Bemis, M. C. Foll, D. Guo, S. S. Lakkimsetty, O. Vitek, Cardinal v.3: a versatile open-source software for mass spectrometry imaging analysis. Nat Methods 20 (12), 1883–1886 (2023), doi:10.1038/s41592-023-02070-z, https://www.ncbi.nlm.nih.gov/pubmed/37996752.

10. Bruker Daltonics, SCiLS Lab: mass spectrometry imaging analysis software, Bruker Daltonics GmbH, Bremen, Germany (2024), rRID:SCR 014426.

11. L. van der Maaten, G. Hinton, Visualizing Data using t-SNE. Journal of Machine Learning Research 9 (86), 2579–2605 (2008).

12. L. McInnes, J. Healy, J. Melville, UMAP: Uniform Manifold Approximation and Projection for Dimension Reduction. arXiv preprint arXiv:1802.03426 (2018), doi:10.48550/arXiv.1802.03426, https://arxiv.org/abs/1802.03426.

13. W. M. Abdelmoula, et al., Data-driven identification of prognostic tumor subpopulations using spatially mapped t-SNE of mass spectrometry imaging data. Proceedings of the National Academy of Sciences 113 (43), 12244–12249 (2016), doi:10.1073/pnas.1510227113.

14. W. M. Abdelmoula, et al., Peak learning of mass spectrometry imaging data using artificial neural networks. Nature Communications 12, 5544 (2021), doi:10.1038/s41467-021-25744-8.

15. T. Alexandrov, et al., METASPACE: A community-populated knowledge base of spatial metabolomes in health and disease. bioRxiv (2019), doi:10.1101/539478, https://www.biorxiv.org/content/early/2019/02/13/539478.

16. A. R. Buchberger, K. DeLaney, J. Johnson, L. Li, Mass Spectrometry Imaging: A Review of Emerging Advancements and Future Insights. Anal Chem 90 (1), 240–265 (2018), doi:10.1021/acs.analchem.7b04733, https://www.ncbi.nlm.nih.gov/pubmed/29155564.

17. T. Alexandrov, Spatial Metabolomics and Imaging Mass Spectrometry in the Age of Artificial Intelligence. Annu Rev Biomed Data Sci 3, 61–87 (2020), doi:10.1146/annurev-biodatasci-011420-031537, https://www.ncbi.nlm.nih.gov/pubmed/34056560.

18. N. Verbeeck, R. M. Caprioli, R. Van de Plas, Unsupervised Machine Learning for Exploratory Data Analysis in Imaging Mass Spectrometry. Mass Spectrometry Reviews 39 (3), 245–291 (2020), doi: 10.1002/mas.21602.

19. J. Gildenblat, J. Pahnke, Dimensionality reduction with strong global structure preservation. Pattern Analysis and Applications 29, 38 (2026), doi:10.1007/s10044-025-01585-9, https://doi.org/10.1007/s10044-025-01585-9.

20. S. M. Sunkin, et al., Allen Brain Atlas: an integrated spatio-temporal portal for exploring the central nervous system. Nucleic Acids Res 41 (Database issue), D996–D1008 (2013), doi:10.1093/nar/gks1042, https://www.ncbi.nlm.nih.gov/pubmed/23193282.

21. G. E. Hinton, R. R. Salakhutdinov, Reducing the Dimensionality of Data with Neural Networks. Science 313 (5786), 504–507 (2006), doi:10.1126/science.1127647.

22. L. van der Maaten, Learning a Parametric Embedding by Preserving Local Structure, in Proceedings of the Twelfth International Conference on Artificial Intelligence and Statistics (AISTATS), vol. 5 of Proceedings of Machine Learning Research (2009), pp. 384–391.

23. I. Sobel, G. Feldman, A 3×3 Isotropic Gradient Operator for Image Processing, Presented at the Stanford Artificial Intelligence Project (SAIL) (1968), commonly referred to as the Sobel operator.

24. T. Schramm, et al., imzML – A Common Data Format for the Flexible Exchange and Processing of Mass Spectrometry Imaging Data. Journal of Proteomics 75 (16), 5106–5110 (2012), doi:10.1016/j.jprot.2012.07.026, https://pubmed.ncbi.nlm.nih.gov/22842151/.

25. D. D. Lee, H. S. Seung, Learning the parts of objects by non-negative matrix factorization. Nature 401, 788–791 (1999), doi:10.1038/44565, https://www.nature.com/articles/44565.

26. J. Leuschner, et al., Supervised Non-Negative Matrix Factorization Methods for MALDI Imaging Applications. Bioinformatics 35 (11), 1940–1947 (2019), doi:10.1093/bioinformatics/bty909.

27. J. Lin, Divergence measures based on the Shannon entropy. IEEE Transactions on Information Theory 37 (1), 145–151 (1991), doi:10.1109/18.61115.

28. S. Kullback, R. A. Leibler, On Information and Sufficiency. The Annals of Mathematical Statistics 22 (1), 79–86 (1951), doi:10.1214/aoms/1177729694.

29. V. Petsiuk, A. Das, K. Saenko, RISE: Randomized Input Sampling for Explanation of Black-box Models, in British Machine Vision Conference (BMVC) (2018), doi:10.48550/arXiv.1806.07421.

30. S. Hooker, D. Erhan, P.-J. Kindermans, B. Kim, A Benchmark for Interpretability Methods in Deep Neural Networks, in Advances in Neural Information Processing Systems (NeurIPS) (2019), doi: 10.48550/arXiv.1806.10758.

31. The VGG Image Annotator (VIA) (ACM) (2019), doi:10.1145/3343031.3350535, https://dl.acm.org/doi/10.1145/3343031.3350535.

32. S. J. Russell, P. Norvig, Artificial Intelligence: A Modern Approach (Pearson, Hoboken, NJ), 4 ed. (2020).

33. M. Oquab, et al., DINOv2: Learning Robust Visual Features without Supervision. Transactions on Machine Learning Research (TMLR) (2024), doi:10.48550/arXiv.2304.07193, https://arxiv.org/abs/2304.07193.

34. O. Sener, S. Savarese, Active Learning for Convolutional Neural Networks: A Core-Set Approach, in International Conference on Learning Representations (ICLR) (2018), doi:10.48550/arXiv.1708.00489, https://arxiv.org/abs/1708.00489.

35. M. F. Rittel, et al., Cohort-Scale Spatial Autocorrelation for Tumor Prediction in Mid-Infrared Pathology and Spatial Biomarker Discovery Using MALDI Imaging Lipidomics. Advanced Science 13 (24), e16847 (2026), doi:10.1002/advs.202516847, https://advanced.onlinelibrary.wiley.com/doi/10.1002/advs.202516847.

36. M. Schwaiger-Haber, et al., Using mass spectrometry imaging to map fluxes quantitatively in the tumor ecosystem. Nat Commun 14 (1), 2876 (2023), doi:10.1038/s41467-023-38403-x, https://www.ncbi.nlm.nih.gov/pubmed/37208361.

37. H. Uchino, H. Tsugawa, M. Arita, SMASH Imaging: A Serial Matrix-Assisted Laser Desorption/Ionization Mass Spectrometry Strategy for High-Resolution Imaging Facilitates Dual-Polarity and MS2 Spatial Lipidomics on a Single Tissue Section. Analytical Chemistry 98 (13), 9540–9554 (2026), doi:10.1021/acs.analchem.5c03738.

38. M. D. Ernst, Permutation methods: A basis for exact inference. Statistical Science 19 (4), 676–685 (2004), doi:10.1214/088342304000000396.

39. S. Holm, A simple sequentially rejective multiple test procedure. Scandinavian Journal of Statistics 6 (2), 65–70 (1979).

40. F. Wilcoxon, Individual comparisons by ranking methods. Biometrics Bulletin 1 (6), 80–83 (1945), doi:10.2307/3001968.

